# Convergent Inhibitory Cortical Circuit Disruption Drives Genetically Distinct Absence Seizures

**DOI:** 10.64898/2026.07.10.737818

**Authors:** Siyuan Song, Shu-Ning Natalie Lau, Doan Tran, Cecilia Ljungberg, Jianrong Tang, Qinglong Miao, Atul Maheshwari, Jeffrey L. Noebels, Xiaolong Jiang

## Abstract

Childhood absence epilepsy is one of the most common pediatric epilepsies, characterized by brief involuntary lapse in consciousness and generalized spike-wave discharges (SWDs) on EEG. Despite a growing list of causative genes, the core cortical circuit abnormalities driving this highly stereotyped SWD phenotype remain elusive. Using two well-established absence epilepsy mouse models carrying point mutations in distinct genes altering pre and postsynaptic transmission, stargazer (*Cacng2*) and tottering (*Cacna1a*), we show that a specific translaminar inhibitory microcircuit defined by layer 6 corticothalamic neurons (L6 CT) and a novel Tac1^+^ *Pvalb* interneuron subtype is disrupted in both models. This early functional defect was followed by a secondary, marked dysfunction of deep-layer somatostatin-expressing (*Sst*) interneurons that coincides with the postnatal developmental onset of seizures. Crucially, we established the serial causality of these cascading defects by demonstrating that chemogenetic disruption of the L6 CT-Tac1^+^ *Pvalb* microcircuit alone in wild-type mice is sufficient to induce both the secondary *Sst* interneuron dysfunction and contemporaneous emergence of SWDs. Together, these findings reveal a sequential translaminar synaptic dysfunction or ‘domino effect’ wherein a primary cell-autonomous defect triggers broader circuit adaptations permitting spike-wave hypersynchrony. Our results thus identify a convergent, causative cortical circuit pathology across two genetically heterogeneous absence epilepsy models, highlighting a common functional target for intervention.

## Introduction

Absence epilepsy is the most common seizure disorder in children and is characterized by distinctive 3 Hz bilaterally synchronous spike-wave discharges accompanied by a brief loss of consciousness and voluntary movement^1^. Despite a high remission rate in adolescence, a majority of children with spike-wave epilepsy experience persistent cognitive comorbidities, including deficits in attention, sensory processing, memory, and mood impairment^1–4^. Moreover, approximately 30% of patients are refractory to monotherapy with current front-line medication^5^. The high incidence of pharmacoresistance and the persistence of comorbidities even after full seizure control^6,7^ highlight the need of a holistic understanding of the underlying circuit defects to facilitate development of novel targeted and effective treatment.

Childhood absence epilepsy (CAE) is a genetic rather than acquired disorder, with over 20 distinct monogenic mutations identified^8,9^ alongside numerous unresolved polygenic or oligogenic etiologies^10^. While the hallmark spike-wave discharges (SWDs) of absence seizures reflect aberrant synchronization in the cortico-thalamo-cortical (CTC) network^1,11,12^, it remains enigmatic how mutations in a diverse array of genes functionally converge to produce this highly stereotyped EEG and behavioral pattern. Historically, SWD oscillations has been attributed to intrathalamic circuit disturbances^9,13–16^. However, compelling evidence from various pharmacological and genetic models^17–20^, alongside clinical observations^21–24^, suggest that these seizures depend on a lower threshold for pathological oscillations within a focal onset zone in somatosensory cortex (S1) and mesial frontal cortex. However, the cellular basis for cortical dysfunction has not been widely explored, and as a result, the cellular, synaptic, and circuit abnormalities that drive spike-wave epileptogenesis, as well as how genetic mutations precipitate these abnormalities, remain a critical knowledge gap. Some evidence points toward altered excitability, synaptic inhibition, and connectivity within S1, particularly involving parvalbumin-expressing interneurons and disrupted feedforward inhibition^25–29^, yet it is unclear whether these alterations represent an early primary cause of epilepsy or secondary consequences of repeated seizures. Differentiating primary derangements from downstream secondary remodeling by teasing out causative versus non-causative defects poses a significant challenge in a disorder where the neurodevelopmental defects evolve alongside aberrant brain activity^30–32^. Ultimately, since seizures are network-level phenomena linked to circuit plasticity, bridging the gap between inherited molecular disruption and epilepsy in the intact brain requires serial dynamic, circuit-level analyses.

In this study, we performed a systematic, longitudinal circuit analysis across developmental stages of two well-characterized monogenic mouse models of absence seizures, stargazer (*stg*) and tottering (*tg*) mice, to identify key cortical cells and microcircuitry essential for spike-wave (SW) seizure initiation. Despite exhibiting the same hallmark hypersychronous EEG discharges, the two models harbor unrelated primary genetic dysfunctions; *stg* mice carry a spontaneous mutation in *Cacng2* encoding the AMPA receptor auxiliary protein stargazin/TARPγ2^33,34^, whereas *tg* mice carry a spontaneous mutation in *Cacna1a* encoding the pore-forming α1 subunit of P/Q-type calcium channels^35^. Leveraging the restricted expression of *Cacng2* in cortical interneurons along with single-cell transcriptomics, multi-patch recording, optogenetics, and chemogenetics, we uncovered a critical translaminar inhibitory microcircuit comprising layer 6 corticothalamic neurons (L6 CT) and a novel subtype of parvalbumin (*Pvalb*) interneurons implicated in sensory gain control. In both seizure models, early hereditary genetic lesions impair this intracortical microcircuitry in distinct ways, yet converge on the same functional deficit before seizure onset. In addition, we observed a subsequent sequential synaptic dysfunction of deep-layer somatostatin-expressing (*Sst*) interneurons that coincides with the developmental onset of seizure, suggesting that additional circuit adaptations contribute to SW epileptogenesis in a domino-like fashion. Using mouse genetics and chemogenetics, we further established that disrupting this L6 CT-*Pvalb* cortical microcircuit is sufficient to generate secondary *Sst* interneuron changes and SW discharges in wild-type mice. Our extensive developmental mapping of seizure-prone cortical circuits in two distinct genetic models thus delineates stereotypical, causative cortical circuit abnormalities driven by genetic lesions and circuit remodeling. These findings refine a minimal intracortical circuit pathology shared by two distinct monogenic models, which if replicated in additional CAE models, highlight a common inhibitory translaminar microcircuit as a promising target for the treatment of absence epilepsy and its comorbidities.

## Results

### Stargazin *(Cacng2)* expression defines a novel translaminar cortical inhibitory microcircuit for sensory gain control

To understand how mutations of *Cacng2* give rise to spike-wave absence seizure phenotypes in *stg* mice, we began by examining the expression of *Cacng2* in the mouse somatosensory cortex (S1) using single-cell transcriptomic analysis. Integrating two scRNA-seq datasets of mouse brain encompassing S1 neocortex^36,37^, we performed clustering analysis and cell type annotation (Methods, Fig. 1a, Left). Our analysis revealed that *Cacng2* transcripts are predominantly restricted to a specific GABAergic population within *Pvalb* interneuron subclass, *Pvalb Reln Tac1* cluster (Fig. 1a, Right), aligning with prior Pvalb immunostaining^27^. This expression was corroborated by a Patch-seq dataset of cortical GABAergic interneurons, where *Pvalb Reln Tac1* cells were validated as a discrete infragranular GABAergic subtype with morpho-electrophysiological features distinct from classical *Pvalb* interneurons (Fig. 1b)^38^. Double-color FISH staining within S1 validated this finding, showing that *Cacng2-*expressing cortical cells were *Pvalb*-positive, co-labeled with *Tac1* (tachykinin precursor 1), and predominantly localized to layers 5 and 6 (L5/6) (Fig. 1c; Extended Data Fig. 1a, b). Analysis of an early postnatal (P4) mouse cortex scRNA-seq datase_t_^39^ and additional FISH staining within S1 regions at P4 indicated that this cell-specific expression pattern is established early in development (Extended Data Fig. 1d-f; Methods). Notably, no co-labeling of *Cacng2* and *Sst* was observed at either adult or early postnatal age (Extended Data Fig. 1c, g). These findings collectively indicate the early-establishment and specific expression of *Cacng2* within the newly discovered subtype of *Pvalb* interneurons, the *Pvalb Reln Tac1* type in the S1 (hereafter Tac1^+^ interneurons).

**Fig. 1.**
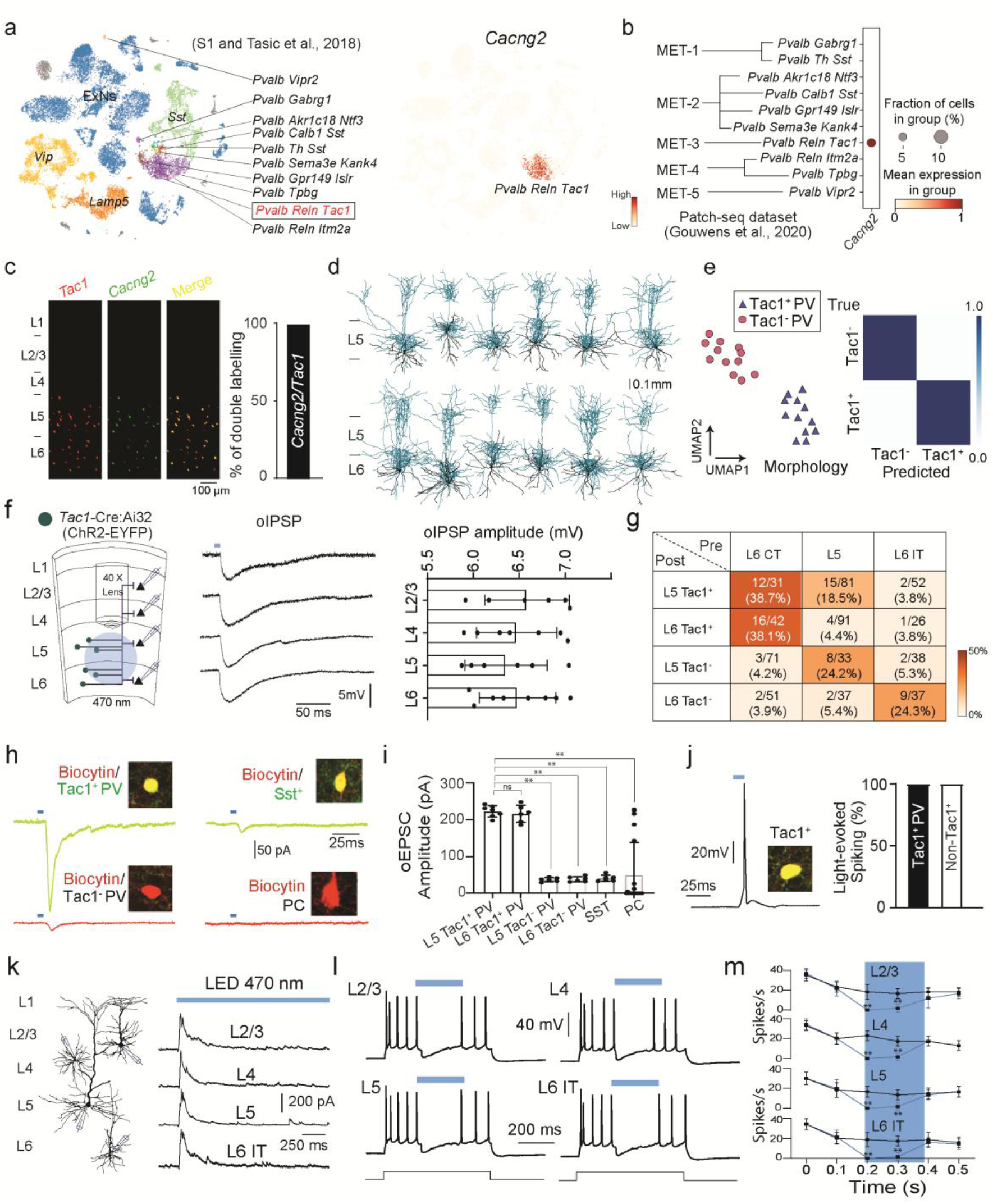
| *Cacng2* expression defines *Tac1*^+^ *Pvalb* subtype that is preferentially recruited by L6 CT neurons. **a,** Left: UMAP representation of two integrated cortex scRNA-seq and snRNA-seq datasets annotated using canonical maker genes shows the presence of cell types and subtypes in the neocortex. Right: UMAP visualization of normalized *Cacng2* expression across the cortical cell types. **b,** Dot plot visualization of *Cacng2* expression in five *Pvalb* MET-types (morphoelectric and transcriptomic type) from the Patch-seq dataset (Gouwens et al., 2020). **c,** Left: Representative micrographs of FISH co-staining for *Tac1* and *Cacng2* in adult somatosensory cortex (S1, parasagittal section). Right: Bar graph depicts proportion of double-labeled cells (*Cacng2^+^/Tac1^+^*) within the total single-labeled cells (*Tac1*^+^ or *Cacng2*^+^) counted from 7 consecutive sections across 6 mice. **d,** Representative morphologies of Tac1^+^ *Pvalb* interneurons in layer 5 (top panel) and 6 (bottom panel). Axon: light blue, and Dendrite: black. The morphology of Tac1*^-^ Pvalb* interneuron morphologies are shown in Extended Data Fig. 2e. **e,** Left: UMAP visualization of Tac1*^+^* (blue triangles) and Tac1*^-^* (red circles) *Pvalb* interneurons based on their morphological features. Right: K-means clustering reveals two distinct morphological subgroups, achieving 100% prediction accuracy for both subgroups. **f,** Left: Schematic diagram illustrating optogenetic activation of *Tac1*-Cre:Ai32 neurons with postsynaptic patch recordings from layers L2/3, L4, L5, and L6. Right: Optogenetic stimulation elicits inhibitory postsynaptic potentials (oIPSPs) across all layers. Representative traces were selected from 7 mice. **g,** Connection matrix summarizing synaptic connectivity rates among layer 6 corticothalamic (CT) neurons, layer 5 pyramidal neurons, layer 6 intratelencephalic (IT) neurons, Tac1^+^ interneurons, and Tac1^-^ interneurons. Note the strongest connectivity from L6 CT neurons onto L5 and L6 Tac1^+^ interneurons. **h,** Optogenetic stimulation of L6 CT neurons in *Ntsr1*-Cre:Ai32 mice evokes strong excitatory postsynaptic currents (oEPSCs) in Tac1⁺ interneurons, with minimal responses in Tac1⁻ interneurons, *Sst* interneurons, and pyramidal cells (PCs). Insets show biocytin-filled recorded neurons together with post hoc immunostaining for Tac1 or Sst. **i,** Quantification of oEPSC amplitudes reveals significantly larger responses in Tac1⁺ neurons than in other cell types (L5 Tac1⁺: n = 7 cells from 6 mice; L6 Tac1⁺: n = 6 cells from 6 mice; L5 Tac1⁻: n = 6 cells from 5 mice; L6 Tac1⁻: n = 6 cells from 5 mice; Sst: n = 6 cells from 6 mice; PCs: n = 21 cells from 8 mice). \*\**p* < 0.01. **j,** Left: Optogenetic activation of L6 CT neurons triggers action potential firing in Tac1⁺ interneurons but not in other recorded cell types. Right: Quantification of the proportion of responsive cells across cell types. **k,** Left: Representative biocytin-recovered pyramidal cells (PCs) recorded from layers L2/3, L4, L5, and L6 IT. Right: Optogenetically evoked inhibitory postsynaptic currents (oIPSCs) recorded in PCs across cortical layers during 1.5 s blue light stimulation of L6 CT neurons. Representative traces were selected from 10 mice. **l,** Optogenetic stimulation of L6 CT neurons suppresses action potential firing in PCs during a +140 pA current injection; blue light was delivered for 200 ms during the depolarizing step. **m,** Quantification of firing inhibition across cortical layers (L2/3: n = 6 cells; L4: n = 5 cells; L5: n = 6 cells; L6 IT: n = 7 cells from 10 animals). \*\**p* < 0.01.

To investigate this *Cacng2*-expressing novel inhibitory subtype (Tac1^+^ interneurons), we began by examining its potential specific Cre driver, the *Tac1*-Cre mouse line^40–42^. Cortical *Tac1*-Cre expression was stable throughout development, was detected as early as postnatal day 14 (P14) and was primarily restricted to deep laminae with only sparse labeling in layer 2/3 (Extended Data Fig. 2a). Double FISH staining for Cre and cardinal interneuron markers revealed that *Tac1*-Cre co-localizes with *Pvalb* but lacks overlap with *Sst*, *Vip*, or *Npy* (Methods; Extended Data Fig. 2b), aligning with prior reports that this line delineates immature *Pvalb* interneurons^40,41^. Notably, *Tac1*-Cre was not detectable in *Pvalb* interneurons within the thalamic reticular nucleus (RTN), a thalamic node for spike-wave discharge (SWD)^16^ (Extended Data Fig. 2c). Whole-cell recordings and morphological reconstructions of Cre-positive cells in the deep layers of S1 demonstrated that their anatomical location, electrophysiological properties, and morphology consistently matched established benchmarks for this specific subtype^38^ (Fig. 1d, e; Extended Data Fig. 2d-f; Extended Data Table 1). Finally, all Cre-positive cells in the deep layers co-stained for stargazin (the protein encoded by *Cacng2*), whereas the sparse supragranular positive cells are mostly stargazin-negative (Extended Data Fig. 2g, h). This provides further evidence that the *Tac1*-Cre line is a reliable tool for selectively targeting *Cacng2*-expressing Tac1^+^ interneuron subtype within the neocortex.

Tac1^+^ interneurons are located in layers 5 and 6 of the somatosensory cortex, and have a massive translaminar axonal arborization, setting them apart from other (Tac1-negative) *Pvalb* interneurons (Fig. 1d; Extended Data Fig. 2e, f), reminiscent of *Pvalb* interneurons known to mediate sensory gain control downstream of layer 6 corticothalamic (L6 CT) neurons^43–45^. To explore this potential correspondence, we first expressed channelrhodopsin-2 (ChR2) in these Tac1^+^ interneurons (*Tac1*-Cre:Ai32) to map their efferent connectivity.

Focal light stimulation limited to deep layers consistently evoked robust inhibitory postsynaptic potentials (IPSPs) of comparable magnitude in principal cells (PCs) across all cortical layers (Fig. 1f), indicating that these particular interneurons exert powerful translaminar inhibition. To map their afferent connectivity and test whether they are preferentially innervated by layer 6 neurons, we employed multi-patch recordings to examine connections between *Pvalb* interneurons and local pyramidal neurons (Methods). L6 CT neurons were genetically labeled (*Ntsr1*-Cre:Ai9) to distinguish them from L6 intratelencephalic (IT) neurons, and Tac1^+^ interneurons were identified via *post hoc* immunostaining. Connectivity rate analysis revealed that Tac1^+^ interneurons receive the highest synaptic input from L6 CT neurons, with other cell types showing minimal projection to them (Fig. 1g). Furthermore, expressing ChR2 in L6 CT neurons (*Ntsr1*-Cre:Ai32) followed by light stimulation evoked robust excitatory postsynaptic currents (EPSCs) in Tac1^+^ interneurons, whereas most other cell types (with the exception of L5a PCs) exhibited only minimal EPSCs (Fig. 1h, i)^46^. In current clamp mode, all Tac1^+^ interneurons reliably fired action potentials upon light stimulation, a response not observed in other cell types (Fig. 1j). As anticipated, light stimulation also evoked widespread translaminar inhibition of PCs across all layers, a process mediated by Tac1^+^ interneurons (Fig. 1k-m). Collectively, these data identify Tac1^+^ interneurons as the primary mediators of L6 CT-driven translaminar inhibition, a circuit mechanism known for sensory gain control^43–45^.

### Stargazer mice exhibit “dormant” Tac1^+^ interneurons resulting in the loss of translaminar inhibition

Given the specificity of *Cacng2* expression in Tac1^+^ interneurons, we then tested whether *stg* cortex exhibits synaptic transmission deficits in these cells due to impaired dendritic AMPA receptor (AMPAR) trafficking by mutated stargazin^27,47,48^. We began by recording and analyzing spontaneous excitatory postsynaptic currents (sEPSCs) in *Pvalb* interneurons within the somatosensory cortex (S1) of adult stargazer *(stg)* and wild-type (WT) littermates. *Pvalb* interneurons were identified by their fast-spiking characteristics and confirmed by *post hoc* morphology recovery (Extended Data Fig. 3a; Methods). Tac1^+^ interneurons were identified either through genetic labeling (crossing *Tac1*-Cre with *stg* mice) or *post hoc* Tac1 immunostaining of fast-spiking neurons recorded from deep layers (Methods). No significant changes in sEPSC amplitude or frequency were detected in fast-spiking *Pvalb* interneurons in superficial layers (L2-L4) between *stg* and WT mice (Extended Data Fig. 3a). However, in *stg* mice there was a marked reduction in both the amplitude and frequency of sEPSCs in L5/6 *Pvalb* interneurons that were Tac1-positive compared with WT, while no significant changes were detected in Tac1-negative *Pvalb* interneurons (Fig. 2a-d). This finding strongly suggests excitatory synapses onto Tac1^+^ interneurons are selectively impaired by the *Cacng2* mutation, consistent with the *Cacng2* expression pattern (Fig. 1). Mapping synaptic connections between Tac1^+^ interneurons and L5/6 pyramidal cells (PCs) with multi-patch recordings further confirmed this selective synaptic disruption in *stg* mice (Fig. 2e, f; Methods). Interestingly, there was no difference in either the amplitude or frequency of N-methyl-D-aspartate (NMDA) receptor-mediated miniature EPSCs in Tac1^+^ interneurons between *stg* and WT (Extended Data Fig. 3b), indicating that excitatory synaptogenesis on Tac1^+^ interneurons proceeds independently of stargazin, and that the observed synaptic deficits stem from “silent” synapses due to impaired AMPAR trafficking. In addition, inhibitory connections from Tac1^+^ interneurons to PCs remained unchanged (Fig. 2f), indicative of their intact efference despite the loss of excitatory drive. In addition to afferent synaptic deficits, Tac1^+^ interneurons in *stg* exhibited a pronounced reduction in intrinsic excitability, as evidenced by significant increases in rheobase and markedly reduced firing frequency compared to WT (Extended Data Fig. 3c, d; Table 1). In light of the established activity-dependent plasticity of *Pvalb* interneuron excitability^49,50^, this attenuated membrane excitability likely reflect a causal, downstream consequence of the lost excitatory synaptic drive.

**Fig. 2.**
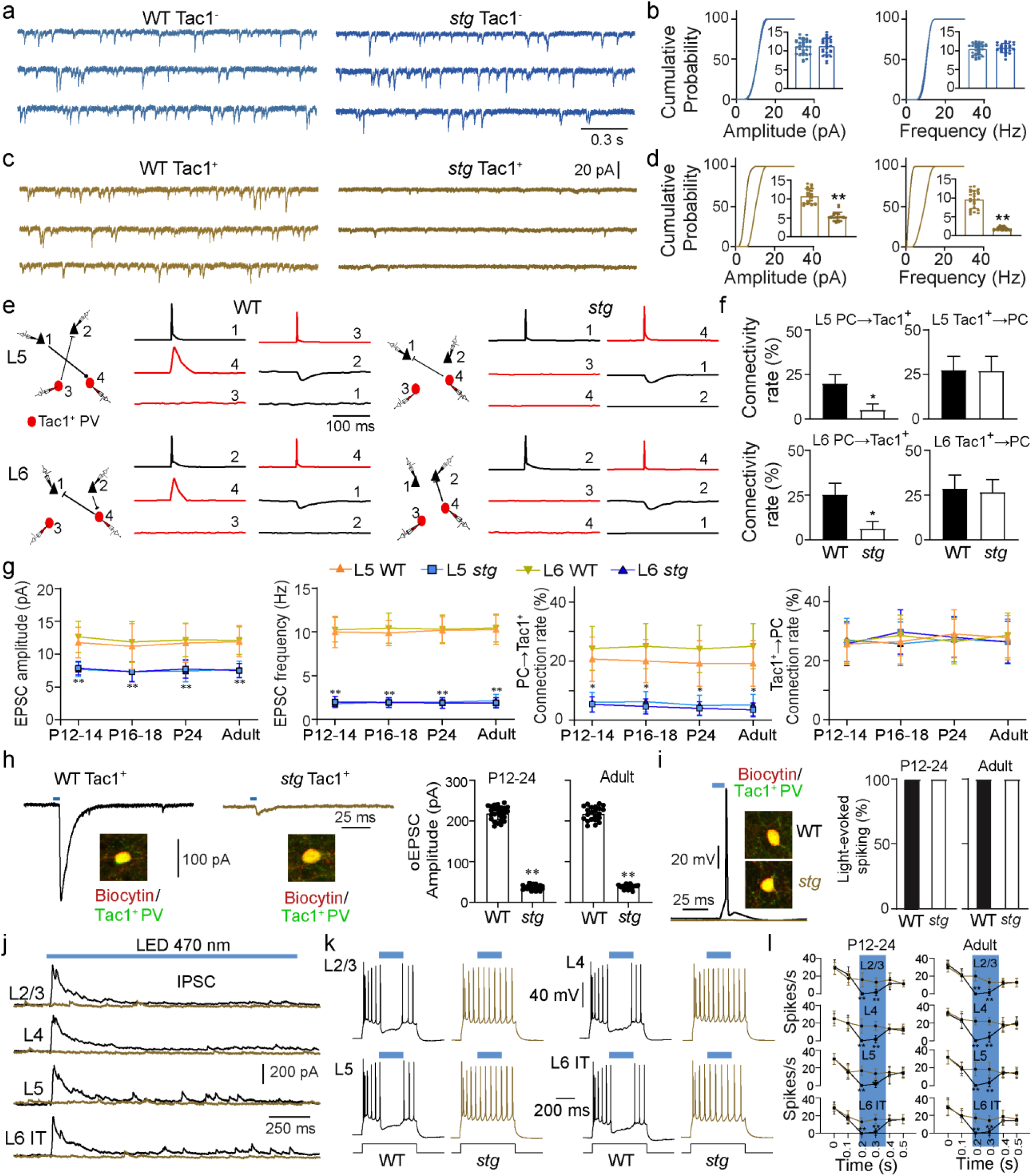
| Stargazer mice exhibit dormant Tac1^+^ *Pvalb* interneurons and loss of L6-driven translaminar inhibition. **a,** Representative traces of spontaneous excitatory postsynaptic currents (sEPSCs) recorded from fast-spiking, Tac1^-^ *Pvalb* interneurons in deep cortical layers (L5/6) from *Tac1*-Cre:Ai9 mice. **b,** Cumulative probability plots and inset bar graphs summarizing sEPSC amplitude and frequency in Tac1^-^ *Pvalb* interneurons from WT and *stg* mice. No significant differences were observed. WT, n = 34 cells from 12 mice; *stg*, n = 27 cells from 11 mice. **c,** Representative traces of sEPSCs recorded from Tac1^+^ *Pvalb* interneurons in deep cortical layers (L5/6) identified in *Tac1*-Cre:Ai9 mice. **d,** Cumulative probability plots and inset bar graphs summarizing sEPSC amplitude and frequency in Tac1^+^ *Pvalb* interneurons from WT and *stg* mice. Student’s t-test; \*\**p* < 0.01. WT, n = 24 cells from 10 mice; *stg*, n = 22 cells from 12 mice. **e,** Representative examples of simultaneous four-cell recordings from pyramidal cells (PCs; black arrows) and Tac1^+^ *Pvalb* interneurons (red circles) in L5 or L6 from WT and *stg* mice. Scale bars indicate injected current (nA), action potential amplitude (mV), and unitary EPSP or IPSP amplitude (mV). **f,** Summary of connection probabilities for PC→Tac1^+^ and Tac1^+^→PC pairs in L5 and L6 in WT and *stg* mice. PC→Tac1^+^ connectivity was significantly reduced in *stg* mice, whereas Tac1^+^→PC connectivity was preserved. PC→Tac1^+^ in L5: 12/61 in WT, 2/40 in *stg*; Tac1^+^→PC in L5: 9/33 in WT, 8/30 in *stg*; PC→Tac1^+^ in L6: 11/44 in WT, 2/33 in *stg*; Tac1^+^→PC in L6: 10/35 in WT, 10/38 in *stg*. \**p < 0.05.* **g,** Left two panels, Quantification of sEPSC amplitude and frequency in L5 and L6 *Tac1*^+^ interneurons across development (P12–14, P16–18, P24, and adult) in WT and *stg* mice. For P12–14: L5, n = 9 WT and n = 9 *stg*; L6, n = 8 WT and n = 10 *stg*. For P16–18: L5, n = 9 WT and n = 9 *stg*; L6, n = 7 WT and n = 9 *stg*. For P24: L5, n = 10 WT and n = 10 *stg*; L6, n = 9 WT and n = 9 *stg*. For adult: L5, n = 8 WT and n = 8 stg; L6, n = 9 WT and n = 10 *stg*. Student’s t-test; \*\**p* < 0.01. Right two panels, summary of connection probabilities for PC→Tac1^+^ and Tac1^+^→PC pairs across development in WT and *stg* mice. PC→Tac1^+^ connectivity was consistently reduced in *stg* mice, whereas Tac1^+^→PC connectivity remained largely unchanged. PC→Tac1^+^ pairs: P12-14, L5: 6/29 WT, 4/75 *stg*; L6: 8/33 WT, 3/50 *stg*. P16-18, L5: 6/30 WT, 3/65 *stg*; L6: 8/32 WT, 3/48 *stg*. P24, L5: 5/26 WT, 3/75 *stg*; L6: 7/29 WT, 2/40 *stg*. Adult, L5: 5/26 WT, 2/58 *stg*; L6: 8/32 WT, 2/39 *stg*. Tac1^+^→PC pairs: P12–14, L5: 10/39 WT, 9/35 *stg*; L6: 10/38 WT, 10/37 *stg*. P16-18, L5: 9/34 WT, 11/37 *stg*; L6: 12/42 WT, 9/35 *stg*. P24, L5: 9/31 WT, 12/43 *stg*; L6: 9/34 WT, 9/33 *stg*. Adult, L5: 10/36 WT, 10/38 *stg*; L6: 10/35 WT, 9/34 *stg*. \**p < 0.05.* **h,** Optogenetic stimulation of L6 corticothalamic (CT) neurons in *Ntsr1*-Cre:Ai32 mice evoked robust excitatory postsynaptic currents (oEPSCs) in Tac1^+^ interneurons in WT mice, but these responses were markedly reduced in *stg* mice. Right, quantification of oEPSC amplitude in Tac1^+^ interneurons. For P12-24 mice: WT, n = 26 cells from 8 mice; *stg*, n = 24 cells from 7 mice. For adult mice: WT, n = 22 cells from 8 mice; *stg*, n = 18 cells from 9 mice. \*\**p* < 0.01. **i,** Left, Optogenetic activation of L6 CT neurons triggered action potential firing in *Tac1*^+^ neurons in WT mice, but this response was largely absent in *stg* mice. Right, Quantification of the proportion of responsive cells across developmental stages (P12-24 and adult), showing robust responses in WT but loss of responsiveness in *stg* mice. **j,** Representative optogenetically evoked inhibitory postsynaptic currents (oIPSCs) recorded from pyramidal cells across cortical layers during 1.5 s blue-light stimulation of L6 CT neurons. Clear oIPSCs were observed in WT mice but were largely absent in *stg* mice. Representative traces were obtained from 11 mice. **k,** Optogenetic stimulation of L6 CT neurons suppressed action potential firing in pyramidal cells during a +140 pA depolarizing current injection; blue light was delivered for 200 ms during the current step. This suppression was robust in WT mice but absent in *stg* mice. **l,** Quantification of light-induced firing inhibition in pyramidal cells across cortical layers in juvenile (P12–24) and adult mice. For P12-24 mice: L2/3, n = 7 WT cells from 5 mice and n = 7 *stg* cells from 6 mice; L4, n = 6 WT cells from 5 mice and n = 6 *stg* cells from 6 mice; L5, n = 6 WT cells from 6 mice and n = 6 *stg* cells from 5 mice; L6, n = 7 WT cells from 6 mice and n = 7 *stg* cells from 5 mice. For adult mice: L2/3, n = 7 WT cells from 6 mice and n = 7 *stg* cells from 7 mice; L4, n = 5 WT cells from 5 mice and n = 5 *stg* cells from 5 mice; L5, n = 6 WT cells from 5 mice and n = 6 *stg* cells from 5 mice; L6, n = 7 WT cells from 7 mice and n = 7 *stg* cells from 6 mice. \*\**p* < 0.01.

Because Tac1^+^ interneurons are innervated primarily by L6 CT neurons to exert robust translaminar inhibition (Fig. 1), we reasoned that AMPAR-mistrafficked Tac1^+^ interneurons in *stg* mice would therefore disrupt this inhibitory microcircuit. To test this, we expressed ChR2 in L6 CT neurons in *stg* and applied light stimulation. In WT littermates, as expected, this stimulation evoked large excitatory postsynaptic potentials (EPSPs) in Tac1^+^ interneurons inducing them to reliably fire action potentials and drive translaminar inhibition of principal cells (PCs) across all cortical layers (Fig. 1h-m; Fig. 2h-l). In contrast, light stimulation in *stg* mice elicited minimal EPSPs in Tac1^+^ interneurons (Fig. 2h), and these cells completely failed to fire action potentials in response to light (Fig. 2i). Consequently, L6 CT-driven translaminar inhibition was virtually abolished in *stg* mice (Fig. 2j-l). Collectively, the loss of excitatory drive, combined with intrinsic hypoexcitability, renders *stg* Tac1^+^ interneurons “dormant”, leading to severely impaired L6 CT-initiated translaminar inhibition within the S1 cortex.

Since *stg* mice do not fully develop SW seizures until approximately three weeks after birth^51,52^, we also reasoned that if the observed circuit disruption directly underlies seizure expression, their developmental onsets should coincide. To test this, we examined the excitatory synapses of Tac1^+^ interneurons at different developmental stages: before seizure onset (P12), around seizure emergence (P17), and after seizure establishment (P>21)^52^. Surprisingly, excitatory synaptic deficits of Tac1^+^ interneurons and the loss of L6 CT-initiated translaminar inhibition in *stg* were consistent across all ages tested. These deficits were detectable as early as P12 (Fig. 2g-i, l), occurring one week before seizure onset. This early manifestation aligns with the developmental trajectory of *Cacng2* expression in wild type mice (Extended Data Fig. 1) and congenital *Cacng2* mutations in *stg* mice. Therefore, functional synaptic deficits of Tac1^+^ interneurons appear to be a primary, cell-autonomous consequence of the mutation that is required, but not solely sufficient for seizure initiation.

### Synaptic dysfunction of deep-layer *Sst* interneurons tracks seizure emergence in *stargazer* mice

Since primary deficits restricted to a subset of cortical *Pvalb* interneurons (Tac1^+^ interneurons) appear insufficient for seizure expression, additional synaptic and circuit adaptations are likely required. To identify these, we examined excitatory synaptic transmission (sEPSCs) of principal cells (PCs) and other major cardinal interneuron types (*Sst*, *Vip*, and *Npy*) across all cortical layers^53^. No significant synaptic changes were observed in most of the cell types examined (Extended Data Fig. 4 and Table 1; Methods), indicating that widespread excitatory connectivity changes are not a feature of this model. However, we discovered profound synaptic deficits in *Sst* interneurons that were localized within layer 5 and layer 6 (L5/6), a cell population that does not normally express *Cacng2* (Extended Data Fig. 1). Whole-cell recordings targeting genetically labeled *Sst* interneurons across the layers revealed that L5/6 *Sst* interneurons in *stg* mice received markedly less excitatory input than those in WT littermates (Fig. 3a), while those in superficial layers remained unaffected (Extended Data Fig. 5a). In concordance, mapping synaptic connections between *Sst* interneurons and PCs across the layers via multi-patch recordings revealed a marked reduction in afferent connectivity from L5/6 PCs onto *Sst* interneurons (Fig. 3b), with no significant changes observed in their superficial counterparts (Extended Data Fig. 5b). In addition to the loss of afferent synaptic inputs, these same *Sst* interneurons exhibited reduced efferent connectivity onto local PCs (Fig. 3b) as well as a significant decrease in their intrinsic excitability (Extended Data Fig. 5c and Table 1).

**Fig. 3.**
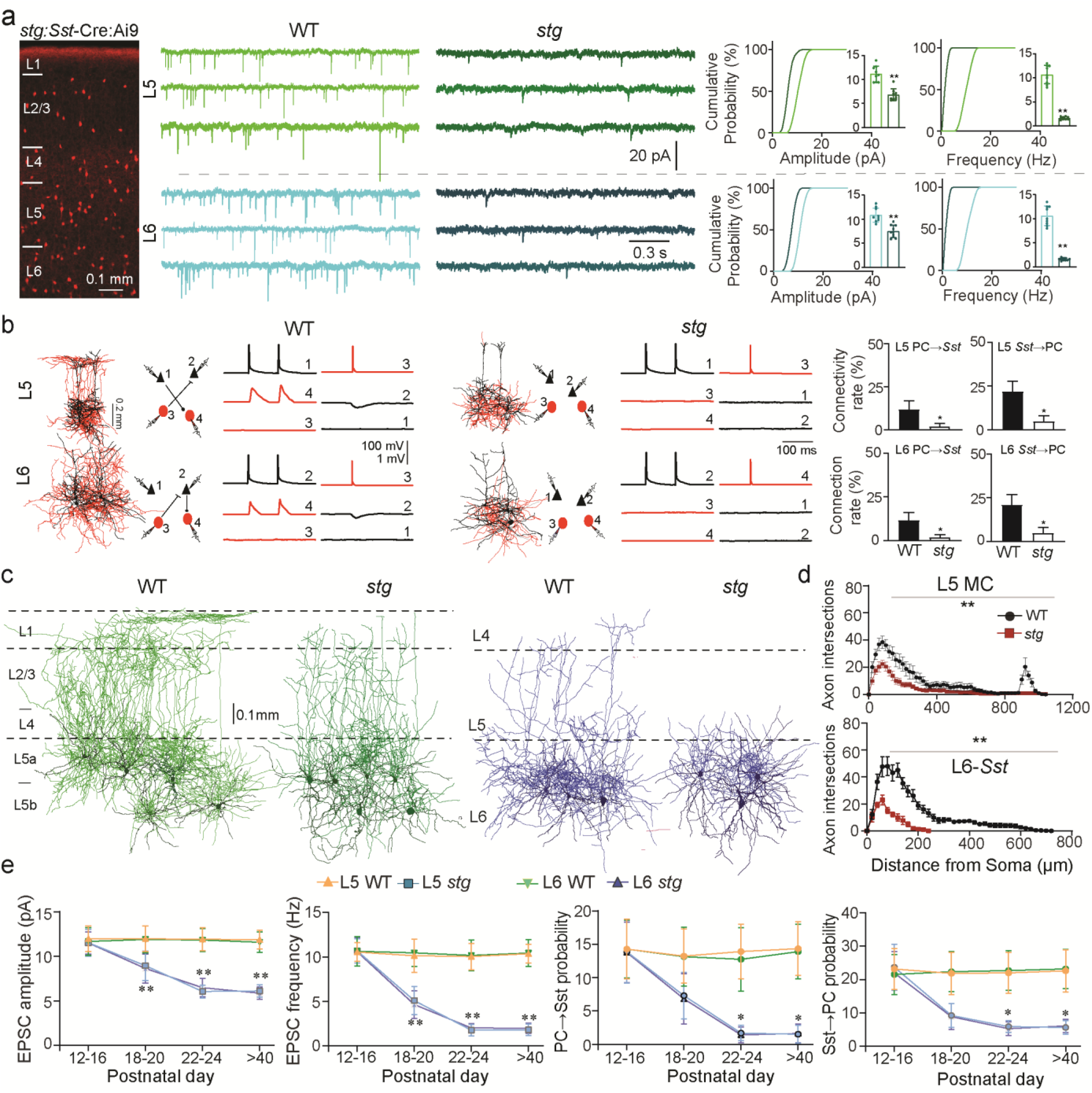
| Stargazer mice exhibit synaptic disruption of deep-layer *Sst* interneurons that track seizure development. **a,** From left to right: representative fluorescence image of the primary somatosensory cortex (S1) from an *stg*:*Sst*-Cre:Ai9 mouse showing *Sst* interneurons (red); representative traces of spontaneous excitatory postsynaptic currents (sEPSCs) recorded from *Sst* interneurons across cortical layers in WT and *stg* mice; and cumulative probability plots with inset bar graphs summarizing sEPSC amplitude and frequency. Cells for WT and *stg*, respectively: L2/3, n = 10 and 8; L4, n = 10 and 9; L5, n = 9 and 9; L6, n = 9 and 10. Student’s t - test; \*\**p* < 0.01. **b,** Left: Representative examples of simultaneous four-cell recordings from principal cells (PCs; black arrows) and *Sst* interneurons (red circles) within each cortical layer in WT and *stg* mice. Scale bars indicate injected current (nA), action potential amplitude (mV), and unitary EPSP or IPSP amplitude (mV). Right: Summary of connection probabilities for PC→*Sst* and *Sst*→PC pairs across layers in WT and *stg* mice. Connectivity between PCs and Sst interneurons was selectively reduced in deep layers in *stg* mice. PC→*Sst* in L5: 5/42 in WT, 1/52 in *stg*; *Sst*→PC in L5: 10/46 in WT, 2/40 in *stg*. PC→*Sst* in L6: 6/50 in WT, 1/59 in *stg*; *Sst*→PC in L6: 10/48 in WT, 2/40 in *stg*. \**p* < 0.05. **c,** Representative morphological reconstructions of L5 Martinotti cells (MCs) and L6 *Sst* interneurons in S1 from WT and *stg* mice. **d,** Sholl analysis comparing axonal arborization of L5 Martinotti cells (top) and L6 *Sst* interneurons (bottom) between WT and *stg* mice. WT, n = 12 L5 MCs and n = 6 L6 *Sst* interneurons; *stg*, n = 12 L5 MCs and n = 6 L6 *Sst* interneurons. *Stg* mice showed a marked reduction in axonal intersections across distances from the soma. \*\**p* < 0.01. **e,** Left two panels, quantification of sEPSC amplitude and frequency in L5 and L6 *Sst* interneurons across development (P12-16, P18-20, P22-24, and P > 40) in WT and *stg* mice. For L5 Sst interneurons: P12-16, n = 20 WT and n = 18 *stg*; P18-20, n = 17 WT and n = 17 *stg*; P22-24, n = 18 WT and n = 18 *stg*; P > 40, n = 19 WT and n = 18 *stg*. For L6 *Sst* interneurons: P12–16, n = 17 WT and n = 17 *stg*; P18–20, n = 17 WT and n = 19 *stg*; P22–24, n = 19 WT and n = 18 *stg*; P > 40, n = 18 WT and n = 19 *stg*. Student’s t-test; \*\**p* < 0.01. Right two panels, summary of connection probability for PC→*Sst* and *Sst*→PC pairs in L5 and L6 across development in WT and *stg* mice. For PC→*Sst* pairs: P12–16, 9/63 in L5 WT, 7/50 in L5 *stg*; 9/64 in L6 WT, 8/58 in L6 *stg*. P18–20, 9/68 in L5 WT, 4/55 in L5 *stg*; 8/61 in L6 WT, 3/44 in L6 *stg*. P22–24, 10/72 in L5 WT, 2/119 in L5 *stg*; 6/47 in L6 WT, 1/76 in L6 *stg*. P > 40, 10/70 in L5 WT, 1/65 in L5 *stg*; 10/72 in L6 WT, 1/61 in L6 *stg*. For *Sst*→PC pairs: P12–16, 11/48 in L5 WT, 9/38 in L5 *stg*; 10/46 in L6 WT, 9/41 in L6 *stg*. P18-20, 10/46 in L5 WT, 4/43 in L5 *stg*; 10/45 in L6 WT, 4/45 in L6 *stg*. P22–24, 10/46 in L5 WT, 10/170 in L5 *stg*; 10/44 in L6 WT, 6/115 in L6 *stg*. P > 40, 10/44 in L5 WT, 7/124 in L5 *stg*; 10/43 in L6 WT, 9/148 in L6 *stg*. \**p* < 0.05.

L5/6 *Sst* interneurons are heterogeneous and encompass multiple distinct subtype_s_^54^, raising the question of whether these impairments are pervasive across the *Sst* population or subtype specific. By broadly grouping *Sst* interneurons into Martinotti cells (MCs) and non-Martinotti cells (NMCs) based on their unique morpho-electrophysiological properties^55,56^, we found that the deficits were confined to MCs, sparing the quasi-fast- spiking NMCs (Extended Data Fig. 5d). Consistent with the diminished efferent connectivity, reconstruction of MCs and layer 6 *Sst* interneurons from *stg* and WT revealed morphological aberrations, highlighted by a significant reduction in axonal arborizations (Fig. 3c). This morphological deficit appeared to generalize across L5 MCs, encompassing both fan-out MC and T-shaped MCs (Fig. 3c)^54,55^. To mitigate slicing artifacts and confirm these findings using an unbiased *in vivo* approach, we crossed *stg* mice with the X98 line, which specifically labels T-shaped MCs^57^. This strategy revealed a near-complete loss of MC axons within layer 1 in *stg* cortex (Extended Data Fig. 5e), suggesting that synaptic inhibition directed to the superficial layers is particularly impacted. Collectively, the loss of excitatory input, diminished inhibitory output, and intrinsic hypoexcitability constitute a severe and selective functional impairment of L5/6 *Sst* interneurons in the *stg* model. Notably, this impairment did not extend to their own inhibitory inputs, since inhibitory postsynaptic currents (IPSCs) recorded in these *Sst* neurons and afferent connectivity from nearby *Vip* interneurons, their primary source of inhibition, remained unchanged (Extended Data Fig. 5f, g) ^58,59^.

Given the absence of *Cacng2* expression in *Sst* interneurons, the synaptic deficits observed in these cells in *stg* cortex likely represent secondary circuit remodeling necessary for seizure generation. If true, these non-cell autonomous deficits should emerge in tandem with the developmental onset of seizures. To investigate this, we assessed the excitatory afference and the inhibitory efference of *Sst* interneurons across the three developmental stages. We found that reductions in the connectivity of L5/6 *Sst* interneurons were not present at P14, and only became apparent at the initial emergence of seizures (P18-P20)^52^, and persisted long into adulthood (Fig. 3e). The temporal alignment between *Sst* interneuron synaptic dysfunction and seizure development one week following the primary Tac1-Pvalb defect strongly suggests that these changes are a secondary circuit adaptation, potentially acting as an additional requisite component for epileptogenesis.

### Chemogenetic activation of Tac1^+^ *Pvalb* interneurons abolishes *stg* seizures

We have identified a two-stage pathological progression in *stg* mice, beginning with the early postnatal presence of “dormant” Tac1^+^ interneurons, followed by a delayed, secondary synaptic dysfunction of L5/6 *Sst* interneurons that temporally coincides with the developmental onset of seizures. This suggests that the generation and expression of SW seizures in *stg* mice require both abnormalities and reversing either one may abolish the seizure phenotype. To test this, we reversed the early functional dormancy of Tac1^+^ interneurons using a chemogenetic (DREADD) approach. Given their intact efferent connectivity (Fig. 2), chemogenetic activation of these neurons would be expected to restore the downstream translaminar inhibition and attenuate seizures. We first generated mice that express the hM3Dq (Gq) receptor under the *Tac1* promoter (*Tac1*-Cre:Gq) to selectively target Gq expression in Tac1^+^ interneurons. This strain was then crossed with *stg* mice to generate *stg*:*Tac1*-Cre:Gq mice. We validated the DREADD approach by administering clozapine-N-oxide (CNO) to L5/6 *Pvalb* interneurons recorded from these mice *in vitro* and found that only those neurons expressing Gq under *Tac1*-Cre promoters (fluorescence-positive) could be depolarized by CNO, confirming the specificity of DREADD expression in our mouse model (Extended Data Fig. 6a). Baseline electroencephalography (EEG) recordings in these mice prior to CNO administration showed characteristic spike-wave discharges (SWD) within the 5-9 Hz range (Fig. 4a, b, e, f). However, a single intraperitoneal injection of 0.1 mg/kg CNO eliminated SWDs for approximately 3 hours (Fig. 4a-d, g, h). Power spectrum analysis showed a significant decrease in seizure waveform activity within the 5-9 Hz frequency band following CNO administration, whereas interictal gamma oscillations were significantly enhanced (indicative of *Pvalb* interneuron activation; Fig. 4f_)_^60^. These effects were consistent across all experimental mice (n = 7), whereas no change in seizure activity was observed in *stg*:*Tac1*-Cre mice lacking hM3Dq expression (Control, n = 6, Fig. 4g, h). In addition, a similar therapeutic effect could be achieved by providing *stg*:*Tac1*-Cre:Gq mice with *ad libitum* access to CNO-laced water (40 mg/L_)_^61,62^. These mice exhibited no seizure activity 4 hours after access to CNO-laced water (Extended Data Fig. 6c, d), with no significant changes observed in vestibulo-motor deficits typical of *stg* mice^63^. These findings indicate that selective activation of cortical Tac1*^+^* interneurons alone is sufficient to eliminate absence seizure occurrence in *stg* mice despite deficient *Sst* signaling not being rescued.

**Fig. 4.**
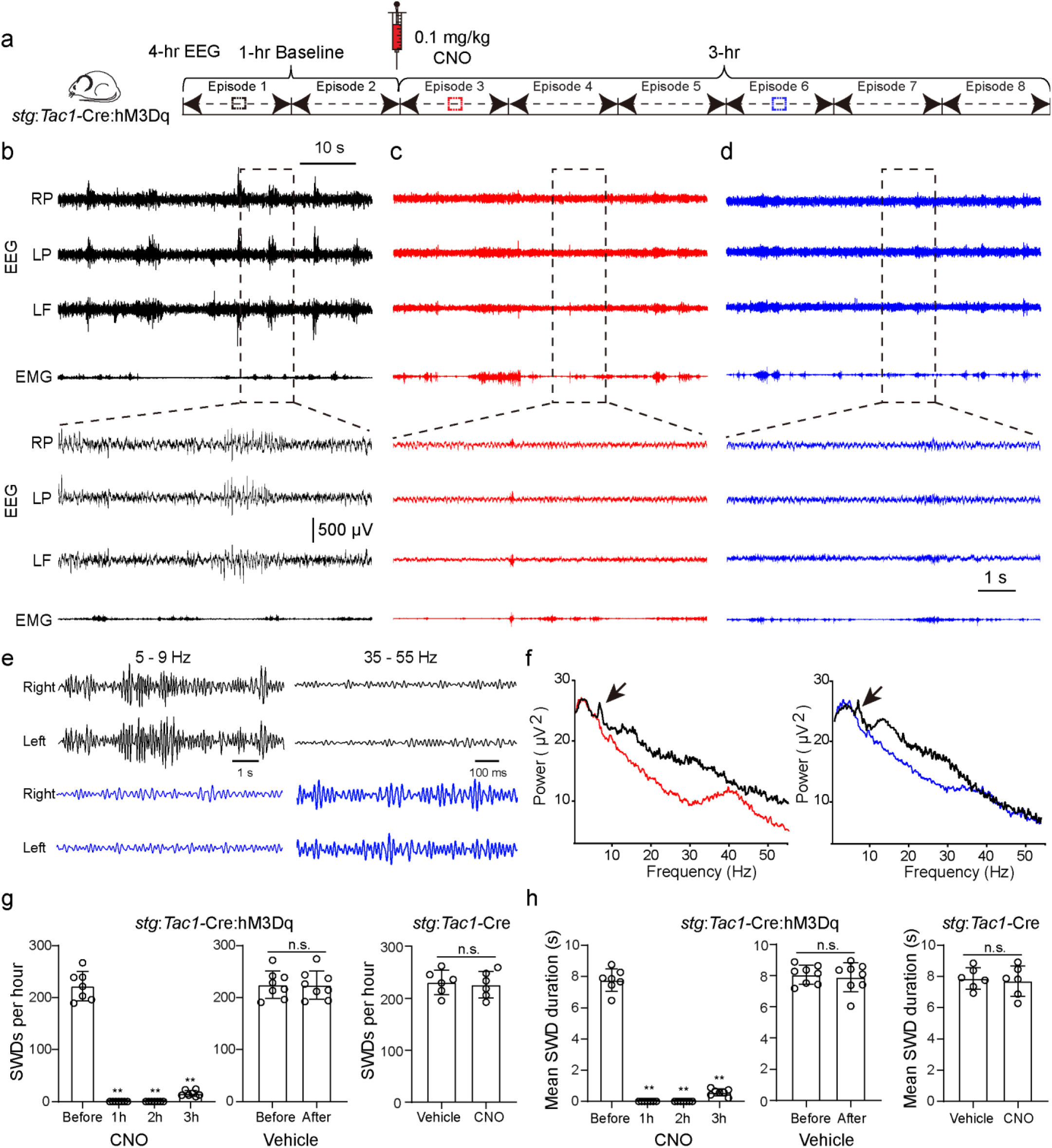
| Chemogenetic activation of Tac1^+^ *Pvalb* interneurons in *stg* inhibits seizure activity. **a,** Schematic of the experimental design detailing electroencephalography (EEG) recordings in *stg*:*Tac1-*Cre:hM3Dq mice. All recording episodes are marked. **b,** Baseline EEG recordings from the left frontal (LF), right parietal (RP), and left parietal (LP) cortices, together with electromyography (EMG), before CNO administration. Bottom, expanded traces showing characteristic spike-wave discharges associated with absence seizures. **c,d,** Representative EEG traces recorded shortly after CNO administration (c, red) and 2 h after CNO administration (d, blue), showing suppression of absence seizure activity across all three cortical recording channels. Bottom, expanded traces. **e,** EEG frequency-band traces before (black) and after chemogenetic activation of Tac1^+^ interneurons (blue) in *stg*:*Tac1*-Cre:hM3Dq mice, showing the disappearance of the dominant 5-9 Hz absence seizure activity after CNO treatment. **f,** Power spectrum analysis of EEG recordings before and after chemogenetic activation of Tac1^+^ interneurons. CNO treatment markedly reduced power in the 5-9 Hz range (black, baseline; red, shortly after activation; blue, 2 h after activation), consistent with seizure suppression (arrows), and was accompanied by increased gamma-band power (35-55 Hz). **g,** Quantification of spike-wave discharges (SWDs) per hour in *stg*:*Tac1*-Cre:hM3Dq mice before and after CNO treatment. Data were analyzed using one-way repeated-measures ANOVA followed by Dunnett’s multiple comparisons test against baseline (Before). A significant effect of time was observed (F(1.060, 6.357) = 389.7, *p* < 0.01). Dunnett’s test showed significant differences between Before and 1 h, 2 h, and 3 h (all \*\**p* < 0.01). n = 7 mice. Vehicle treatment produced no significant change (n = 8 mice, paired t-test). CNO treatment in control *stg*:*Tac1*-Cre mice without hM3Dq expression also produced no significant change (n = 6 mice, Student’s t-test). **h,** Quantification of mean SWD duration in *stg*:*Tac1*-Cre:hM3Dq mice before and after CNO treatment. Data were analyzed using one-way repeated-measures ANOVA followed by Dunnett’s multiple comparisons test against baseline (Before). A significant effect of time was observed (F(1.090, 6.541) = 660.6, *p* < 0.01). Dunnett’s test showed significant differences between Before and 1 h, 2 h, and 3 h (all \*\**p* < 0.01). n = 7 mice. Vehicle treatment produced no significant change (n = 8 mice, paired t-test). CNO treatment in control *stg*:*Tac1*-Cre mice without hM3Dq expression also produced no significant change (n = 6 mice, Student’s t-test).

Tac1 expression is not restricted to the neocortex; it is also expressed in subcortical regions, most notably in the striatum^64,65^ (Extended Data Fig. 7a). To exclude the potential confounding effect of subcortical DREADD expression driven by *Tac1*-Cre in seizure suppression, we investigated *Kcnc2*-Cre as an alternative for targeting L5/6 Tac1^+^ *Pvalb* interneurons^66,67^. *Kcnc2*-Cre expression is sparse at P14 but progressively increases to peak levels by P40 (Extended Data Fig. 6e). Double FISH staining for *Cre* and cardinal interneuron markers revealed that in the neocortex, *Kcnc2*-Cre expression is mostly restricted to deep cortical layers and co-localizes with *Pvalb*, with only sparse overlap in *Sst* and *Npy* interneurons (Extended Data Fig. 6f). Whole-cell recordings confirmed that cells labeled by *Kcnc2*-Cre are predominantly fast-spiking interneurons (i.e., *Pvalb*-positive) (Extended Data Fig. 6g). Notably, *Kcnc2*-Cre was not detectable in *Pvalb* interneurons within the RTN (Extended Data Fig. 6h). Collectively, *Kcnc2*-Cre provides selective access to L5/6 *Pvalb* interneurons while sparing superficial-layer *Pvalb* cortical interneurons, RTN interneurons, and striatal neurons (Extended Data Fig. 7a), thus complementing the *Tac1*-Cre approach. We therefore generated *stg* mice expressing the hM3Dq (Gq) receptor under the *Kcnc2* promoter (*stg*:*Kcnc2*-Cre:hM3Dq). Following *in vitro* verification of hM3Dq (Gq) expression within deep-layer *Pvalb* interneurons (Extended Data Fig. 6i), administration of 0.1 mg/kg CNO in *stg*:*Kcnc2*-Cre:Gq mice abolished seizure activity (n = 7; Extended Data Fig. 6j-l), mirroring the results observed in the *Tac1*-Cre cohort.

Our developmental circuit mapping in *stg* suggests that the early disruption of L6 CT-Tac1^+^ inhibitory microcircuit alone is insufficient for immediate seizure expression but initiates secondary, delayed functional and morphological defects within *Sst* interneurons necessary for seizure emergence. To test this sequential assumption, we mimicked the initial inhibitory microcircuit disruption observed in *stg* by suppressing Tac1^+^ interneurons with a DREADD approach in WT mice. We reasoned that while this targeted suppression would not immediately provoke spike-wave discharges (SWD), it would trigger the secondary *Sst* interneuron remodeling required for seizures to emerge. We generated mice expressing the hM4Di (Gi) receptor under the *Tac1* promoter (*Tac1*-Cre:Gi) to selectively suppress Tac1 interneurons upon CNO administration. DREADD expression specificity was confirmed via *in vitro* whole-cell recordings as done on Gq mice (Extended Data Fig. 6b). To achieve a stable, prolonged inhibition, mice were implanted for EEG recordings and given *ad libitum* access to CNO-laced drinking water (40 mg/L; Fig. 5a). While effective CNO delivery and subsequent interneuron suppression occurred within hours of access to CNO-laced water (Extended Data Fig. 6c-d), EEG recordings showed a total absence of SWDs during the first four days, confirming that loss of translaminar inhibition alone is insufficient for immediate seizure generation. However, continued EEG monitoring detected the appearance of synchronous bilateral cortical SWDs during visually monitored behavioral quiescence on the 5th day of continuous CNO treatment, with SWD frequency peaking between 10-12 days (Fig. 5a, b, e and Extended Data Video 1). These induced events exhibited the hallmarks of typical absence seizures in *stg* mice; SW frequencies predominantly in the 5-9 Hz range (Fig. 5b, d) and a distinct lack of SWD activity in the dorsal striatum (CPu) (Extended Data Fig. 7b-d) as seen in rat models^68,69^. SWD duration and frequency of occurrence were not significantly different from those observed in *stg* mice at peak seizure activity (Fig. 5e).

**Fig. 5.**
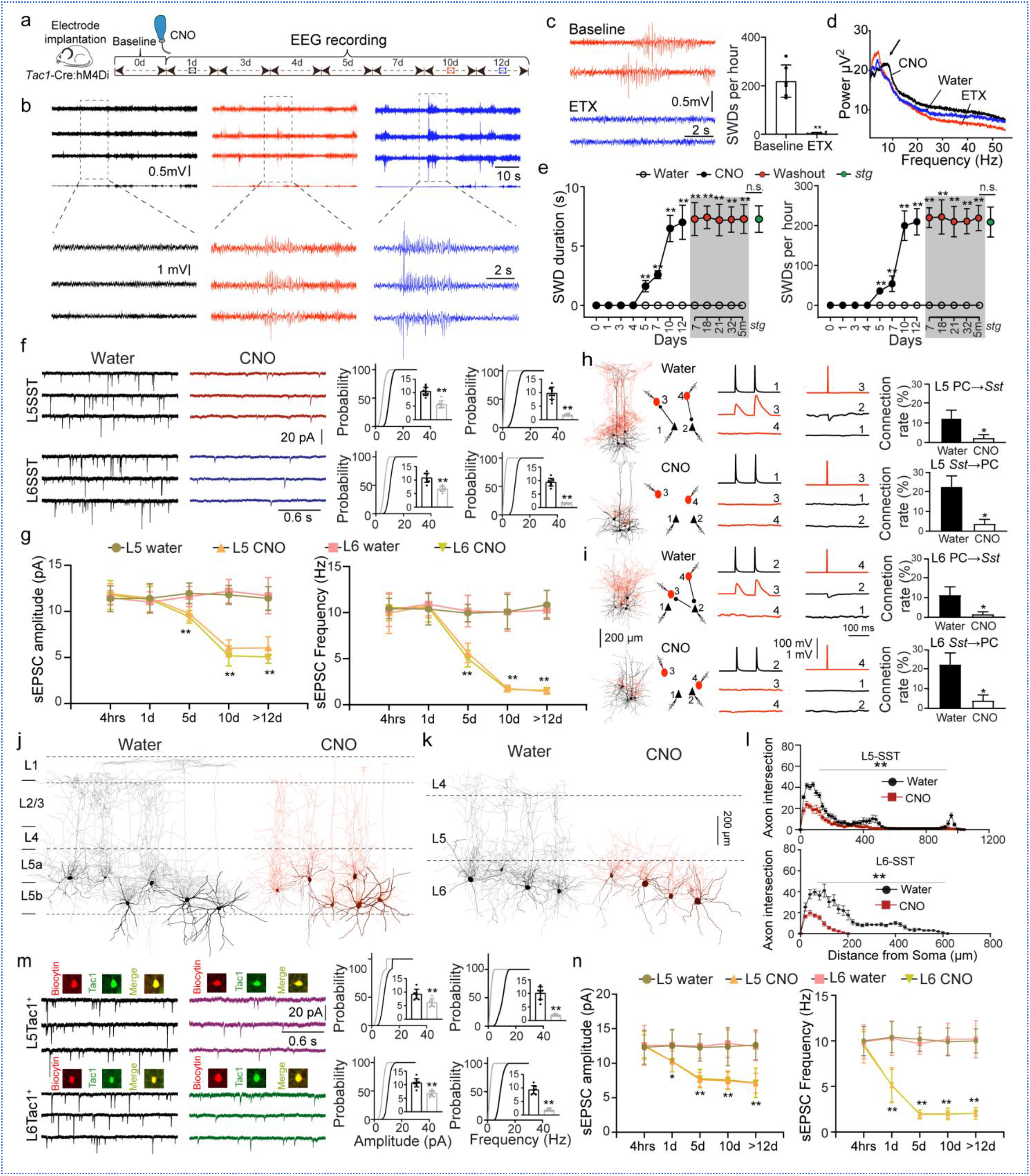
| Chemogenetic inhibition of Tac1^+^ *Pvalb* interneurons in WT leads to *Sst* interneuron defects and absence seizure onset. **a**, Schematic of the experimental design for EEG recordings in WT:*Tac1*-Cre:hM4Di mice. CNO was delivered in drinking water (40 mg/L). Recording sessions across the experimental timeline are indicated. **b,** Representative EEG traces recorded before CNO treatment and after 10 or 12 days of CNO administration in WT:*Tac1*-Cre:hM4Di mice. Expanded traces below illustrate the emergence of spike-wave discharges after chronic CNO treatment. **c,** Left: Representative EEG traces before and after ethosuximide (ETX) administration in WT:*Tac1*-Cre:hM4Di mice after 10 days of CNO treatment. Right: Quantification of spike-wave discharges per hour (SWDs/h) before and after ETX administration, showing marked suppression of seizure activity by ETX. \*\**p* < 0.01; n = 6 mice. **d,** Power spectrum analysis of EEG recordings from WT:*Tac1*-Cre:hM4Di mice under water, CNO, and CNO + ETX conditions. Chronic CNO treatment increased power in the 5-9 Hz range, whereas ETX reduced this seizure-associated activity toward baseline levels. **e,** Daily SWD duration (left) and frequency (right) analyzed from approximately 12:00-13:00 in WT:*Tac1*-Cre:hM4Di mice. Epileptic activity was monitored during 12 days of CNO treatment and across a subsequent plain-water washout phase to assess post-withdrawal seizure persistence. Spontaneous SWDs from *stg* mice were included for comparison with peak induced seizure activity, with no significant differences detected in SWD duration or frequency of occurrence. CNO group, n = 7 mice; water group, n = 5 mice; stg group, n = 5 mice. **f,** Representative traces of spontaneous excitatory postsynaptic currents (sEPSCs) recorded from post hoc-identified L5 (top) and L6 (bottom) Sst interneurons in WT:*Tac1*-Cre:hM4Di mice after 12 days of CNO treatment or in water-treated controls. Right, cumulative probability plots and inset bar graphs summarize sEPSC amplitude and frequency. For L5 *Sst* interneurons: n = 10 cells in Water and n = 8 cells in CNO. For L6 *Sst* interneurons: n = 10 cells in Water and n = 9 cells in CNO. \*\**p* < 0.01. **g,** Quantification of mean sEPSC amplitude (left) and frequency (right) in L5 and L6 *Sst* interneurons at 4 h, 1 day, 5 days, 10 days, and 12 days after CNO administration. For L5 *Sst* interneurons: 4 h, n = 9 CNO and n = 9 Water; 1 day, n = 10 CNO and n = 8 Water; 5 days, n = 8 CNO and n = 9 Water; 10 days, n = 10 CNO and n = 10 Water; 12 days, n = 9 CNO and n = 10 Water. For L6 *Sst* interneurons: 4 h, n = 10 CNO and n = 9 Water; 1 day, n = 9 CNO and n = 10 Water; 5 days, n = 9 CNO and n = 9 Water; 10 days, n = 10 CNO and n = 8 Water; 12 days, n = 8 CNO and n = 10 Water. \*\**p* < 0.01. **h,** Representative examples of simultaneous four-cell recordings from principal cells (PCs; black arrows) and post hoc-verified L5 *Sst* interneurons (red circles) in water-treated mice and mice treated with CNO for 12 days. Representative morphological reconstructions are also shown. Right, summary of connection probabilities for L5 PC→*Sst* and L5 *Sst*→PC pairs in the two groups. PC→*Sst*: 6/46 in Water, 2/80 in CNO; *Sst*→PC: 13/57 in Water, 3/69 in CNO. \**p* < 0.05. **i,** Representative examples of simultaneous four-cell recordings from principal cells (PCs; black arrows) and post hoc-verified L6 *Sst* interneurons (red circles) in water-treated mice and mice treated with CNO for 12 days. Representative morphological reconstructions are also shown. Right, summary of connection probabilities for L6 PC→*Sst* and L6 *Sst*→PC pairs in the two groups. PC→*Sst*: 6/57 in Water, 2/108 in CNO; *Sst*→PC: 11/48 in Water, 2/47 in CNO. \**p* < 0.05. **j,k,** Representative morphological reconstructions of L5 Martinotti cells (j) and L6 Sst interneurons (k) from WT:*Tac1*-Cre:hM4Di mice treated with water or CNO for 12 days. **l,** Sholl analysis comparing axonal arborization of L5 Martinotti cells (top) and L6 *Sst* interneurons (bottom) between water-treated and CNO-treated WT:*Tac1*-Cre:hM4Di mice. Water: n = 13 L5 MCs and n = 7 L6 *Sst* interneurons; CNO: n = 12 L5 MCs and n = 6 L6 *Sst* interneurons. Chronic CNO treatment caused a marked reduction in axonal intersections across distances from the soma. \*\**p* < 0.01. **m,** Left: Representative images showing biocytin-filled recorded neurons followed by Tac1 immunostaining to verify Tac1^+^ identity. Representative traces of sEPSCs recorded from L5 (top) and L6 (bottom) Tac1^+^ interneurons in WT:*Tac1*-Cre:hM4Di mice treated with CNO for 12 days or in water-treated controls. Right: Cumulative probability plots and inset bar graphs summarize sEPSC amplitude and frequency in Tac1^+^ interneurons. For L5 Tac1^+^ interneurons: n = 9 cells in Water and n = 8 cells in CNO. For L6 Tac1^+^ interneurons: n = 9 cells in Water and n = 9 cells in CNO. \*\**p* < 0.01. **n,** Quantification of mean sEPSC amplitude (left) and frequency (right) in L5 and L6 Tac1^+^ interneurons at 4 h, 1 day, 5 days, 10 days, and 12 days after CNO administration. For L5 Tac1^+^ interneurons: 4 h, n = 9 CNO and n = 9 Water; 1 day, n = 10 CNO and n = 9 Water; 5 days, n = 9 CNO and n = 8 Water; 10 days, n = 9 CNO and n = 9 Water; 12 days, n = 10 CNO and n = 10 Water. For L6 Tac1^+^ interneurons: 4 h, n = 8 CNO and n = 9 Water; 1 day, n = 9 CNO and n = 10 Water; 5 days, n = 10 CNO and n = 9 Water; 10 days, n = 8 CNO and n = 10 Water; 12 days, n = 9 CNO and n = 10 Water. *p* < 0.05, \*\**p* < 0.01.

Administration of the standard treatment absence epilepsy, ethosuximide (ETX, 20 mg/kg), completely abolished seizure activity in CNO-administered mice^70^, further confirming the similarity of these events (Fig. 5c, d). Motor deficits evaluated by parallel rod and rotarod tests were not detected in these mice, suggesting that the ataxic phenotype in the genetic model is accounted for by *stg* deficits outside of the cortex (Extended Data Fig. 7e,f)^63^. The trajectory of seizure development in these adult WT mice mirrors the progression seen in *stg* mice; while the initial inhibitory loss is present early, seizures emerge only after a predictable delay.

Remarkably, when mice were returned to regular drinking water after 12 days of CNO administration, the seizures in CNO-administered mice persisted for up to 5 months (Fig. 5e), indicating that a temporary two-week loss of translaminar inhibition induces long-lasting functional circuit changes essential for sustaining chronic seizures.

To confirm that the secondary circuit dysfunction necessary for seizure generation lies within deep-layer *Sst* interneurons, we analyzed spontaneous excitatory postsynaptic currents (sEPSCs) recorded in L5/6 interneurons from somatosensory cortical slices of WT mice following varying durations of free access to CNO-laced water. We initially selected presumptive *Sst* interneurons on the basis of their spiking properties and subsequently verified their identity via *post hoc* immunostaining for Sst (Extended Data Fig. 7g; Methods). Compared to WT mice with regular drinking water and different durations of CNO treatment (4 hours, 1 day, 5 days, 10 day and > 12 day), we observed a progressive decline in both the amplitude and frequency of sEPSCs, which reached a nadir at 10 days, coinciding with full development of the seizure phenotype (Fig. 5f, g), a temporal pattern that closely mirrors the pathophysiology observed in *stg* mice. To further confirm this synaptic deficit, we performed multi-patch recordings of *Sst* interneurons (identified by *post hoc* immunostaining) and PCs in L5/6 in mice following CNO-induced seizures. We identified a marked reduction in the afferent connectivity of local PCs onto *Sst* interneurons (Fig. 5h, i). Furthermore, these same *Sst* interneurons exhibited reduced afferent connectivity onto local PCs and reduced intrinsic excitability (Fig. 5h, i and Extended Data Table 1). Finally, we reconstructed the morphology of *Sst* interneurons from an additional cohort of mice following CNO-induced seizures. We found that L5/6 *Sst* interneurons in these mice with pharmacologically-induced seizures had lost a significant portion of their axonal arborization, particularly within layer 1 (Fig. 5j-l). Together, these data demonstrate that pharmacologically induced seizures fully recapitulate the *Sst* interneuron phenotype observed in *stg* mice.

The observation that seizures persist beyond CNO treatment in treated WT mice suggests long-lasting circuit reorganization in their neocortex. To investigate this, we extended our recordings of *Sst* interneurons following CNO discontinuation. Notably, *Sst* interneurons exhibited the same synaptic deficits observed during CNO treatment, suggesting that this circuit reorganization is likely permanent (Extended Data Fig. 7g, h). Five months after CNO withdrawal, the broader *Sst* abnormalities remained evident. PC→*Sst* and *Sst*→PC connectivity remained markedly reduced in both L5 and L6, accompanied by persistently reduced intrinsic excitability, including decreased firing responses and maximal action potential number and increased rheobase (Extended Data Fig. 7i, j). Sholl analysis further revealed a persistent reduction in axonal arborization in both L5 and L6 *Sst* interneurons (Extended Data Fig. 7k). These findings demonstrate that both the functional and structural abnormalities of deep-layer *Sst* interneurons persist long after CNO withdrawal. Although CNO discontinuation should relieve Tac1^+^ interneurons from inhibition, the persistence of seizures argues that temporary chemogenetic inhibition can induce permanent plastic changes in these cells themselves. To test this, we analyzed spontaneous excitatory postsynaptic currents (sEPSCs) in Tac1^+^ interneurons (identified via *post hoc* immunostaining; Fig. 5m) from the mice with pharmacologically induced seizures, revealing a significant reduction in both amplitude and frequency. This indicates that the suppression of Tac1^+^ interneuron activity drives a selective (do you need to say “selective”?) loss of their excitatory synaptic inputs (Fig. 5m, n), reminiscent of an activity-dependent plasticity mechanism for cortical *Pvalb* interneurons^71,72^. Tracking these interneurons across various durations of CNO treatment revealed that these synaptic changes developed gradually during CNO-mediated inhibition (Fig. 5n) and persisted 32 days after CNO withdrawal (Extended Data Fig. 7i, j). Furthermore, immunohistochemical analysis of S1 revealed a significant, selective(ok?) decrease in GluR4 in the excitatory synapses onto Tac1^+^ interneurons following CNO administration, further mirroring the *stg* model (Extended Data Fig. 7k, l_)_^27,73^. Altogether, these data demonstrate that suppressing Tac1^+^ *Pvalb* interneuron activity in WT recapitulated the circuit deficits of *stg* mice that sustain absence seizures.

Finally, we investigated whether *tottering* mice (*tg/tg*), an absence seizure model harboring a loss of function mutation in *Cacna1a* (encoding the P/Q calcium channel alpha subunit), exhibit the same functional circuit deficit as *stg* mice. Since P/Q channels mediate exocytotic neurotransmitter release, we first used whole-cell recordings to analyze baseline excitatory synaptic transmission (sEPSCs) onto cortical interneurons. We focused on *Pvalb* interneurons across different layers in WT and *tg* mice, and Tac1^+^ *Pvalb* interneurons were either genetically labeled or *post hoc* differentiated by immunostaining (Methods). As found in *stg* mice, Tac1^+^ interneurons in *tg* exhibited a marked reduction in sEPSC amplitude and frequency (Fig. 6a) alongside reduced intrinsic excitability (Extended Data Fig. 8b and Table 1) compared to WT interneurons, while Tac1^-^ *Pvalb* interneurons in L5/6 (Fig. 6b) and *Pvalb* interneurons in L2-4 (Extended Data Fig. 8a) were not significantly impacted by the *Cacna1a* mutation. Since Tac1^+^ interneurons are primarily innervated by L6 CT neurons, we next tested whether evoked presynaptic glutamate release from L6 CT terminals is specifically impaired. We expressed ChR2 in L6 CT neurons in *tg* mice (Methods) and applied light stimulation during patch-clamp recording. As expected, optogenetic stimulation of L6 CT neurons in WT littermates evoked large EPSCs in Tac1^+^ interneurons (verified by *post hoc* staining) that reliably triggered action potential and drove translaminar inhibition of PCs (Fig. 6c-f; see also Fig. 1, 2). However, the same stimulation in *tg* mice only evoked minimal EPSCs (Fig. 6c-f). Consequently, *tg* Tac1^+^ interneurons failed to fire action potentials, and downstream L6-driven translaminar inhibition was virtually abolished (Fig. 6c-f). Crucially, these synaptic deficits appeared early in development, independent of and prior to seizure onset, suggesting that they are primary cortical defects due to developmental loss of *Cacna1a* (Fig. 6c, d and Extended Data Fig. 8c). To understand this selective synaptic vulnerability, we analyzed a cortex snRNA-seq dataset and found that *Cacna1a* expression is highly enriched in L6 CT neurons (Extended Data Fig. 8d; Methods). FISH staining further confirmed that this robust expression profile is established as early as postnatal day 4 (P4) (Extended Data Fig. 8e). Together, these data demonstrate that L6 CT-Tac1^+^ interneuron synapses rely almost exclusively on presynaptic P/Q channels for glutamate release, and that the *Cacna1a* mutation causes a failure of these synapses and subsequent translaminar inhibition, likely due to functional compensation failure or an incomplete shift in reliance from P/Q-type to N-type calcium channels for neurotransmitter release^29,74,75^.

**Fig. 6.**
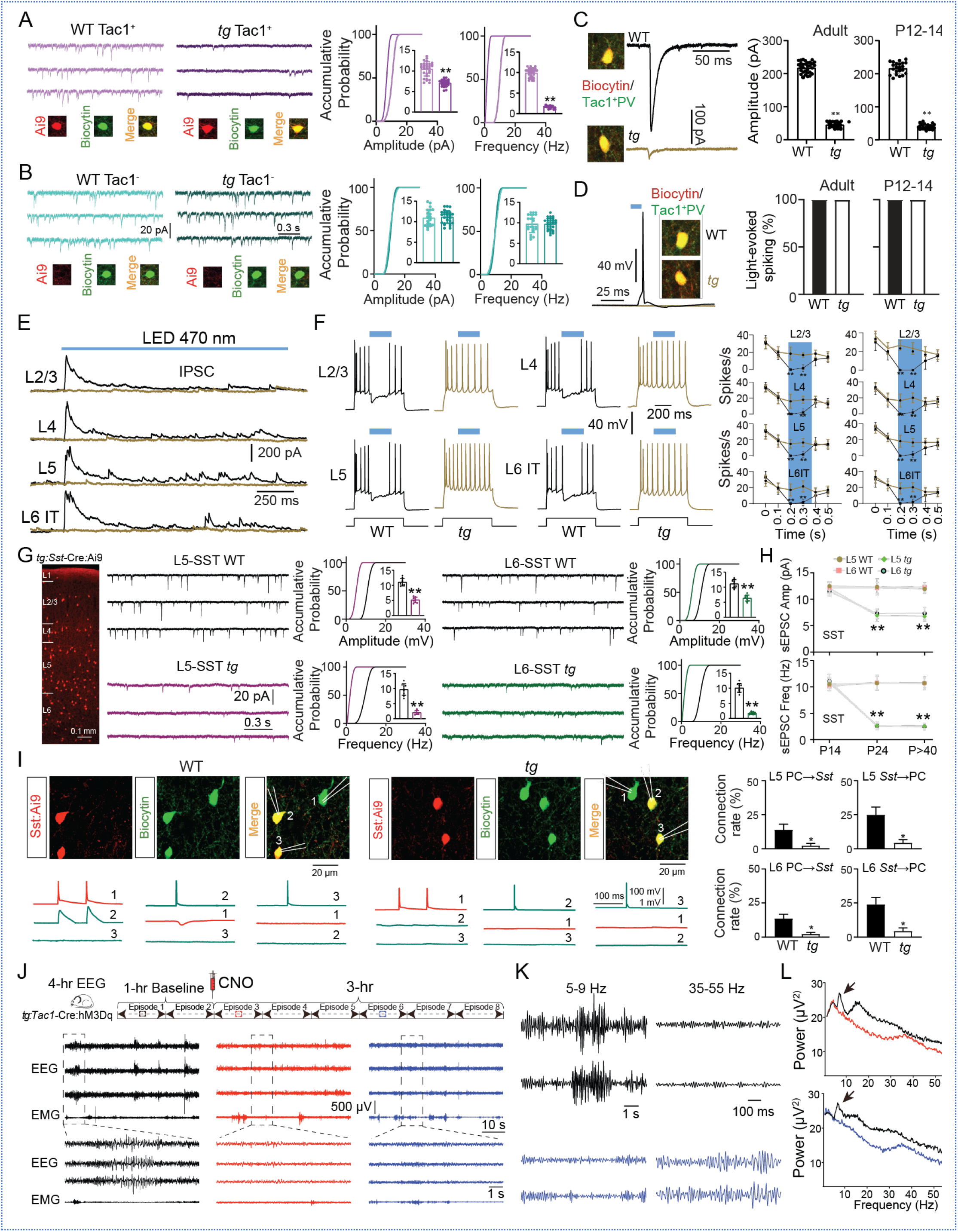
| Primary synaptic disruption in Tac1^+^ interneurons and secondary impairment in *Sst* interneurons of tottering mice. **a,** Representative traces of spontaneous excitatory postsynaptic currents (sEPSCs) recorded from Tac1^+^ *Pvalb* interneurons in WT and tottering (*tg*) mice. Recorded cells were confirmed by biocytin filling and Tac1 immunostaining. Right, cumulative probability plots and inset bar graphs show a significant reduction in sEPSC amplitude and frequency in Tac1^+^ interneurons from *tg* mice compared with WT mice. WT, n = 32 cells; *tg*, n = 27 cells; \*\**p* < 0.01. **b,** Representative traces of sEPSCs recorded from Tac1^−^ interneurons in WT and *tg* mice. Right, cumulative probability plots and inset bar graphs show no significant differences in sEPSC amplitude or frequency between WT and *tg* Tac1^−^ interneurons. WT, n = 23 cells; *tg*, n = 29 cells. **c,** Optogenetic stimulation of L6 corticothalamic (CT) neurons in *Ntsr1*-Cre:Ai32 mice evoked robust excitatory postsynaptic currents (oEPSCs) in Tac1^+^ interneurons in WT mice, but these responses were markedly reduced in *tg* mice. Right, quantification of oEPSC amplitude in Tac1^+^ interneurons. For adult mice: WT, n = 31 cells from 12 mice; *tg*, n = 30 cells from 11 mice. For P12-24 mice: WT, n = 22 cells from 8 mice; *tg*, n = 24 cells from 10 mice. \*\**p* < 0.01. **d,** Left, optogenetic activation of L6 CT neurons triggered action potential firing in Tac1^+^ interneurons in WT mice, but this response was largely absent in *tg* mice. Right, quantification of the proportion of responsive cells across developmental stages (adult and P12–24), showing robust responses in WT mice but loss of responsiveness in *tg* mice. **e,** Representative optogenetically evoked inhibitory postsynaptic currents (oIPSCs) recorded from pyramidal cells across cortical layers during 1.5 s blue-light stimulation of L6 CT neurons. Robust oIPSCs were observed in WT mice but were largely absent in *tg* mice. Representative traces were obtained from 12 mice. **f,** Left: Optogenetic stimulation of L6 CT neurons suppressed action potential firing in pyramidal cells during a +140 pA depolarizing current injection; blue light was delivered for 200 ms during the current step. This suppression was robust in WT mice but absent in *tg* mice. Right: Quantification of light-induced firing inhibition in pyramidal cells across cortical layers in juvenile (P12-24) and adult mice. For P12-24 mice: L2/3, n = 6 cells from 5 WT mice and n = 6 cells from 6 *tg* mice; L4, n = 5 cells from 5 WT mice and n = 5 cells from 5 *tg* mice; L5, n = 6 cells from 5 WT mice and n = 6 cells from 6 *tg* mice; L6, n = 7 cells from 6 WT mice and n = 7 cells from 6 *tg* mice. For adult mice: L2/3, n = 6 cells from 6 WT mice and n = 6 cells from 5 *tg* mice; L4, n = 6 cells from 6 WT mice and n = 6 cells from 6 *tg* mice; L5, n = 6 cells from 5 WT mice and n = 6 cells from 5 *tg* mice; L6, n = 7 cells from 5 WT mice and n = 7 cells from 6 *tg* mice. \*\**p* < 0.01. **g,** Left, representative fluorescence image showing *Sst* interneurons in the cortex of *Sst*-Cre:Ai9 mice. Representative sEPSC traces from L5 *Sst* interneurons in WT and *tg* mice are shown, together with cumulative probability plots and inset bar graphs demonstrating significantly reduced sEPSC amplitude and frequency in *tg* mice. WT, n = 10 cells; *tg*, n = 8 cells; *p* < 0.01. Right, representative sEPSC traces from L6 *Sst* interneurons in WT and *tg* mice, with cumulative probability plots and inset bar graphs showing significantly reduced sEPSC amplitude and frequency in tg mice. WT, n = 10 cells; *tg*, n = 9 cells; *p* < 0.01. **h,** Quantification of sEPSC amplitude and frequency in L5 and L6 *Sst* interneurons across developmental stages (P14, P24, and P > 40) in WT and tg mice. For L5 *Sst* interneurons: P14, n = 9 WT and n = 8 *tg*; P24, n = 8 WT and n = 9 *tg*; P > 40, n = 10 WT and n = 10 *tg*. For L6 *Sst* interneurons: P14, n = 8 WT and n = 10 *tg*; P24, n = 10 WT and n = 8 *tg*; P > 40, n = 9 WT and n = 9 *tg*. Student’s t-test was used for WT versus *tg* comparisons at each developmental stage. \*\**p* < 0.01. **I,** Left: Biocytin-filled patched cells from WT and *tg* mice labeled with *Sst*:Ai9 were used for post hoc morphological confirmation of *Sst* interneurons. Bottom: Representative traces from simultaneous recordings show principal cells (PCs) and *Sst* interneurons. Right: Summary of connection probabilities for PC→*Sst* and *Sst*→PC pairs in L5 and L6. In both layers, connection probabilities for PC→*Sst* and *Sst*→PC were significantly reduced in *tg* mice compared with WT mice. For L5 PC→*Sst*, WT = 9/66 and *tg* = 2/68; for L6 PC→*Sst*, WT = 15/110 and *tg* = 3/119; for *Sst*→L5 PC, WT = 13/54 and *tg* = 3/68; for *Sst*→L6 PC, WT = 16/65 and *tg* = 3/67. \**p* < 0.05. **j,** Schematic of the experimental design for EEG and EMG recordings in *tg*:*Tac1*-Cre:hM3Dq mice. Representative baseline and post-activation EEG traces show disappearance of absence seizure activity after CNO administration. Expanded traces highlight representative spike-wave discharges before (red) and after (blue) chemogenetic activation. **k,** Frequency-band analysis of EEG traces in *tg*:*Tac1*-Cre:hM3Dq mice shows dominant 5-9 Hz absence seizure activity before chemogenetic activation, which was largely eliminated after activation. In parallel, gamma-band activity (35-55 Hz) increased after activation, consistent with effective suppression of seizure activity by chemogenetic activation of Tac1^+^ interneurons. **l,** Power spectrum analysis of EEG recordings in *tg*:*Tac1*-Cre:hM3Dq mice shows a marked reduction in 5-9 Hz power after chemogenetic activation (arrows), accompanied by an increase in gamma-band power (35-55 Hz).

To investigate if *tg* mice exhibit the same additional deficit on *Sst* interneurons as *stg*, we performed whole cell recordings of genetically labeled *Sst* interneurons across cortical layers in *tg* mice (Fig. 6g). Similar to the *stg* recordings, deep-layer (L5/6) *Sst* interneurons in *tg* mice exhibited a marked reduction in sEPSC frequency and amplitude compared to WT controls (Fig. 6g), whereas those in superficial layers (L2-4) remained little affected by the *Cacna1a* mutation (Extended Data Fig. 8f). In alignment with these results, multi-patch recordings revealed a significant reduction in afferent connectivity from L5/6 PCs onto *Sst* interneurons (Fig. 6i). Unlike Tac1^+^ interneurons, these *Sst* interneurons also displayed diminished inhibitory output onto local PCs (Fig. 6i) and a pronounced decrease in intrinsic excitability (Extended Data Fig. 8g and Table 1). Crucially, developmental time-course analysis revealed that these physiological impairments in *tg* mice were absent prior to the seizure onset (P < 14), and only became apparent at the initial emergence of seizures (P18-P20) and persisted long into adulthood (P>40) (Fig. 6h)^51,76,77^. Together these data indicate a secondary, functional deficit of deep-layer *Sst* interneurons that mirrors the *stg* circuit phenotype.

To further test if the *tg* mice share the same functional cortical circuit defects as *stg*, we investigated whether rescuing these defects could also mitigate absence seizures in *tg*. To this end, we crossed *Tac1*-Cre:Gq mice with *tg* mice to obtain *tg*:*Tac1*-Cre:Gq mice. We validated our DREADD approach as in *stg* mice (Extended Data Fig. 6a). Baseline electroencephalography (EEG) recordings in these mice prior to CNO administration again showed characteristic spike-wave patterns, primarily occurring within the 5-9 Hz range (Fig. 6j). As in *stg* mice, a single intraperitoneal injection of 0.1 mg/kg CNO eliminated absence seizures for approximately 3 hours (Fig. 6j, k) in all *tg* mice (Extended Data Fig. 8h, I; n = 6). In contrast, no post-injection change in seizure activity was detected in control mice (Extended Data Fig. 8i). Power spectrum analysis showed a significant decrease in waveform activity within the 5-9 Hz frequency band following CNO administration, accompanied by a significant increase in gamma oscillations (Fig. 6k, l).

## Discussion

Despite extensive analysis of pathogenic thalamic remodeling in monogenic models of CAE^15,16,51,78–80^, it has remained largely unclear whether, and how, CAE gene mutations impair cortical circuits that ultimately give rise to the SWD underlying absence epilepsy. This limitation represents a critical gap in our understanding of the disorder and hinders our ability to develop therapies that directly target the precise cellular or neural circuit abnormalities mediating the pharmacoresistance and cognitive comorbidities which affect a majority of cases^7^. We examined two well-established mouse absence epilepsy models to test whether mutations in their genes converge on shared functional cortical circuit deficits mediating the stereotyped SWD episodes. By extensively mapping functional cortical microcircuitry across genotype and developmental stage, we uncovered a stereotyped, chronological sequence of circuit pathogenesis: a serial dysfunction of translaminar inhibitory circuits initiating in deep-layer *Pvalb* interneurons that sequentially progresses to *Sst* interneurons. Our findings provide evidence for a key emerging theme in monogenic epileptogenesis, namely the importance of early postnatal circuit remodeling defects that underlie the progressive lowering of seizure threshold until the appearance of electrographic and clinical seizures. In these absence seizure models, this cascade occurs at both nodes of the thalamocortical circuit, where cortical microcircuit remodeling may act in concert with the remodeling of thalamic neurons for seizure generation^15,51,78,79^. Ultimately, the intersection of these genetic lesions and neurodevelopment yields a dynamically stereotyped disease pathophysiology^30^.

### Cortical circuit mechanisms underlying absence seizures in *stg*

Stargazer (*stg*/*stg*) mice harbor a *Cacng2* mutation that impairs AMPA receptor trafficking^48,73,81^, leading to deficient excitatory synaptic transmission^34^. Mutations in human *CACNG2*^80^ and the closely related *CACNG3*^81^ genes have been identified in human absence epilepsy and the electrophysiological, pharmacological, and behavioral similarities between absence seizures in human and *stg* mice^8,82–87^ validate this mutant as a robust model for dissecting epileptic circuitry. By leveraging the restricted expression of *Cacng2* in cortical interneurons, we uncovered a translaminar cortical inhibitory microcircuit connecting layer 6 corticothalamic (L6 CT) neurons to a novel *Pvalb* interneuron subtype, Tac1^+^ interneurons. While classically known for their thalamic projections, L6 CT neurons also have extensive intracortical projections, particularly targeting a deep-layer *Pvalb* interneuron subpopulation with extensive translaminar axons. L6 CT neurons mediate sensory gain control primarily by recruiting this specific intracortical *Pvalb* population^43–45^. Here, we establish the molecular identity of this specific *Pvalb* subpopulation as Tac1^+^ interneurons. Because *Cacng2* expression is exquisitely restricted to these Tac1^+^ interneurons, the *stg* mutation selectively dismantles this L6 CT-Tac1^+^ translaminar inhibitory microcircuit. Consequently, the disruption of L6 CT-driven cortical inhibitory circuitry represents a primary defect resulting directly from the *stg* gene mutation^88^.

While the disrupted microcircuit is necessary for epileptogenesis, this primary defect exists prior to seizure onset. Absence seizures are not manifested at birth despite the congenital nature of the mutation. Clinically, childhood absence epilepsy (CAE) typically presents between 3 and 8 years of age, and mouse models like *stargazer* (*stg*) exhibit a stereotypical seizure-free latent period. This temporal gap indicates that the mature epileptic phenotype requires a progressive, developmental sequence of cumulative circuit remodeling^30^. Thus, epileptogenesis emerges not directly from the primary mutation, but from maladaptive circuit plasticity driven by subclinical, aberrant network activity. While pathogenic remodeling in the thalamus is heavily documented in absence seizures^15,16,51,78–80^, our findings demonstrate that concomitant maladaptive plasticity in the cortex may be equally important. We identified a delayed, non-cell-autonomous functional impairment of deep-layer *Sst* interneurons, particularly those Martinotti cells (MCs), which lack *Cacng2* expression. In normal neocortex, layer 5 *Sst* MCs project to layer 1 (L1) to innervate the apical dendrites of pyramidal cells (PCs), providing the critical lateral disynaptic inhibition necessary to regulate columnar excitability and prevent hypersynchrony during high-frequency PC firing^89^. In *stg* mice, the failed recruitment of these *Sst* interneurons and a profound reduction of their L1 projections narrowly coincides with seizure onset. This suggests that the loss of this specific apical inhibitory tone acts as the critical tipping point for epilepsy, aligning with broader evidence that *Sst* interneuron dysfunction is a core pathogenic driver in various epilepsies featuring absence seizures as part of a more complex phenotype^90,91^. Despite overwhelming evidence, to definitively establish the causal necessity of this secondary defect in SWD generation will require development of early interceptive strategies that preserve both axonal integrity and intrinsic excitability prior to seizure onset, such as cell-type-specific genetic or molecular rescue strategies informed by single-cell transcriptomics.

Mechanistically, this secondary change reflects a maladaptive activity-dependent plasticity as a response to the primary genetic defect. The initial mutation causes a cell-autonomous disconnection of Tac1^+^ *Pvalb* interneurons from the cortical network, establishing a “hypo-GABAergic” state that severely impairs L6-mediated sensory gain control. In response to this translaminar loss of perisomatic inhibition, the cortical network dynamically retunes its connectivity by reducing excitatory drive from PCs and stripping *Sst* interneurons of their essential wiring via activity-dependent plasticity^92^. The obligatory nature of this maladaptive reaction is underscored by our *in vivo* manipulations that replicated the defect in healthy cortex: chemogenetic inhibition of translaminar Tac1^+^ *Pvalb* interneurons in wild-type mice required at least four days of continuous intervention to precipitate *Sst* interneuron dysfunction and absence seizures. Therefore, rather than being a direct consequence of the initial lesion, the full expression of the epileptic phenotype may depend upon a highly specific functional impairment within the *Sst* interneuron network.

Our findings support the view that the deep layers of the somatosensory cortex (S1) constitute a critical cortical initiation node, demonstrating that localized cellular pathology within this region is sufficient to initiate SW discharges^18–20,93–97^. Previous studies have also documented thalamic abnormalities in *stg* mice, including impaired excitatory drive onto inhibitory neurons of the thalamic reticular nucleus (TRN)^16,48,98^. However, the broad cortico-thalamic expression of genetic tools used in many previous studies has largely failed to isolate cortical from thalamic contributions^98,99^. Isolating distinct cortical from thalamic mechanisms has therefore remained a longstanding challenge in absence epilepsy. In this study, we overcame this challenge by leveraging a cortex-restricted *Tac1*-Cre line, enabling precise manipulation of S1 L5/6 Tac1^+^ interneurons without affecting the TRN. Strikingly, chemogenetic activation of just Tac1^+^ *Pvalb* interneurons in S1 (60% of L5/6 *Pvalb* interneurons) was sufficient to abolish seizure occurrence in *stg* mice, whereas their suppression in wild-type mice induced SWDs phenocopies of the genetic models. Collectively, these data demonstrate that isolated pathophysiological dysfunction within this deep-layer cortical circuit are sufficient to drive absence epilepsy, regardless of any changes within thalamic cell populations^15,16,51,78–80^.

In addition to seizures, the disruption of L6-driven cortical inhibition identified in somatosensory cortex would be expected to compromise sensory processing, linking to well-documented interictal deficits reported in both patients and mouse models including impaired sensory coding, reduced stimulus selectivity, and altered tactile processing^26,100,101^. These findings also align well with the clinical presentation of CAE, where sensory modulation disorders and attention deficits are highly prevalent^4,101^. Because layer 6 neurons and sensory gain control are essential for attention processing^102–104^, the L6-driven gain control deficits in *stg* mice offer a compelling mechanistic link between absence seizures and their common cognitive comorbidities.

### Shared circuit mechanisms across genetic lesions

The list of causative gene mutations for absence epilepsy continues to expand across a functionally diverse spectrum of biology^8,9,105^. It remains enigmatic how such a wide array of functionally distinct genes converges to produce this highly stereotypical electroclinical phenotype. One prevailing hypothesis is that distinct gene mutations converge onto the same functional circuit deficits, which in turn give rise to identical EEG patterns and clinical phenotypes. To test this hypothesis, we examined another model of absence seizures, the *tottering* (*tg*) mouse, which harbor mutations in *Cacna1a* encoding the P/Q-type calcium channel alpha subunit^35^. In humans, at least ten loss of function *Cacna1a* mutations (distinct from those linked to ataxia and familial hemiplegic migraine) have been identified in patients with absence epilepsy^106^, strongly supporting *tg* as a translational model for dissecting epileptic circuitry. We found that the developmental loss of *Cacna1a*, which impairs presynaptic glutamate release, causes synaptic defects in the same cortical microcircuit affected by the loss of the AMPA receptor trafficking gene in *stg* mice. Despite the widespread expression of P/Q-type calcium channels across neural circuits, a critically vulnerable locus resides within cortical layer 6 corticothalamic (L6 CT) neurons, consistent with findings that the specific deletion of *Cacna1a* from L6 CT neurons is sufficient to drive absence seizures^97^. While parallel primary defects in other circuits remain to be fully characterized, such as L6 CT projections delivering P/Q-deficient synaptic input to thalamic circuits to influence both nodes of the SWD network^97,107^, our results with cortex-specific manipulations suggest that defects within the cortical node alone is sufficient for inducing seizure generation.

The primary cortical synaptic defect in *tg* mice is thus functionally overlapping with that in *stg* mice, despite operating through distinct mechanisms, presynaptically in *tg* and postsynaptically in *stg*. This shared primary defect would be expected to trigger the same delayed functional reorganization of cortical networks. Indeed, we identified similar secondary circuit deficits centered on deep-layer *Sst* interneurons in both models. These observations not only establish a convergent, secondary circuit mechanism underlying absence seizures but also highlight specific cellular vulnerabilities amenable to targeted phenotypic rescue. Crucially, we demonstrated that this chronological sequence of circuit pathogenesis can be exogenously induced in adult wild -type mice. Transient suppression of Tac1^+^ interneurons for two weeks, mimicking synaptic disruption observed in the genetic models, was sufficient to trigger both absence seizures and enduring, *Sst*-centered circuit reorganization. This manipulation effectively recapitulated the core pathological architecture shared by the genetic models. Together, our findings provide a mechanistic framework for how distinct genetic mutations converge to generate identical seizure phenotypes: disparate subcellular disruptions yield a unified primary defect, which subsequently drives a stereotypic circuit-level pathology. Determining whether this convergent downstream pathway extends to other models of absence epilepsy, such as those driven by GluR4^108,109^ and PICK1 mutations^110^, as well as polygenic models of absence epilepsy^99^, represents a critical next step for future investigations. If these stereotypic circuit deficits prove to be a universal feature of experimental and human absence epilepsy, it would offer an important strategic shift in the therapeutic paradigm, redirecting focus from correcting individual genetic lesions toward resolving emergent circuit imbalances using next-generation precision therapeutics for intractable epilepsy such as interneuron grafting, gene therapies, and cell type-specific pharmacology.

### In summary

Our findings reveal a sequential cortical circuit pathology in which an early functional deficit of deep-layer Tac1^+^ *Pvalb* interneurons precedes the delayed functional and structural impairment of deep-layer *Sst* interneurons associated with seizure emergence. This temporal separation suggests that epileptogenesis involves progressive secondary circuit remodeling rather than an immediate consequence of the initial defect alone. The delayed *Sst* interneuron abnormalities may therefore represent a potential temporal window for intervention. Approaches that preserve or restore the function of deep-layer Tac1^+^ *Pvalb* and *Sst* interneuron circuits, or prevent the secondary remodeling that links these abnormalities, may provide a useful framework for developing strategies to limit seizure development across CAE syndromes.

## Acknowledgments

We thank members of the Jiang and Noebels labs for technical assistance, discussion, and comments on the manuscript. Research reported in this publication is supported by research grants R01 MH122169 (to X.J.), R01 MH120404 (to X.J.), R01 NS110767 (to X.J., J.N.) and R01 NS29709 (to JLN) from the National Institutes of Health. Research reported in this publication was also supported by the Main Street America Fund, a Shared Instrumentation grant from the NIH (1S10OD016167), and by the Eunice Kennedy Shriver National Institute of Child Health & Human Development of the National Institutes of Health under Award Number P50HD103555 for the use of the Microscopy Core facility, the RNA In Situ Hybridization Core facility, and In Vivo Neurophysiology Core facility at Baylor College of Medicine. We thank Dr. Andreas Tolias for helping with the initial data collection.

## Author contributions

S.S., N.L., D.T., C.L., J.T., and X.J. performed the experiments. S.S. and X.J. analyzed the data. X.J., J.N., and A.M. designed and supervised all experiments and data analyses. S.S. and X.J. prepared the manuscript with input from all co-authors.

## Data availability

Source data underlying the figures and tables are provided with this paper. The previously published single-cell, single-nucleus RNA-sequencing and Patch-seq datasets analyzed in this study are publicly available through the Allen Brain Atlas (Tasic et al. and Yao et al.; Tasic et al., GEO accession GSE115746), NeMO Archive (Gouwens et al.), Gene Expression Omnibus (GSE153164) and Single Cell Portal (SCP1290) (Di Bella et al.), and UCSC Cell Browser (Anderson et al.), as detailed in the Methods. The remaining data generated in this study are available from the corresponding author upon reasonable request.

## Code availability

Custom analysis code used for the EEG, single-cell RNA-sequencing and Patch-seq analyses in this study is publicly available at GitHub: https://github.com/siyuansong001/absence-seizure-analysis.

## Methods

### Animals

All experiments were conducted in adherence to the guidelines set forth by the Institutional Animal Care and Use Committee at Baylor College of Medicine. Animals were housed in a standard animal facility with a 12-hour light/dark cycle (06:00 to 18:00), maintained at 20-22°C with a humidity range of 30%-70%. For slice electrophysiology, we utilized wild-type (WT) and transgenic mice of both sexes, all on a C57BL/6J background. Our study spanned a broad range of animal ages, from postnatal (P)12 to beyond P50, to account for the varying incidence of absence seizures at different developmental stages. Additionally, we examined *Cacng2* expression at P4. Stargazer (*stg*) and tottering (*tg*) mice served as established absence seizure models ^33,35,52,111^. To systematically target distinct neural populations for morphological, electrophysiological, optogenetic, and chemogenetic characterization, we employed multiple Cre-driver and reporter lines. For Tac1^+^ *Pvalb* interneurons, *Tac1*-Cre mice were crossed with Ai9 (n=32), Ai32 (n=12), or hM4Di (n=38) reporters; *Tac1*-Cre:Ai9 mice were further crossed with *stg* (n=16) and *tg* (n=15) lines to study these cells within the seizure models. To target layer 5/6 *Pvalb* interneurons, *Kcnc2*-Cre or *Tac1*-Cre mice (The Jackson Laboratory) were crossed with hM3Dq reporters and subsequently with *stg* mice (*stg*:*Kcnc2*-Cre:hM3Dq, n=18; *stg*:*Tac1*-Cre:hM3Dq, n=16). Layer 6 corticothalamic (CT) neurons were targeted via *Ntsr1*-Cre crosses with Ai9 (n=8) or Ai32 (n=38) reporters, with the latter further crossed to generate *stg*:*Ntsr1*-Cre:Ai32 (n=21) and *tg*:*Ntsr1*-Cre:Ai32 (n=22) models. Additional interneuron subtypes in the *stg* model were targeted by crossing *stg* mice with *Sst*-Cre:Ai9 (n=52), *Vip*-Cre:Ai9 (n=22), and *Npy*-hrGFP (n=18) lines (for neurogliaform cells), while *Sst* interneurons in the *tg* model were labeled using *tg*:*Sst*-Cre:Ai9 mice (n=42). Finally, to examine the axonal distribution of layer 5/6 *Sst* interneurons in the *stg* model, we crossed *stg* with X98-EGFP mice (*stg*:X98-EGFP, n=8), a line generously provided by Dr. Aric Agmon at West Virginia University ^57^.

### Slice preparation and electrophysiology

Brain slices were prepared following previously established methods ^112^. Briefly, mice were deeply sedated using 3% isoflurane before decapitation. Brains were quickly extracted and immersed in chilled, oxygenated N-methyl-D-glucamine (NMDG) solution. Brain regions containing the somatosensory cortex (S1) were sectioned parasagittally into 300-µm-thick slices using a Leica microtome (VT1200). Slices were placed in oxygenated NMDG solution at 33 ± 0.5°C for 10 minutes, followed by incubation in artificial cerebrospinal fluid (ACSF:125 mM NaCl, 2.5 mM KCl, 1.25 mM NaH_2_PO_4_, 25 mM NaHCO_3_, 1 mM MgCl_2_, 25 mM glucose, and 2 mM CaCl_2_ at pH 7.4) at the same temperature for 40-60 minutes before electrophysiological recordings.

Whole-cell patch-clamp recordings (single-cell or multi-cell recordings) were performed on S1 neurons as described previously ^112^. We used borosilicate glass pipettes (with resistances of 3-5 MΩ) crafted using a micropipette puller (P-1000, Sutter) and filled with a potassium-based intracellular solution (120 mM potassium gluconate, 10 mM HEPES, 4 mM KCl, 4 mM MgATP, 0.3 mM Na_3_GTP, 10 mM sodium phosphocreatine, and 0.5% biocytin, pH 7.25), or the cesium-based internal solution (115 mM cesium methanesulfonate, 20 mM CsCl, 10 mM HEPES, 2.5 mM MgCl_2_, 4 mM Na_2_ATP, 0.4 mM Na_3_GTP, 10 mM sodium phosphocreatine, 0.6 mM EGTA, pH 7.25). Recordings were conducted using Quadro EPC 10 amplifiers (HEKA Electronics, Lambrecht, Germany), and PatchMaster (HEKA) and custom-written MATLAB-based programs (MathWorks) were used to operate the recording system and perform online and offline data analysis. *Pvalb* interneurons were identified by their distinct firing pattern (fast-spiking) and unique morphology in *stg* mice, as it is not feasible to create a transgenic *stg* mutant expressing a *Pvalb*-driven reporter to label *Pvalb* interneurons due to the tight linkage of the *Cacng2* and *Pvalb* loci on chromosome 15 ^27^. *Sst* interneurons were identified through fluorescence using *Sst*-Cre mice or through *post hoc* immunostaining for Sst following slice recordings (see below). Cells were only accepted for analysis if the initial series resistance was less than 40 MΩ and did not change by more than 20% during the recording period. The series resistance was compensated by at least 50% in voltage-clamp mode. No correction was made for the liquid junction potential.

Simultaneous whole-cell recordings from up to eight neurons were performed using two Quadro EPC 10 amplifiers. To identify synaptic connections, current pulses were injected into presynaptic neurons (2 nA for 2 ms at 0.01-0.1 Hz) to evoke AP while the membrane potential of other simultaneously recorded neurons w as monitored to detect unitary inhibitory or excitatory postsynaptic potentials (uI(E)PSPs). The uIPSPs were measured while the membrane potentials of the putative postsynaptic cells were held at -60 ± 3 mV, whereas uEPSPs were measured while the membrane potentials of the putative postsynaptic cells were held at - 70 ± 3 mV. To estimate the connection probability between two cell types, we used the equation as follows: Connectivity rate (%) = total identified synaptic connections/the total potential connections tested × 100%^113^. The spike threshold and firing pattern in response to sustained depolarizing currents were recorded for each neuron by injecting increasing current steps (+10 pA) every second. Spontaneous excitatory synaptic events were recorded at -70 mV for a minimum duration of 30 seconds. To examine NMDA receptor-mediated excitatory postsynaptic currents (NMDA-EPSCs), neurons were recorded with the cesium-based solution and clamped at +40 mV to release the blocking effects of magnesium ions^114^. Neurons were then fixed for *post hoc* morphological recovery with biocytin staining as well as immunostaining.

To explore the dynamics of inhibitory synaptic transmission in tottering mice, five consecutive presynaptic stimuli (2 nA, 2 ms) were delivered at 100 ms intervals to *Pvalb* interneurons. The resulting unitary inhibitory postsynaptic potential (uIPSP) amplitudes were recorded in postsynaptic principal cells (PCs) held at −60 ± 3 mV. Changes in uIPSP amplitudes across the five stimuli were analyzed to assess synaptic dynamics.

### Optogenetics and chemogenetics in brain slices

To activate ChR2-expressing neurons in brain slices (*Tac1*-Cre:Ai32 and *Ntsr1*-Cre:Ai32 mice, as well as the corresponding crosses with stargazer (*stg*) or tottering (*tg)* mice where indicated), 470 nm LED blue light (intensity ∼6 mW/mm²) for varying durations was delivered through a 40× objective len on the microscope, generating a light spot with a spatial spread of ∼0.25 mm², covering half of the cortical layers in the recorded slices. Neurons were considered responsive if the latency of the response (EPSC, EPSP, IPSC, or IPSP) was less than 7 ms, and the response amplitude exceeded three times the standard deviation of the baseline.

Optogenetically evoked inhibitory postsynaptic currents (oIPSCs) were recorded using cesium-based intracellular solutions, with the membrane potentials clamped at +7 mV, while optogenetically evoked excitatory postsynaptic currents (oEPSCs) were recorded with the membrane potentials clamped at -80 mV. Each recorded neuron was identified through fluorescence or through *post hoc* immunostaining with the appropriate markers following slice recordings (see below). Spikes were induced in pyramidal neurons from layers 2/3, 4, 5, and L6 intratelencephalic (IT) neurons using +140 pA current injections lasting for 600 ms. 200 ms light pulses were delivered 200 ms after current onset, and the resulting suppression of firing was quantified as the change in spikes per second (Hz).

To verify DREADDs expression in targeted neurons *in vitro*, current-clamp recordings were performed on these neurons, and their firing rates and membrane potentials were recorded and compared before and after bath application of 10 μM clozapine-N-oxide (CNO; Hello Bio).

### Morphological recovery after slice recording

Morphological recovery of recorded neurons and light microscopic examination of their morphology were carried out following previously described method_s_^112^. In brief, upon the completion of whole- cell recordings, the slices were immediately fixed in 2.5% glutaraldehyde/4% paraformaldehyde in 0.1M PB at 4 °C for 10-14 days and then processed with an avidin-biotin-peroxidase method with an ABC kit (Vector Laboratories, PK-4000) to reveal cell morphology. Recovered neurons were examined and reconstructed using a 100× oil immersion objective lens and camera lucida system (Neurolucida, MicroBrightField). For morphological feature extraction, only cells with intact dendritic and axonal structures were included. We took special care to exclude reconstructions of the neurons that showed any signs of axonal damage (due to the slicing procedure).

### RNA fluorescence *in situ* hybridization

Wild-type or transgenic mice at different stages were deeply anesthetized with 3% isoflurane and decapitated immediately, and their brains were immediately removed from the skull and embedded in optimal cutting temperature (OCT, Sakura) compound on dry ice. The brain blocks were then sagittally sectioned into 25 μm thick slices. RNA fluorescence in situ hybridization (FISH) was performed on sections containing S1 or reticular thalamic nucleus (RTN) at the RNA In Situ Hybridization Core at Baylor College of Medicine, using a

previously described in situ hybridization method with slight modifications^115^. Briefly, digoxigenin (DIG) and fluorescein isothiocyanate (FITC) labeled mRNA antisense probes were first synthesized from reverse-transcribed mouse cDNA. Following fixation and acetylation, the two probes were then simultaneously hybridized to the brain sections. Following a series of washing and blocking steps, DIG-labeled probes were visualized with tyramide-Cy3 Plus, and FITC-labeled probes were visualized with tyramide-FITC Plus.

Sections were counterstained with 4’,6-diamidino-2-phenylindole (DAPI) to delineate the S1 cortical layers. The brain sections were then mounted with Prolong Diamond (Invitrogen). Images were captured at 20× magnification using LSM 710 or 880 confocal microscope (Zeiss). ImageJ was used to quantify the mean density of expression and examine gene expression levels and co-labeling rates across cortical layers.

Fluorescence intensity was quantified using ImageJ software. For each region of interest (ROI), the mean fluorescence intensity was calculated as the sum of all pixel intensity values within the ROI divided by the number of pixels. The results are presented in arbitrary units (a.u.), as is standard for relative fluorescence quantification. Mean = (Sum of pixel intensities in the region of interest) / (Number of pixels in the region of interest). Typical FISH images were exported for the final layout with ZEN 2.3 (Zeiss). Primer sequences for the probes were derived from Allen Brain Atlas (http://www.brain-map.org). The sequences of the probes used for FISH are listed in Extended Data Table 2.

### Immunofluorescence staining

To identify Tac1 co-localization with stargazin, *Tac1*-Cre:Ai9 mice at P4 or adult stages (P > 60) were euthanized with 3% isoflurane in fume hood and then perfused transcardially with 20mL PBS followed by 20mL of 4% paraformaldehyde (PFA) in 0.1M PB. The brain was removed, post-fixed in 4% PFA for 1 day, and then sagittally sectioned into 50 μm thick slices in PBS. The slices were washed three times in PBS before blocking with 10% goat serum and 1% Triton X in PBS for 1 hour. The slices were incubated first with mouse anti-stargazin antibody (1:10, NeuroMab, 75-242) followed by incubation with goat anti-mouse Alexa Fluor 488 secondary antibody (1:200, Thermo Fisher Scientific, A-11001). To examine GluR4 receptors in different conditions, 50 μm thick brain slices were incubated first with rabbit anti-GluR4 antibody (1:200, EMD Millipore, AB1508) and mouse anti-PSD-95 antibody (1:400, Abcam, ab13552) as primary antibodies, and then incubated with goat anti-mouse Alexa Fluor 633 (Thermo Fisher Scientific, A-21052) or goat anti-mouse Alexa Fluor 405 (Thermo Fisher Scientific, A-31553), together with goat anti-rabbit Alexa Fluor 488 (1:200, Thermo Fisher Scientific, A-11034), as secondary antibodies for at least 1 h. Slices were then mounted with the mounting medium (Abcam, ab104139).

To stain Tac1 or somatostatin (Sst) in recorded neurons (biocytin-filled), brain slices were immediately fixed after whole-cell recording with 4% PFA (Sigma-Aldrich, P6148) for 12-24 h at 4°C, washed three times with PBS, and incubated overnight at room temperature with primary antibodies, rabbit anti-Tac1 (1:200, Proteintech, 13839-1-AP) or rabbit anti-Sst (1:200, ImmunoSTAR, 20067). The slices were subsequently washed with PBS and incubated for 2 hours at room temperature with streptavidin conjugated to Alexa Fluor 568 (Thermo Fisher Scientific, S11226) or Alexa Fluor 430 (Thermo Fisher Scientific, S11237) (to visualize biocytin in recorded neuron) together with one of the following secondary antibodies (1:200), depending on the experimental requirements: goat anti-rabbit Alexa Fluor 555 (Thermo Fisher Scientific, A-21428), goat anti-rabbit Alexa Fluor 488 (Thermo Fisher Scientific, A-11034), or goat anti-rabbit Alexa Fluor 647 (Thermo Fisher Scientific, A-21245). Images were captured at 10× or 20× magnification using LSM 710 confocal microscope (Zeiss).

### EEG recording

Mice were sedated using isoflurane (1.5%-3% in oxygen, delivered via a Matrox Vip300 ventilator), and electrodes attached to a connector were surgically implanted into the subdural space over the frontal and parietal cortices ^116^ and/or striatum to enable EEG recording. Both sexes of mice underwent simultaneous video-EEG and behavioral monitoring (LabChart 8.0, ADI Systems). EEG monitoring spanned 4 hours, generally started from 11am, and mice were able to move freely within the testing enclosure. Mice received a single intraperitoneal injection of clozapine-N-oxide (CNO; i.p., 0.1 mg/kg) after one hour of EEG recordings.

To chronically suppress Tac1^+^ *Pvalb* interneurons, CNO was dissolved in drinking water (40 mg/L), and provided to mice *ad libitum* in light-protected bottles^61,62^. Epileptic activity was assessed by measuring the frequency and duration of spike-wave discharges (SWDs) lasting at least 0.5s over 6 hours of EEG recording per day starting from 11am. To control for diurnal variation, SWDs were quantified from a one-hour window (approximately 12:00 PM to 1:00 PM) within the 6-hour daily EEG recordings. A one-time intraperitoneal injection of ethosuximide (ETX, 200 mg/kg) was administered to evaluate its effectiveness in blocking these discharges. All EEG signals were amplified using a g.BSAMP biosignal amplifier (Austria), filtered through a 0.5-Hz high-pass and a 50-Hz low-pass filter (AD Instruments) and captured with LabChart 8.0 (AD Instruments). EEG files were first visualized using Sirenia Seizure 2.2.12 (Pinnacle Technology), and SWDs were manually identified if their amplitudes exceeded twice the root mean square value of the baseline EEG signal. Power spectral density analyses were conducted using custom mne-python cod_e_^117,118^, utilizing the plot_psd function (with frequency ranges from 0 to 55 Hz) for visualization.

For additional EEG recordings in the caudate-putamen (CPu), the recording probe was positioned in the CPu using stereotaxic coordinates (0.0, 2.4, 2.5), as previously described ^119^. The electrode placement was verified through post-mortem examination of the electrode tract.

### Motor test (Rotarod test and Foot-slip test)

Rotarod test was conducted on a rotarod apparatus (Ugo Basile_)_^120,121^. The rotation speed gradually increased from 4 to 40 rpm over the first 5 minutes and then remained constant at 40 rpm for an additional 5 minutes or until all mice fell off, whichever came first. Mice underwent four trials per day for two consecutive days, with a 30-minute inter-trial interval. Latency to fall was recorded, with “falling” defined as either physical detachment from the rod or the completion of two full passive rotations. Data were expressed as mean ± SD and were analyzed using a Student’s t-test.

The foot-slip test was conducted in a Plexiglass enclosure measuring 45 cm × 21 cm ^120,121^ and fitted with a grid of parallel rods on the floor. Each mouse was placed in the center of the grid, and the number of foot slips and the distance traveled were recorded over 10 minutes using a camera linked to ANY-maze software (Stoelting Co.). Foot-slip counts were normalized to total distance traveled. The data were expressed as mean ± SD and were analyzed using a Student’s t-test.

## QUANTIFICATION AND STATISTICAL ANALYSIS

### Reference datasets

#### Tasic Dataset (Tasic et al., 2018)^37^

This dataset comprises mouse cortical single-cell RNA sequencing (scRNA-seq) data. The raw expression matrix and corresponding cell type annotations were sourced from the Allen Brain Atlas (https://portal.brain-map.org/atlases-and-data/rnaseq) and the Gene Expression Omnibus (GEO) under accession number GSE115746.

#### Yao Dataset (Yao et al., 2021)^122^

For this dataset, we analyzed single-nucleus RNA sequencing (snRNA-seq) data specific to the adult mouse primary somatosensory cortex (S1). This subset was extracted from a broader dataset encompassing the entire adult mouse isocortex and hippocampal formation. The raw expression matrix and cell type annotations were accessed via the Allen Brain Atlas (https://portal.brain-map.org/atlases-and-data/rnaseq).

#### Gouwens Dataset (Gouwens et al., 2020)^38^

This dataset provides scRNA-seq data of mouse cortical GABAergic interneurons acquired via the Patch-seq technique, which captures transcriptomic, electrophysiological, and morphological features from the same cells. The data is accessible through the NeMO Archive (https://data.nemoarchive.org/other/AIBS/AIBS_patchseq/transcriptome/scell/SMARTseq/).

#### Di Bella Dataset (Di Bella et al., 2021)^39^

This dataset contains scRNA-seq data from the developing mammalian cortex at postnatal day 4 (P4). The raw expression matrix and cell type annotations were obtained from the GEO (accession number GSE153164) and the Single-Cell Portal (SCP1290).

#### Anderson Dataset (Anderson et al., 2023)^123^

The dataset encompasses multiple scRNA-seq and snRNA-seq datasets from embryonic and postnatal stages of mouse caudate-putamen (CPu). The dataset is accessible through the https://cells.ucsc.edu/?ds=mouse-striatal-dev.

### Data mining and computational analysis of single-cell transcriptomics datasets

To analyze the gene expression in the primary somatosensory cortex (S1), we obtained single-nucleus RNA-sequencing (snRNA-seq) data^36^, comprising a barcode table, gene table, and gene expression matrix. Data processing and visualization were performed using Scanpy^124^ according to established best practice_s_^125^.

Dimensionality reduction was achieved via principal component analysis (PCA) utilizing 50 principal components, followed by uniform manifold approximation and projection (UMAP) with 30 nearest neighbors. Cluster annotations relied on established Allen Institute marker gene lists^36,37^. Excitatory neurons were classified into 9 cell types including L2/3 IT, L4/5 IT, L5 IT, L5 ET, L5/6 NP, L6 IT, L6 CT, L6b, and L6 Car3. Inhibitory neurons, marked by *Gad1* and *Gad2* expression, were divided into 6 groups including *Pvalb*, *Sst*, *Sst Chodl*, *Vip*, *Sncg*, and *Lamp5* interneurons^36,37^. Expression of *Cacng2* in distinct cell clusters was visualized using UMAP plots.

To investigate *Cacng2* expression in patch-seq dataset, we examined *Cacng2* expression across annotated *Pvalb* MET subtypes in an established Patch-seq reference datase_t_^38^. Dot plots generated with Scanpy were used to visualize *Cacng2* expression across various Pvalb MET subtypes.

To investigate early developmental expression patterns, we analyzed an additional snRNA-seq dataset from the S1 region of postnatal day 4 (P4) mice^39^, again utilizing Scanpy for processing and clustering. Neuronal clusters were annotated using the following established marker genes: *Arpp21* and *Satb2* for excitatory neurons^39,126^; *Dlx2*, *Gad1*, and *Gad2* for inhibitory neurons; *Cxcl14, Vip,* and *Htr3a* for CGE-derived interneurons^127^; *Sst* for immature *Sst* interneurons^128,129^; *Mef2c, Igfbp4, Elmo2, Erbb4, Plcxd3, Nxph2* for immature *Pvalb* interneurons ^128,129^ before the onset of *Pvalb* expression^130^ (Extended Data Fig. 1d). Expression of *Nxph2*, *Cacng2*, *Tac1*, *Ntsr1,* and *Cacna1a* in distinct cell clusters was visualized using UMAP plots.

To examine *Tac1* expression in the caudate and putamen (CPu), we obtained the gene expression matrix from the CPu snRNA-seq datase_t_^123^. This dataset was imported into Python and analyzed using the Scanpy package to identify the cell types and clusters exhibiting *Tac1* enrichment (Extended Data Fig. 7a).

### Electrophysiological feature extraction and analysis

Electrophysiological analyses were performed using a Python-based framework. We extracted 29 electrophysiological properties from the original membrane voltage recordings, employing custom Python scripts that were modified from the Allen Institute’s Software Development Kit to suit our specific research conditions^113^. The analysis quantified a range of parameters, such as the resting membrane potential, input resistance, the membrane’s time constant, rheobase, and action potential features, all deduced from the membrane responses to hyperpolarizing or depolarizing currents. Additionally, we also quantified the maximal action potentials generated during the step protocol, spike-frequency adaptation, sag ratio, rebound amplitude, and the variability and pattern of bursts in interspike intervals.

### Morphological reconstruction and analysis

The cells that passed the morphological criteria were reconstructed and included in the analysis. Neuronal morphologies were reconstructed with Neurolucida (MicroBrightField) and analyzed with Neurolucida Explorer (including Sholl analysis of the dendritic and axonal arbors). For visualization (Figs. 1, 3 and 5; Extended Data Figs. 2-5), reconstructions were imported into Adobe Illustrator (Adobe). For more comprehensive morphological feature analysis, each morphology file was converted into the SWC format with NLMorphologyConverter 0.9.0 (http://neuronland.org) for feature extraction using MorphoPy (https://github.com/berenslab/MorphoPy)^131.^

For clustering analysis of Tac1-positive and Tac1-negative *Pvalb* interneurons, the features from both dendrites and axons were included (the features used for clustering analysis and UMAP embedding calculation are shown in Fig. 1). Morphological parameters were normalized before applying Uniform Manifold Approximation and Projection (UMAP). UMAP embeddings were generated from standardized morphological features using the umaplearn Python library (version 0.5.3). Unsupervised K-means clustering was performed on the normalized features and the confusion matrix was used to show the performance of K-means clustering.

## Supplemental Information

### Extended Data Figures

**Extended Data Fig. 1.**
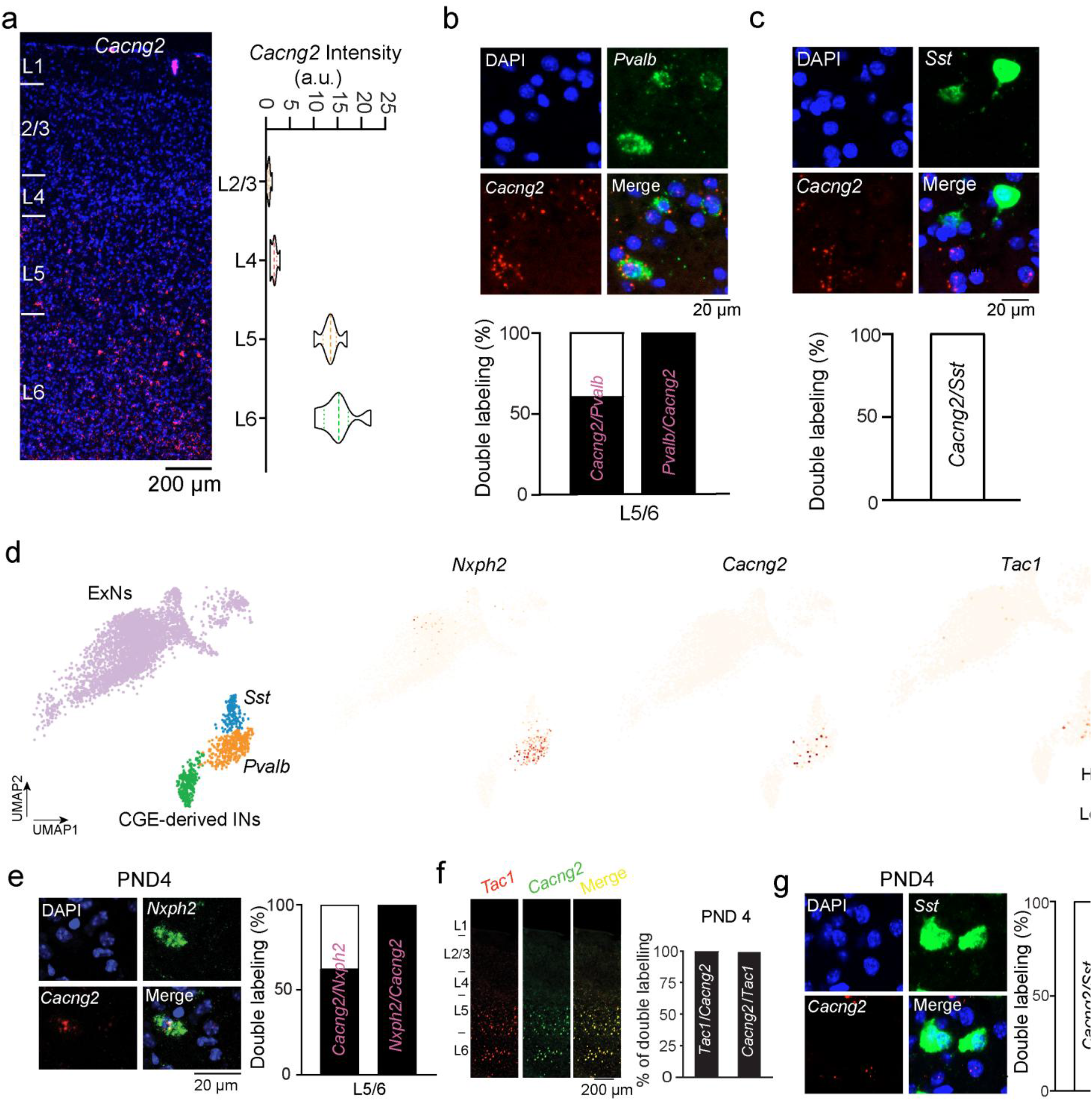
| *Cacng2* expression in the mouse neocortex. Related to Fig. 1. **a,** Left: A representative micrograph of FISH staining for *Cacng2* in a parasagittal section of adult mouse S1. Right: Violin plots of the density of *Cacng2* signals across cortical layers counted across 6 consecutive sections (n= 8 mice). The shape of each violin plot reflects the distribution of data points, whereas the width corresponds to the number of occurrences. **b,** Top: Representative FISH micrographs showing *Pvalb* (green), *Cacng2* (red), and *DAPI* (blue). Bottom: Bar graph depicting the proportion of double-labeled cells (*Pvalb^+^/Cacng2^+^*) cells within the total single-labeled cells (*Pvalb^+^*or *Cacng2^+^*) in layers 5/6 (L5/6) counted across 7 consecutive sections (n = 5 mice). **c,** Top: Representative FISH micrographs showing *Cacng2* (red) and *Sst* (green) and DAPI (blue). Bottom: Bar graph depicts the proportion of double-labeled cells (*Sst^+^/Cacng2^+^*) cells within the total *Sst*^+^ cells counted across 6 consecutive sections (n = 5 mice). **d,** Left: UMAP visualization of the cortical neuronal population collected at postnatal day 4 (P4) (Di Bella et al., 2021), annotated with marker genes, showing immature excitatory neurons (ExN), CGE-derived interneurons, immature *Sst* interneurons, and immature *Pvalb* interneurons. Right: UMAP visualization of normalized expression of *Nxph2* (as a specific marker for immature *Pvlb* interneurons), *Tac1* and *Cacng2*. CGE: caudal ganglionic eminence. **e,** Left: Representative FISH micrographs showing *Nxph2* (green) and *Cacng2* (red) in P4 of mouse S1. Right: Bar graph depicts the proportion of double-labeled cells (*Nxph2^+^/Cacng2^+^*) cells within the total single-labeled cells (*Nxph2^+^or Cacng2^+^*) in layers 5/6 (L5/6) counted across 6 consecutive sections (n=6 mice). **f,** Left: Representative FISH micrograph showing *Cacng2* (green) and *Tac1* (red) in a parasagittal section of mouse S1 at P4. Right: Bar graph depicts the proportion of double-labeled cells (*Tac1^+^/Cacng2^+^*) within the total single-labeled cells (*Tac1^+^* or *Cacng2^+^*) counted across 7 consecutive sections (n = 6 mice). **g,** Left: Representative FISH micrographs showing *Cacng2* (red) and *Sst* (green) and DAPI (blue) at P4. Right: Bar graph depicts the proportion of double-labeled cells (*Sst^+^/Cacng2^+^*) cells within the total *Sst*^+^ cells counted across 6 consecutive sections (n = 6 mice).

**Extended Data Fig. 2.**
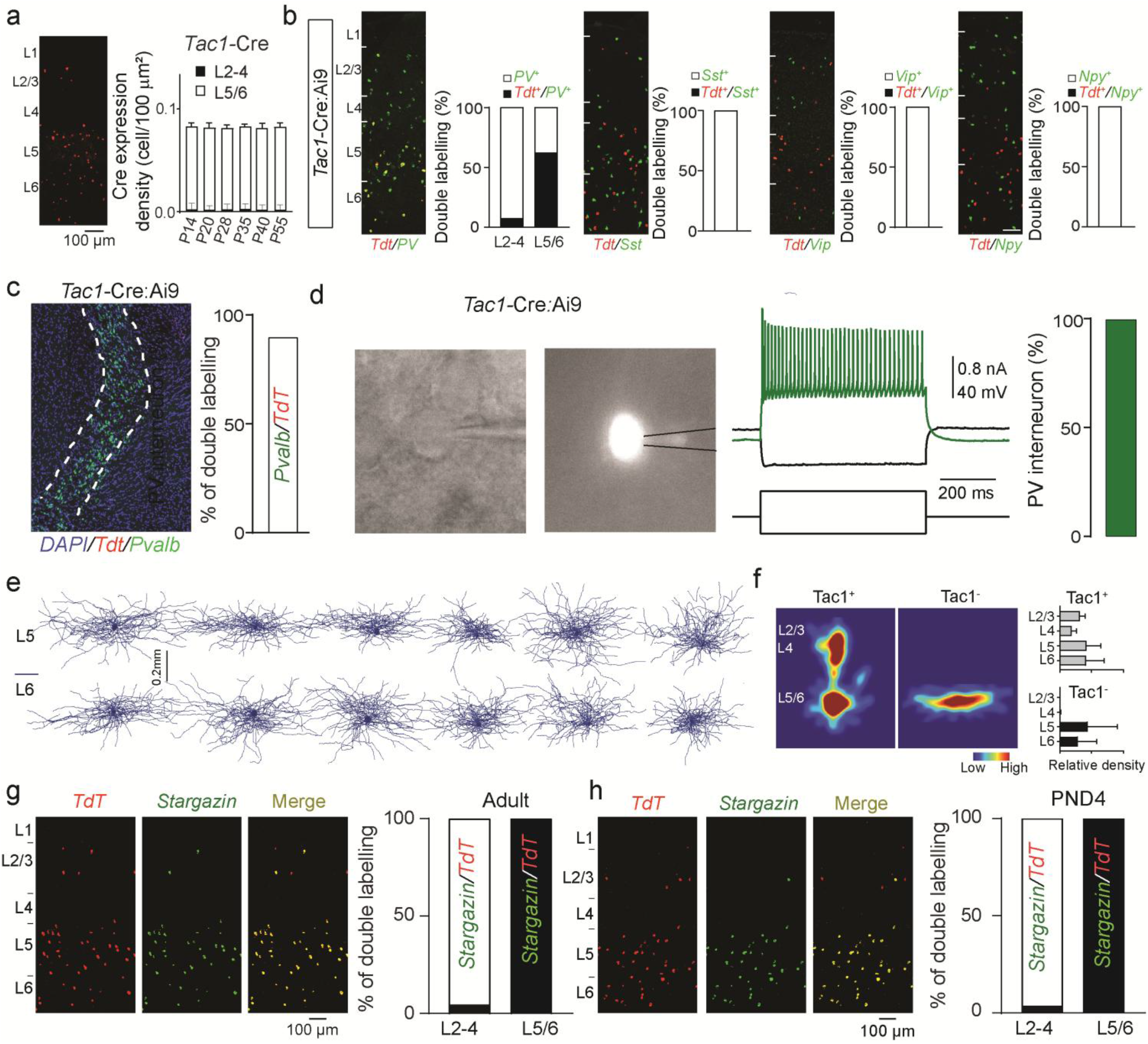
| Characterization of *Tac1*-Cre line in the neocortex. Related to Fig. 1. **a,** Left: Micrograph shows RNA FISH detection of Cre mRNA in the *Tac1*-Cre line at P55. The representative image is the parasagittal S1 section from at least two brain-wide experiments. Right: Developmental characterization of Cre expression in the *Tac1*-Cre line. Quantification in layers 2-4 was performed across 6 consecutive sections from 6 mice at each age. Quantification in layers 5/6 was performed across 6 consecutive sections from 6 mice at P14, P20, and P55, across 5 consecutive sections from 6 mice at P28 and P40, and across 4 consecutive sections from 6 mice at P35. **b,** Representative double FISH staining for *Tdt* mRNA (red) together with *Pvalb*, *Sst*, *Vip*, or *Npy* mRNA (green) in parasagittal S1 sections of adult *Tac1*-Cre:Ai9 mice. Bar graphs depict the proportion of *Tdt*⁺/*Pvalb*⁺ cells within total *Pvalb*⁺ cells in superficial layers (L2-4) and layers 5/6 (L5/6). For *Sst*, *Vip*, and *Npy*, bar graphs depict the proportion of *Tdt*⁺/*Sst*⁺, *Tdt*⁺/*Vip*⁺, and *Tdt*⁺/*Npy*⁺ cells within total *Sst*⁺, *Vip*⁺, and *Npy*⁺ cells, respectively. Quantification for *Pvalb* was performed across 7 consecutive sections from 6 mice; for *Sst*, across 6 consecutive sections from 6 mice; for *Vip*, across 6 consecutive sections from 5 mice; and for *Npy*, across 7 consecutive sections from 6 mice. **c,** Left: Representative micrograph showing double FISH staining for *Tdt* mRNA (red) and *Pvalb* mRNA (green) in the thalamic reticular nucleus (RTN) of adult *Tac1*-Cre:Ai9 mice. Right: Bar graph depicting the proportion of double-labeled cells (*Pvalb*⁺/*Tdt*⁺) within the total single-labeled *Tdt*⁺ cells in the RTN, counted across 7 consecutive sections from 8 mice. **d,** Representative whole-cell recording and firing pattern of a *Tac1*-Cre-labeled interneuron in deep layers of S1. Left: Brightfield image of the recorded neuron. Middle: Fluorescence image of the labeled cell. Right: Representative firing pattern in response to depolarizing current injection. Bar graph shows the proportion of recorded deep-layer *Tac1*-Cre-labeled neurons displaying fast-spiking properties characteristic of *Pvalb* interneurons (n = 20 cells from 8 mice). **e,** Representative morphologies of Tac1^-^ interneurons in layers 5 and 6 of S1. Morphological reconstructions were obtained from n = 12 cells from 8 mice. **f,** Left: Axon density maps of Tac1⁺ and Tac1⁻ interneurons. Middle: Representative layer-wise axon density distributions. Right: Quantification of relative axon density across cortical layers for Tac1⁺ and Tac1⁻ interneurons. Morphological reconstructions were obtained from n = 12 cells per group from 8 mice. **g,** Left: Representative immunofluorescence micrographs showing TdT (red), stargazin (green), and merged signals in the S1 cortex of adult *Tac1*-Cre:Ai9 mice. Right: Bar graphs depicting the proportion of double-labeled cells (stargazin⁺/Tdt⁺) within the total single-labeled Tdt⁺ cells in superficial layers (L2–4) and layers 5/6 (L5/6), counted across 8 sections from 8 mice. In L2–4, 2 of 42 TdT⁺ cells were stargazin⁺/TdT⁺ double-labeled cells (2/42, 4.8%). In L5/6, all TdT⁺ cells were stargazin⁺/TdT⁺ double-labeled cells (329/329, 100%). **h,** Left: Representative immunofluorescence micrographs showing TdT (red), stargazin (green), and merged signals in the S1 cortex of *Tac1*-Cre:Ai9 mice at postnatal day 4 (PND4). Right: Bar graphs depicting the proportion of double-labeled cells (stargazin⁺/Tdt⁺) within the total single-labeled Tdt⁺ cells in superficial layers (L2-4) and layers 5/6 (L5/6), counted across 9 sections from 7 mice. In L2–4, 2 of 47 TdT⁺ cells were stargazin⁺/TdT⁺ double-labeled cells (2/47, 4.3%). In L5/6, all TdT⁺ cells were stargazin⁺/TdT⁺ double-labeled cells (367/367, 100%).

**Extended Data Fig. 3.**
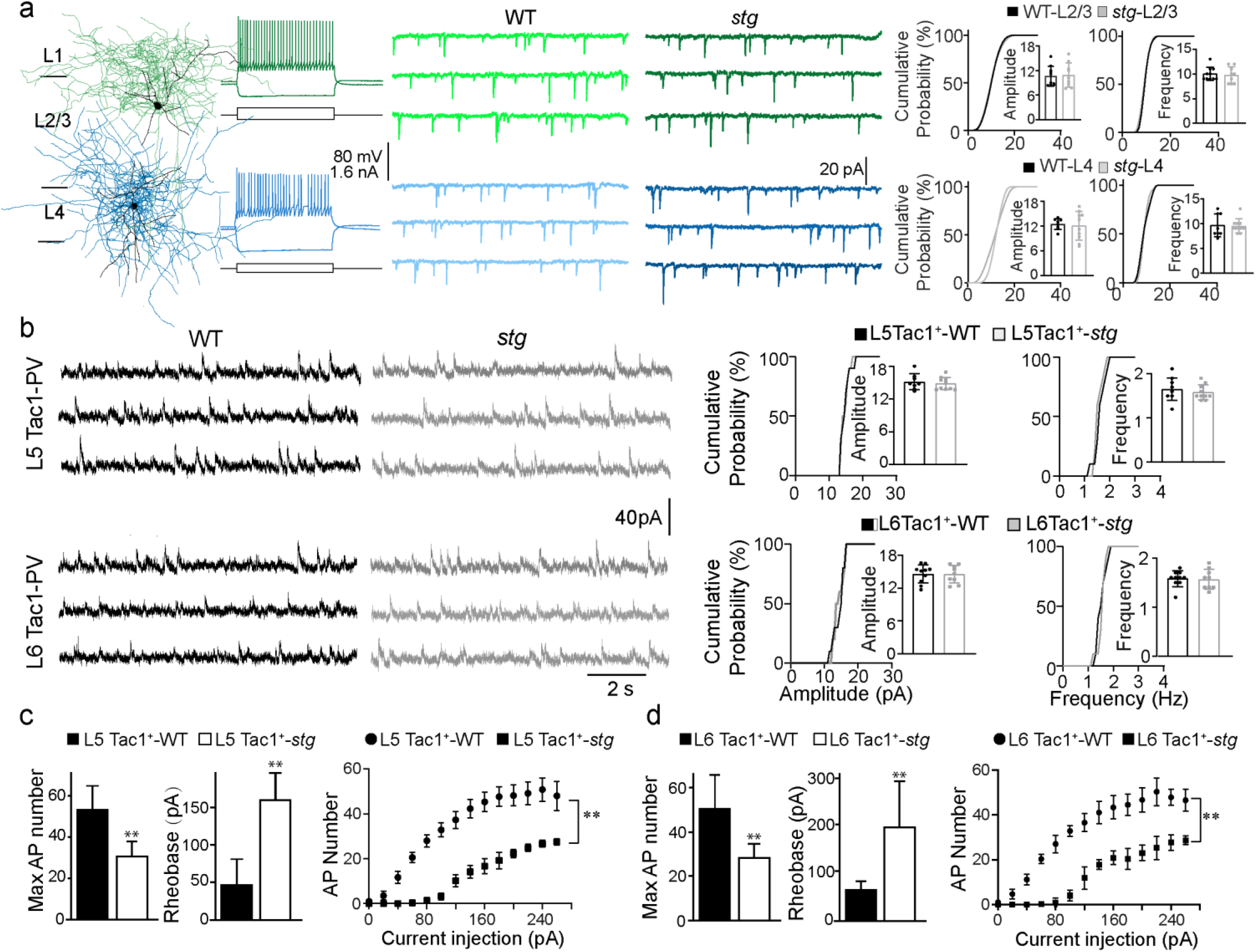
| Characterization of Tac1^+^ *Pvalb* interneurons in *stg*. Related to Fig. 2. **a,** Spontaneous excitatory inputs onto superficial *Pvalb* interneurons (L2/3 and L4). Left: Identification of *Pvalb* interneurons in L2/3 and L4 by their typical morphology and fast-spiking firing patterns. Middle: Representative traces of spontaneous excitatory postsynaptic currents (sEPSCs). Right: Cumulative probability plots and inset bar graphs summarize sEPSC amplitude and frequency in WT and *stg* mice. Cells for WT and *stg*, respectively: L2/3 (n = 10, 9) and L4 (n = 8, 10). Data are color-coded by layer and were analyzed using Student’s t-test. **b,** NMDA receptor-mediated EPSCs in deep-layer (L5/6) Tac1⁺ *Pvalb* interneurons. Left: Representative NMDA-EPSC traces. Right: Cumulative probability plots and inset bar graphs summarize NMDA-EPSC amplitude and frequency in WT and *stg* mice. Cells for WT and *stg*, respectively: L5 (n = 10, 10) and L6 (n = 10, 10). Statistical comparisons were performed using Student’s *t*-test. **c,** Intrinsic excitability of L5 Tac1⁺ *Pvalb* interneurons. Left: Bar graphs showing the maximum number of action potentials (APs) and rheobase in L5 *Tac1*⁺ interneurons from WT and *stg* mice. Right: Firing response to increasing current injections (F-I curve). For both panels, n = 6 neurons per genotype. Maximal AP number and rheobase were compared using Student’s t -test (maximal AP number: t(10) = 4.466, \*\**p* < 0.01; rheobase: t(10) = 5.519, \*\**p* < 0.01), and the F-I curve was analyzed by two-way ANOVA (F(1, 10) = 12.10, \*\**p* < 0.01). See Extended Data Table 1 for additional electrophysiological data. **d,** Intrinsic excitability of L6 Tac1⁺ *Pvalb* interneurons. Left: Bar graphs showing the maximum number of APs and rheobase in L6 Tac1⁺ interneurons from WT and *stg* mice. Right: Firing response to increasing current injections (F-I curve). For both panels, n = 6 neurons per genotype. Maximal AP number and rheobase were compared using Student’s t-test (maximal AP number: t(10) = 3.541, \*\**p* < 0.01; rheobase: t(10) = 3.472, \*\**p* < 0.01), and the F-I curve was analyzed by two-way ANOVA (F(1, 10) = 31.69, \*\**p* < 0.01). See Extended Data Table 1 for additional electrophysiological data.

**Extended Data Fig. 4.**
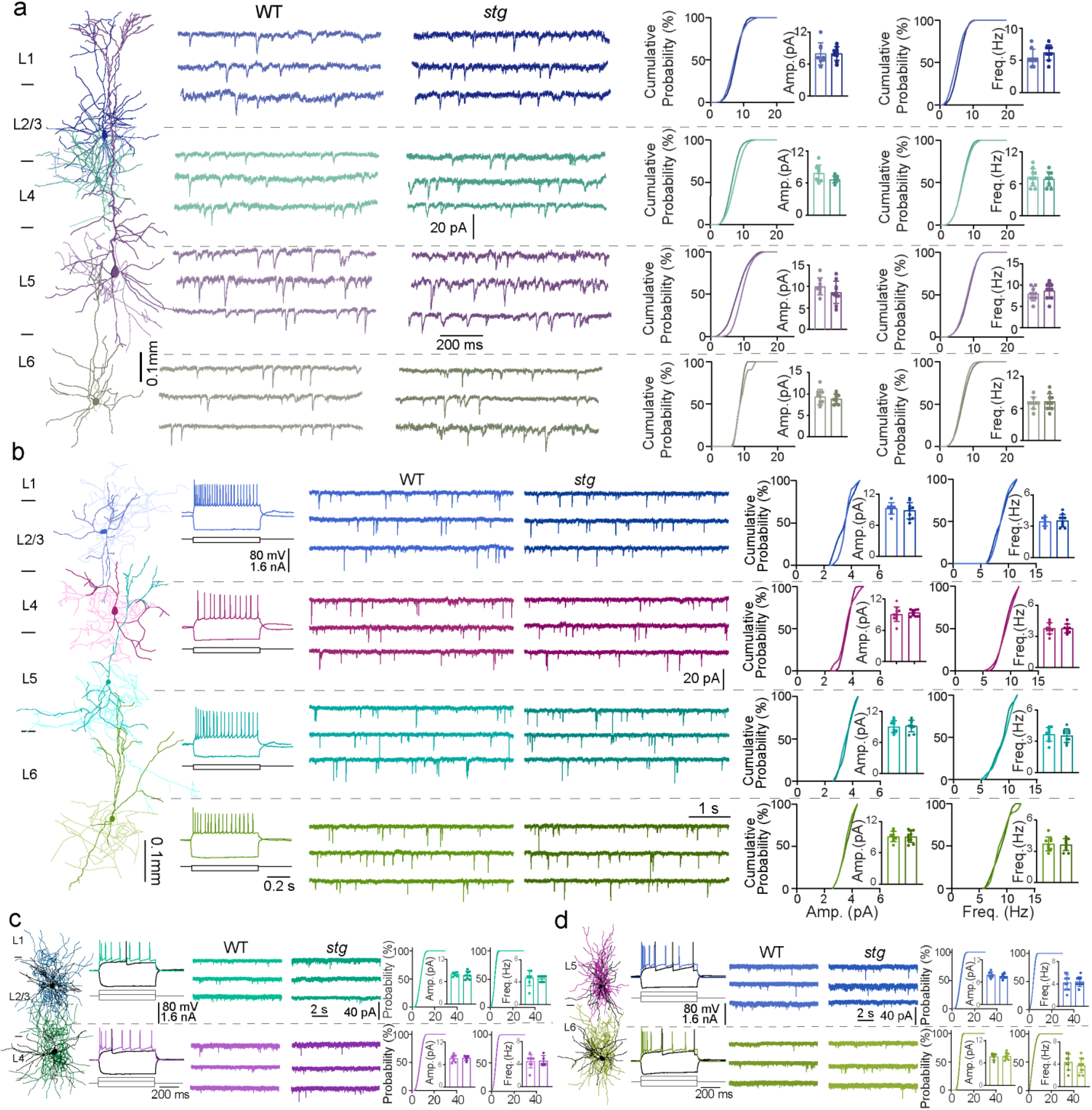
| Synaptic transmission of principal cells (PCs), *Vip* interneurons, and neurogliaform cells across cortical layers in S1 of WT and *stg* mice. Related to Fig. 3. **a**, Spontaneous excitatory postsynaptic currents (sEPSCs) in principal cells (PCs) across layers and genotypes. From left to right: Representative morphology of a recorded PC, sample sEPSC traces from each cortical layer in wild -type (WT) and stargazin mutant (*stg*) mice, and cumulative probability plots with inset bar graphs summarizing sEPSC amplitude and frequency. Cells for WT and *stg*, respectively: L2/3 (n = 8, 9), L4 (n = 10, 9), L5 (n = 8, 10), L6 (n = 9, 10). **b,** sEPSCs in *Vip* interneurons. From left to right: Representative morphology and firing patterns, sample sEPSC traces, and cumulative probability plots with inset bar graphs for sEPSC amplitude and frequency. Cells for WT and *stg*, respectively: L2/3 (n = 8, 10), L4 (n = 10, 9), L5 (n = 10, 9), L6 (n = 9, 10). **c,d,** sEPSCs in neurogliaform cells (NGCs) in superficial (c) and deep (d) layers. In both panels: representative morphology and firing patterns (left), sample sEPSC traces (middle), and summary plots for sEPSC amplitude and frequency (right). Cells for WT and *stg*, respectively: L2/3 (n = 7, 8), L4 (n = 8, 5), L5 (n = 10, 9), L6 (n = 8, 10). Statistical comparisons were performed using Student’s *t*-test.

**Extended Data Fig. 5.**
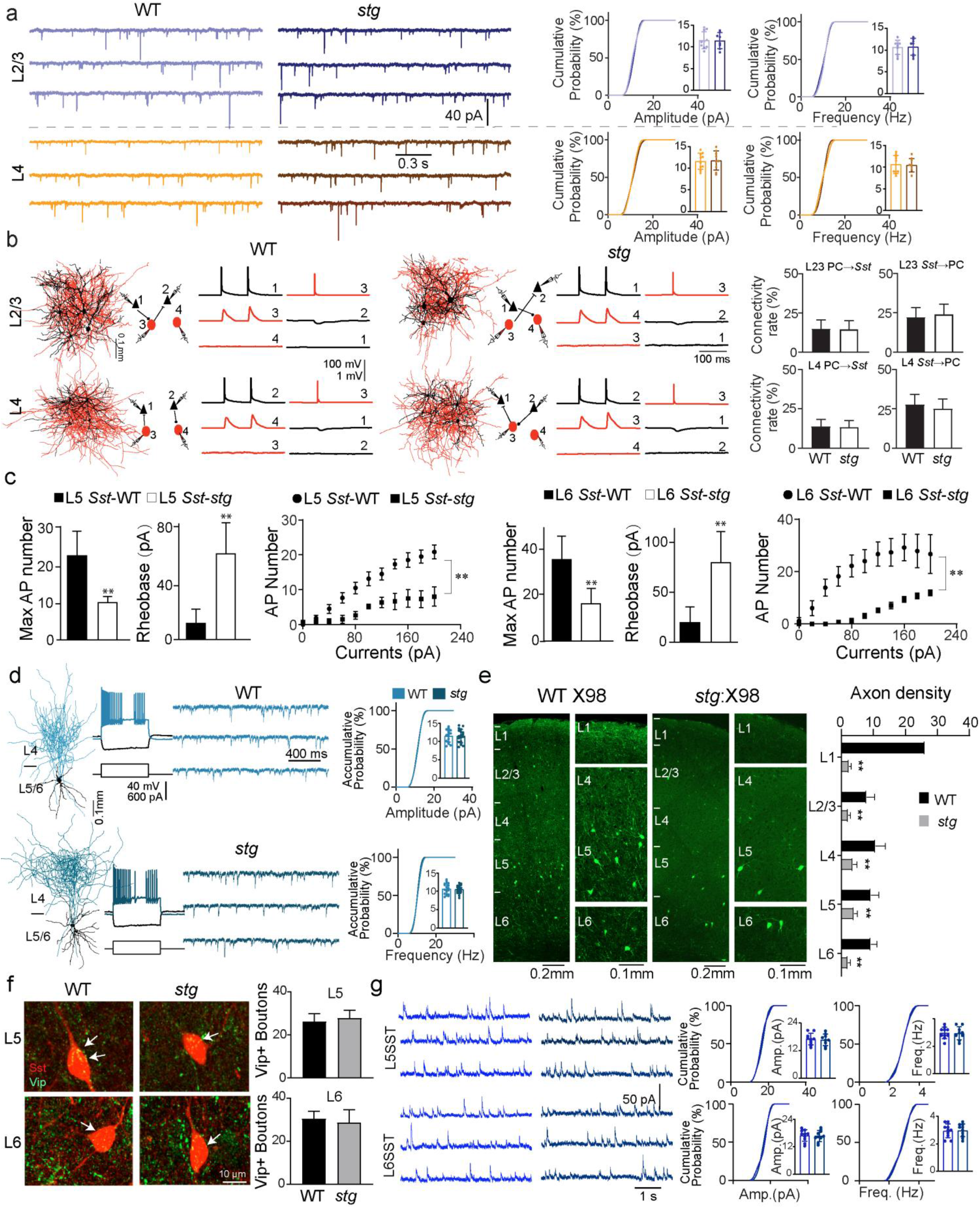
| Characterization of *Sst* interneurons in WT and *stg* mice. Related to Fig. 3. **a,** Sample traces of spontaneous excitatory postsynaptic currents (sEPSCs) recorded in *Sst* interneurons across cortical layers and genotypes (WT vs. *stg*). Data are color-coded by layer. Cumulative probability plots and inset bar graphs summarize sEPSC amplitude and frequency in *Sst* interneurons from each layer. Cells for WT and *stg*, respectively: L2/3 (n = 10, 8) and L4 (n = 10, 9). Statistical comparisons were performed using Student’s *t*-test. **b,** Left: Representative examples of simultaneous 4-cell recordings from principal excitatory cells (PCs, black triangles) and Sst interneurons (SSTs, red circles) in WT and *stg* mice. Reconstructed morphologies of recorded neurons are shown together with schematic connectivity diagrams and representative unitary synaptic traces. Recorded neurons were located close to each other (generally less than 150 μm apart). Vertical scale bars indicate the amplitudes of injected currents in nA, action potentials (APs) in mV, and unitary excitatory or inhibitory postsynaptic potentials (uEPSPs or uIPSPs) in mV. Right: Connection probabilities of PC→*Sst* and *Sst*→PC across cortical layers and genotypes (WT vs. *stg*). PC→*Sst* in L2/3: 6/40 in WT and 6/41 in *stg*; *Sst*→PC in L2/3: 10/45 in WT and 10/40 in *stg*. PC→*Sst* in L4: 8/57 in WT and 8/61 in *stg*; *Sst*→PC in L4: 13/47 in WT and 12/48 in *stg*. **c,** Intrinsic excitability of L5 and L6 *Sst* interneurons. Left panels: Bar graphs showing the maximum number of action potentials (APs) and rheobase in L5 or L6 *Sst* neurons from WT and *stg* mice. Right panels: Firing responses to increasing current injections (F-I curves). For all panels, n = 6 neurons per genotype. For L5 *Sst* interneurons, maximal AP number and rheobase were compared using Student’s *t*-test (maximal AP number: t(10) = 4.815, \*\**p* < 0.01; rheobase: t(10) = 5.010, \*\**p* < 0.01), and the F-I curve was analyzed by two-way ANOVA (F(1, 10) = 25.21, \*\**p* < 0.01). For L6 *Sst* interneurons, maximal AP number and rheobase were compared using Student’s *t*-test (maximal AP number: t(10) = 4.025, \*\**p* < 0.01; rheobase: t(10) = 4.380, \*\**p* < 0.01), and the F-I curve was analyzed by two-way ANOVA (F(1, 10) = 13.75, \*\**p* < 0.01). See Table S1 for additional electrophysiological data. **d,** Quasi-fast-spiking *Sst* interneurons. Left: Representative morphologies. Middle: Representative firing patterns and sample sEPSC traces. Right: Summary plots of sEPSC amplitude and frequency. Cells for WT and *stg*, respectively: n = 16 and n = 19. **e,** Left: Representative micrographs showing axonal distributions in X98 mice. In WT mice, X98-labeled axons span all cortical layers (L1-L6), whereas in *stg*:X98 mice, axonal loss is observed across these layers. Right: Bar graphs quantify axon density across cortical layers (L1, L2/3, L4, L5, and L6) in WT and *stg* mice. For L1: n = 6 sections from 6 WT mice and n = 5 sections from 5 *stg* mice; for L2/3: n = 6 sections from 5 WT mice and n = 6 sections from 5 *stg* mice; for L4: n = 5 sections from 5 WT mice and n = 5 sections from 5 *stg* mice; for L5: n = 5 sections from 4 WT mice and n = 6 sections from 5 *stg* mice; for L6: n = 6 sections from 5 WT mice and n = 6 sections from 6 *stg* mice. Statistical comparisons were performed using Student’s *t*-test (L1: t(9) = 18.76, \*\**p* < 0.01; L2/3: t(10) = 5.614, \*\**p* < 0.01; L4: t(8) = 4.780, \*\**p* < 0.01; L5: t(9) = 4.813, \*\**p* < 0.01; L6: t(10) = 8.521, \*\**p* < 0.01. **f,** Left: Representative micrographs showing immunostaining for Vip in *Sst*-Cre:Ai9 (WT) and *stg*:*Sst*-Cre:Ai9 (*stg*) mice. Arrows denote Vip⁺ boutons overlapping with the soma of *Sst* interneurons. Right: Bar graphs showing the average number of Vip⁺ boutons on each *Sst* interneuron in L5 (upper) and L6 (bottom) from WT and *stg* mice (n = 5 cells for L5-WT, n = 6 cells for L5-*st*g, n = 5 cells for L6-WT, and n = 6 cells for L6-*stg*). **g,** Left: Sample traces of spontaneous inhibitory postsynaptic currents (sIPSCs) recorded in L5/6 *Sst* interneurons from WT and *stg* mice. Right: Cumulative probability plots and inset bar graphs summarize sIPSC amplitude and frequency in L5 (upper) and L6 (lower) *Sst* interneurons. Cells for WT and *stg*, respectively: L5 (n = 10, 9) and L6 (n = 10, 10). Statistical comparisons were performed using Student’s *t*-test.

**Extended Data Fig. 6.**
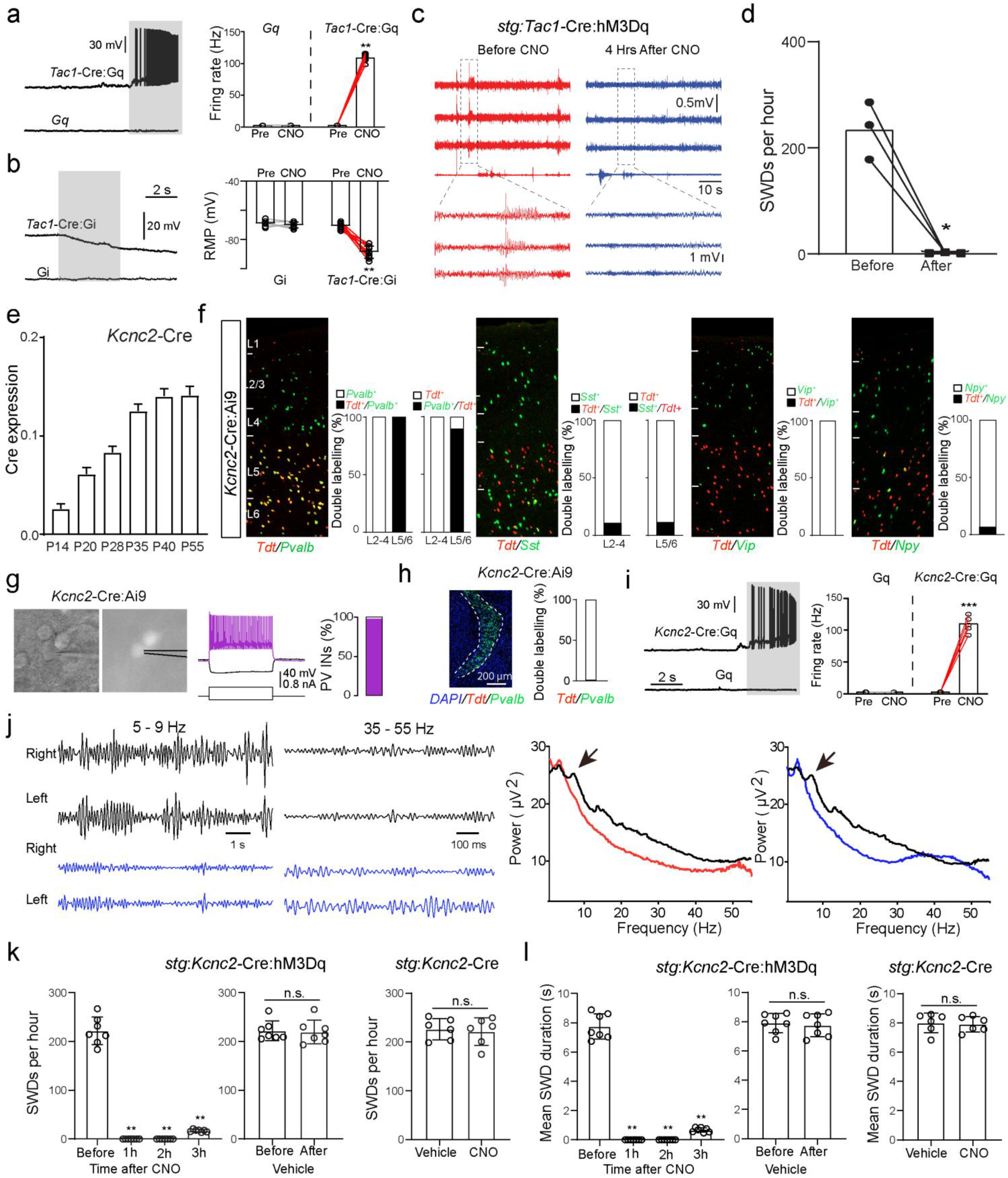
| Validation of chemogenetic manipulations and characterization of the Kcnc2-Cre line. Related to Fig. 4. **a,** Representative membrane potential traces from recorded L5/6 neurons in *Tac1*-Cre:Gq mice in response to clozapine-N-oxide (CNO, 10 µM; gray shading). Right, summary of firing rates before and after CNO administration. n = 8 cells from 3 *Tac1*-Cre:Gq mice; paired t-test, t(7) = 4.562, \*\**p* < 0.01. **b,** Representative membrane potential traces from recorded L5/6 neurons in *Tac1*-Cre:Gi mice in response to clozapine-N-oxide (CNO, 10 µM; gray shading). Right, summary of membrane potential before and after CNO administration. n = 9 cells from 6 *Tac1*-Cre:Gi mice; paired t-test, t(8) = 3.401, \*\**p* < 0.01. **c,** To test whether orally administered CNO acts rapidly in the brain, CNO was provided to *stg*:Tac1-Cre:hM3Dq mice and seizure activity was monitored. Representative EEG traces recorded before (red) and 4 h after CNO drinking (40 mg/L; blue) show a marked reduction in spike-wave discharges (SWDs). Bottom, expanded traces. **d,** Quantification of SWDs per hour within a 1-hour EEG window (approximately 12:00-13:00) demonstrates a significant reduction in discharge frequency following CNO administration. n = 3 mice; paired t-test, t(2) = 7.311, \**p* < 0.05. **e,** Bar graph showing Cre-dependent tdTomato expression driven by *Kcnc2*-Cre across cortical layers at different developmental stages (P14–P55). *Kcnc2*-Cre: n = 5 slices from 3 mice at P14; n = 4 slices from 4 mice at P20; n = 5 slices from 4 mice at P28; n = 6 slices from 3 mice at P35; n = 5 slices from 5 mice at P40; n = 6 slices from 6 mice at P55. **f,** FISH staining for the cardinal interneuron markers *Pvalb*, *Sst*, *Vip*, and *Npy* in *Kcnc2*-Cre:Ai9 mice at P45 across cortical layers. TdTomato (*Tdt*) signals predominantly overlap with *Pvalb* in L5/6, with minor overlap with *Sst*, *Vip*, and *Npy* in L2/3. n = 5 slices from 5 mice for *Pvalb*; n = 5 slices from 4 mice for *Sst*; n = 3 slices from 4 mice for *Vip*; n = 3 slices from 4 mice for *Npy*. **g,** Electrophysiological characterization of labeled neurons in *Kcnc2*-Cre:Ai9 mice by patch-clamp recording. Left, example patch-clamp image; middle, representative firing pattern; right, quantification. 45 recorded neurons in *Kcnc2*-Cre:Ai9 mice were identified as *Pvalb* interneurons based on their fast-spiking properties. **h,** Representative membrane potential traces from recorded L5/6 neurons in *Kcnc2*-Cre:Gq mice in response to clozapine-N-oxide (CNO, 10 µM; gray shading). Right, summary of firing rates before and after CNO administration. n = 6 cells from 4 *Kcnc2*-Cre:Gq mice; paired t-test, t(5) = 22.85, \*\*\**p* < 0.001. **i,** Left: FISH staining for *Pvalb* in the thalamic reticular nucleus (TRN) of *Kcnc2*-Cre:Ai9 mice. Right: Quantification of the percentage of TdTomato-positive cells overlapping with *Pvalb* in the TRN. n = 4 slices from 3 mice. **j,** Left: EEG frequency band traces before (top, black) and after chemogenetic activation (bottom, blue) in *stg*:*Kcnc2*-Cre:hM3Dq mice, showing disappearance of the dominant 5-9 Hz absence seizure activity after CNO treatment. Right: Power spectrum analysis shows a marked reduction in 5-9 Hz power (black, baseline; red, shortly after activation; blue, 2 h after activation), consistent with seizure suppression (arrows), together with an increase in gamma-band power (35-55 Hz). **k,** Quantification of SWDs per hour in *stg*:*Kcnc2*-Cre:hM3Dq mice before and after CNO treatment. Data were analyzed using one-way repeated-measures ANOVA, followed by Dunnett’s multiple comparisons test against baseline (Before). A significant effect of time was observed (F(1.015, 6.089) = 403.2, \*\**p* < 0.01). Dunnett’s test showed significant differences between Before and 1 h, 2 h, and 3 h (all \*\**p* < 0.01). n = 7 mice. Vehicle treatment produced no significant change (n = 7 mice, paired t-test). CNO treatment in control *stg*:*Kcnc2*-Cre mice without hM3Dq expression also produced no significant change (n = 6 mice, Student’s t-test). **l,** Quantification of mean SWD duration in *stg*:*Kcnc2*-Cre:hM3Dq mice before and after CNO treatment. Data were analyzed using one-way repeated-measures ANOVA, followed by Dunnett’s multiple comparisons test against baseline (Before). A significant effect of time was observed (F(1.025, 6.150) = 471.7, \*\**p* < 0.01). Dunnett’s test showed significant differences between Before and 1 h, 2 h, and 3 h (all \*\**p* < 0.01). n = 7 mice. Vehicle treatment produced no significant change (n = 7 mice, paired t-test). CNO treatment in control *stg*:*Kcnc2*-Cre mice without hM3Dq expression also produced no significant change (n = 6 mice, Student’s t-test).

**Extended Data Fig. 7.**
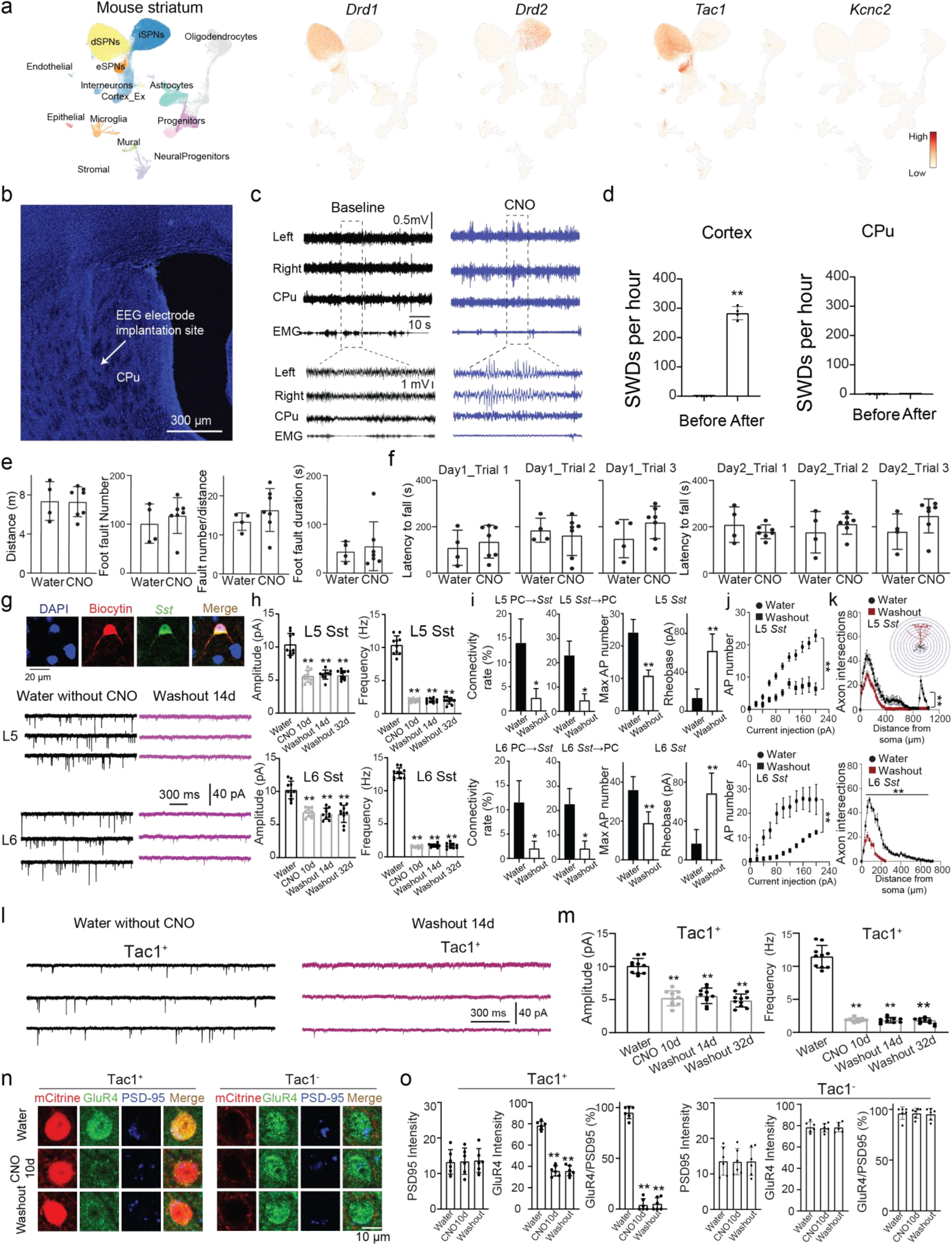
| Chronic Tac1^+^ interneuron inhibition induces cortical but not striatal spike-wave activity and persistent circuit remodeling. Related to Fig. 5. **a,** UMAP visualization of mouse CPu neurons adapted from Anderson et al. (2023), showing *Tac1* co-expression in the *Drd1* but not *Drd2* cluster. **b,** Representative image of the EEG electrode implantation site in the CPu core. **c,d,** EEG recordings show that 10-14 days of CNO treatment in *Tac1*-Cre:Gi mice induces seizure activity in the cortex but not in the CPu. Quantification in (d), paired t-test, t(3) = 25.18, \*\**p* < 0.01. **e,f,** Foot-slip and rotarod tests show no significant motor deficits after CNO treatment. Water, n = 4; CNO, n = 7; unpaired Student’s t -test. **g,h,** Representative traces and quantification of sEPSCs in L5 and L6 Sst interneurons. CNO treatment significantly reduced sEPSC amplitude and frequency, with persistent effects after washout. n = 10 cells per group. One-way ANOVA with Dunnett’s multiple comparisons test versus Water: L5 amplitude, F(3, 36) = 49.49, \*\**p* < 0.01; L5 frequency, F(3, 36) = 297.9, \*\**p* < 0.01; L6 amplitude, F(3, 36) = 27.23, \*\**p* < 0.01; L6 frequency, F(3,36) = 1412, \*\**p* < 0.01; all Dunnett’s comparisons vs Water, \*\**p* < 0.01. **i,** Persistent impairment of synaptic connectivity and intrinsic excitability of deep-layer *Sst* interneurons 5 months after CNO withdrawal. Left, summary of connection probabilities for PC→*Sst* and *Sst*→PC pairs in L5 and L6. PC→*Sst* connectivity: L5, 7/50 in Water and 2/74 in Washout; L6, 6/52 in Water and 2/96 in Washout. *Sst*→PC connectivity: L5, 12/52 in Water and 3/65 in Washout; L6, 10/44 in Water and 2/45 in Washout. \**p* < 0.05. Right, maximal action potential (AP) number remained significantly reduced and rheobase remained significantly increased in both L5 and L6 *Sst* interneurons after the 5-month washout period. n = 6 cells per group. Student’s t-test; L5 maximal AP number, t(10) = 7.467, \*\**p* < 0.01; L5 rheobase, t(10) = 6.049, \*\**p* < 0.01; L6 maximal AP number, t(10) = 4.656, \*\**p* < 0.01; L6 rheobase, t(10) = 5.018, \*\**p* < 0.01. **j,** Firing responses to increasing current injections in L5 and L6 *Sst* interneurons 5 months after CNO withdrawal. Firing responses remained significantly reduced in Washout compared with Water mice. n = 6 cells per group; two-way ANOVA: L5, F(1, 10) = 48.74, \*\**p* < 0.01; L6, F(1, 10) = 12.58, \*\**p* < 0.01. **k,** Sholl analysis of axonal arborization in L5 and L6 *Sst* interneurons 5 months after CNO withdrawal. Axonal intersections remained significantly reduced in Washout compared with Water mice. L5, n = 12 cells per group, two-way ANOVA, F(1, 22) = 31.91, \*\**p* < 0.01; L6, n = 6 cells per group, two-way ANOVA, F(1, 10) = 153.9, \*\**p* < 0.01. **l,m,** Representative traces and quantification of sEPSCs in *Tac1*^+^ interneurons. CNO treatment significantly reduced sEPSC amplitude and frequency, with persistent effects after washout. n = 10 cells per group. One-way ANOVA with Dunnett’s multiple comparisons test versus Water: amplitude, F(3, 36) = 48.35, \*\**p* < 0.01; frequency, F(3, 36) = 282.7, \*\**p* < 0.01; all Dunnett’s comparisons vs Water, \*\**p* < 0.01. **n,o,** Representative images and quantification of synaptic GluR4 in *Tac1*-Cre:Gi mice. CNO treatment disrupted GluR4 synaptic trafficking in Tac1^+^ neurons but not *Tac1*^−^ neurons, without changing total PSD95 levels. Red, mCitrine; green, GluR4; blue, PSD95. Washout, 14 days. In Tac1^+^ neurons, GluR4 intensity was reduced (one-way ANOVA, F(2, 15) = 173.6, \*\**p* < 0.01) and GluR4/PSD95 colocalization was decreased (F(2, 15) = 448.4, \*\**p* < 0.01), followed by Dunnett’s multiple comparisons test versus Water (all \*\**p* < 0.01). n = 6 cells per group. No significant changes were detected in Tac1^−^ neurons.

**Extended Data Fig. 8.**
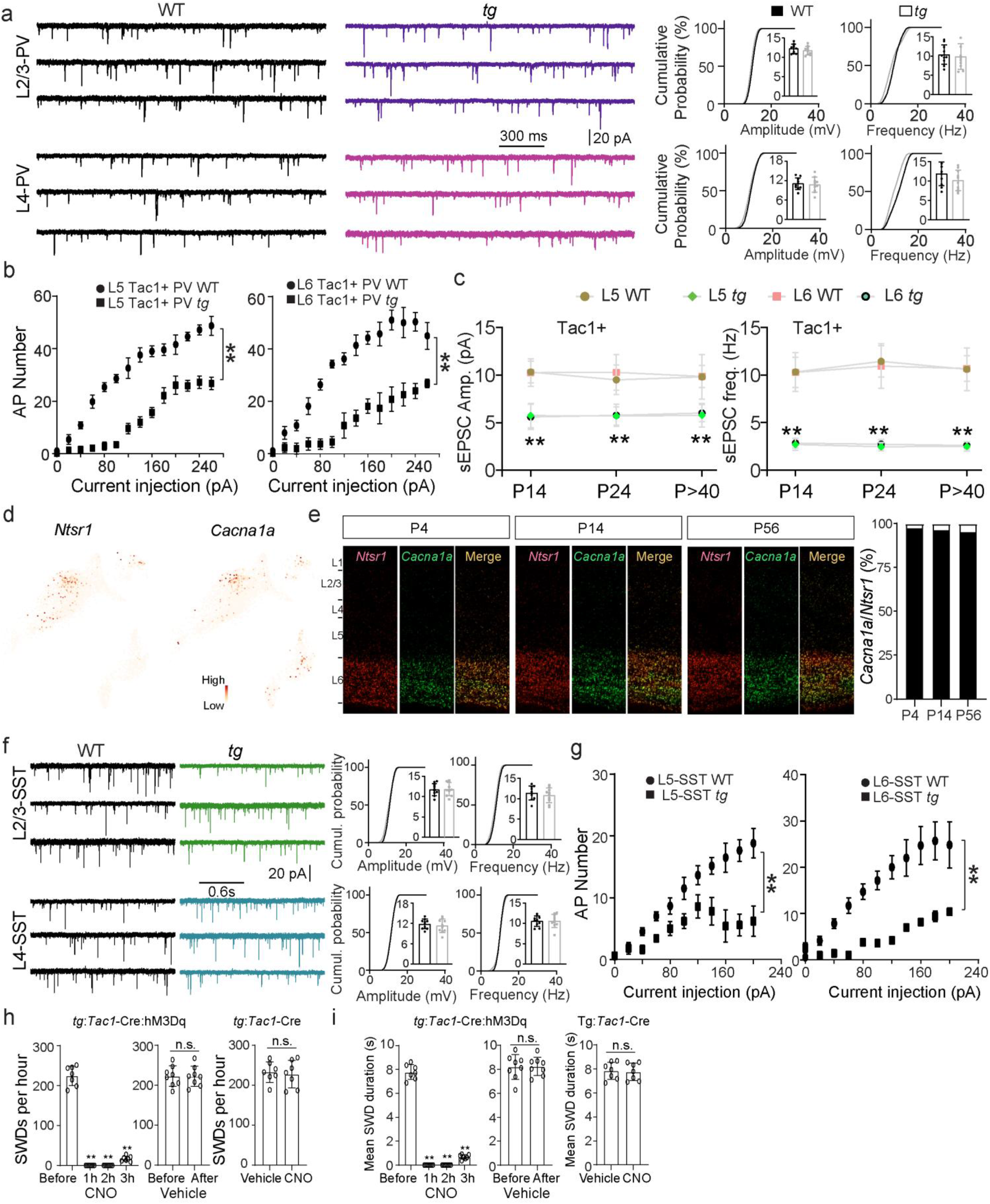
| Characterization of cell type-specific synaptic and intrinsic excitability defects, and developmental *Cacna1a* expression in *tg*. Related to Fig. 6. **a,** Representative traces of spontaneous excitatory postsynaptic currents (sEPSCs) recorded from L2/3 *Pvalb* (PV) interneurons and L4 PV interneurons in WT and *tg* mice. Cumulative probability plots and inset bar graphs show no significant differences in sEPSC amplitude or frequency between WT and Tg mice in either L2/3 or L4 PV interneurons. n = 10 cells for L2/3 PV WT, n = 10 cells for L2/3 PV *tg*; n = 10 cells for L4 PV WT, n = 10 cells for L4 PV *tg*. **b,** Firing responses, measured as the number of action potentials evoked by increasing current step injections, in L5 Tac1^+^ interneurons (left) and L6 Tac1^+^ interneurons (right) from WT and *tg* mice. In L5 Tac1^+^ interneurons, WT: n = 5 cells, *tg*: n = 7 cells; two-way ANOVA, F(1, 10) = 215.3, \*\**p* < 0.01. In L6 Tac1^+^ interneurons, WT: n = 5 cells, *tg*: n = 5 cells; two-way ANOVA, F(1, 8) = 74.62, \*\**p* < 0.01. **c,** Quantification of sEPSC amplitude and frequency in L5 and L6 Tac1^+^ interneurons across developmental stages (P14, P24, and P > 40) in WT and *tg* mice. For L5 *Tac1*^+^ interneurons: P14, n = 9 WT and n = 9 *tg*; P24, n = 10 WT and n = 8 *tg*; P > 40, n = 8 WT and n = 10 *tg*. For L6 *Tac1*^+^ interneurons: P14, n = 7 WT and n = 7 *tg*; P24, n = 10 WT and n = 10 *tg*; P > 40, n = 8 WT and n = 9 *tg*. Student’s t-test was used for WT versus *tg* comparisons at each developmental stage; significant differences are indicated by \*\**p* < 0.01. **d,** UMAP visualization of cortical neuronal populations at P4 showing strong co-distribution of *Ntsr1* and *Cacna1a*. **e,** Double staining for *Ntsr1* (L6 corticothalamic marker, red) and *Cacna1a* (green) at P4, P14, and P56. Merged images are shown on the right. The proportion of *Ntsr1* neurons co-expressing *Cacna1a* was 97.8% at P4 (n = 6 mice), 96.7% at P14 (n = 7 mice), and 95.6% at P56 (n = 10 mice). **f,** Representative traces of sEPSCs recorded from L2/3 *Sst* interneurons and L4 *Sst* interneurons in WT and *tg* mice. Cumulative probability plots and inset bar graphs show no significant differences in sEPSC amplitude or frequency between WT and Tg mice in either L2/3 or L4 *Sst* interneurons. N = 10 cells for L2/3 *Sst* WT, n = 10 cells for L2/3 *Sst tg*; n = 10 cells for L4 *Sst* WT, n = 10 cells for L4 *Sst tg*. **g,** Firing responses, measured as the number of action potentials evoked by increasing current step injections, in L5 *Sst* interneurons (left) and L6 *Sst* interneurons (right) from WT and Tg mice. In L5 *Sst* interneurons, WT: n = 6 cells, *tg*: n = 5 cells; two-way ANOVA, F(1,9) = 18.91, \*\**p* < 0.01. In L6 *Sst* interneurons, WT: n = 7 cells, *tg*: n = 5 cells; two-way ANOVA, F(1,10) = 15.64, \*\**p* < 0.01. **h,** Quantification of spike-wave discharges (SWDs) per hour in *tg*:*Tac1*-Cre:hM3Dq mice before and after CNO treatment. Data were analyzed using one-way repeated-measures ANOVA followed by Dunnett’s multiple comparisons test against baseline (Before). CNO treatment produced a significant time-dependent effect on SWD frequency (F(1.123, 6.736) = 537.4, *p* < 0.01). Post hoc Dunnett’s comparisons revealed significant reductions at 1 h, 2 h, and 3 h compared with Before (all \*\**p* < 0.01). n = 7 mice. Vehicle-treated mice showed no significant change in SWD frequency (n = 8 mice, paired t-test). In control *tg*:*Tac1*-Cre mice lacking hM3Dq expression, CNO treatment also did not significantly alter SWD frequency (n = 7 mice, Student’s t-test). **i,** Quantification of mean SWD duration in *tg*:*Tac1*-Cre:hM3Dq mice before and after CNO treatment. Data were analyzed using one-way repeated-measures ANOVA followed by Dunnett’s multiple comparisons test against baseline (Before). A significant effect of time was detected after CNO treatment (F(1.086, 6.517) = 854.5, *p* < 0.01). Dunnett’s post hoc analysis showed that mean SWD duration was significantly different at 1 h, 2 h, and 3 h compared with Before (all \*\**p* < 0.01). n = 7 mice. No significant change was observed after vehicle treatment (n = 8 mice, paired t -test). Similarly, CNO treatment did not significantly affect mean SWD duration in control *tg*:Tac1-Cre mice without hM3Dq expression (n = 7 mice, Student’s t-test).

