## Supplemental table1 for "Convergent Inhibitory Cortical Circuit Disruption Drives Genetically Distinct Absence Seizures"

**Supplementary Data Table 1. Electrophysiological properties of PV and SST interneurons in tg mice, related to Fig.6**

| **CellType** | **PV** | | | | | | | | **SST** | | | | | | | |
|---|---|---|---|---|---|---|---|---|---|---|---|---|---|---|---|---|
| **Layer** | **II/III** | | **IV** | | **V** | | **VI** | | **II/III** | | **IV** | | **V** | | **VI** | |
| **Group** | **WT** | ***tg*** | **WT** | ***tg*** | **WT** | ***tg*** | **WT** | ***tg*** | **WT** | ***tg*** | **WT** | ***tg*** | **WT** | ***tg*** | **WT** | ***tg*** |
| **RMP (mV)** | -72.03±3.69 | -72.17±1.02 | -73.52±2.97 | -70.33±10.75 | -72.77±2.13 | -75.66±5.03 | -68.70±3.65 | -72.88±6.68 | -73.26±1.70 | -66.55±8.35 | -67.06±7.81 | -72.20±8.33 | -70.25±1.40 | -76.12±4.09 | -72.23±10.58 | -73.21±4.96 |
| **Input resistance (MOhm)** | 279.58±83.05 | 230.23±71.06 | 301.32±71.43 | 339.73±37.43 | 290.52±52.38 | 239.82±84.60 | 232.32±105.60 | 246.50±121.40 | 318.08±93.30 | 219.30±87.42 | 325.37±59.52 | 313.99±141.32 | 282.27±93.51 | 273.37±68.29 | 339.19±156.92 | 249.08±58.40 |
| **Membrane time constant (ms)** | 12.20±3.06 | 10.80±1.78 | 16.70±7.30 | 15.99±3.46 | 13.03±5.36 | 10.81±0.82 | 9.83±2.88 | 11.43±2.85 | 29.32±10.28 | 25.83±10.88 | 35.39±7.29 | 30.18±7.20 | 28.33±7.94 | 12.92±11.83 | 27.79±14.59 | 21.77±3.29 |
| **AP threshold (mV)** | -42.42±5.64 | -42.50±6.45 | -38.79±4.82 | -38.90±4.54 | -46.66±9.24 | -38.50±2.20 | -40.95±4.39 | -40.97±6.90 | -43.81±4.17 | -41.06±1.79 | -46.39±5.86 | -43.66±5.51 | -38.04±6.69 | -41.01±4.79 | -43.27±2.79 | -38.99±11.33 |
| **AP amplitude (mV)** | 85.87±9.16 | 87.59±5.50 | 85.75±6.76 | 82.25±8.53 | 87.39±1.56 | 83.79±2.05 | 81.52±2.87 | 81.30±4.27 | 95.08±5.53 | 87.66±5.39 | 95.06±4.74 | 92.31±5.40 | 93.57±4.94 | 88.32±11.55 | 85.79±6.64 | 90.23±8.32 |
| **AP width (ms)** | 0.68±0.20 | 0.59±0.10 | 0.66±0.17 | 0.72±0.12 | 0.67±0.13 | 0.68±0.07 | 0.55±0.12 | 0.73±0.12 | 1.29±0.54 | 1.24±0.51 | 1.17±0.19 | 1.05±0.23 | 0.98±0.10 | 1.16±0.54 | 0.81±0.18 | 1.04±0.12 |
| **Upstroke-to-downstroke ratio** | 1.92±0.34 | 1.66±0.24 | 1.78±0.26 | 1.96±0.34 | 1.70±0.22 | 2.11±0.22 | 1.66±0.31 | 1.93±0.23 | 3.70±1.76 | 3.45±1.27 | 3.08±0.81 | 3.24±1.51 | 2.90±0.35 | 4.06±2.32 | 2.20±0.64 | 3.37±0.57 |
| **Afterhyperpolarization (mV)** | -17.92±4.30 | -22.26±2.31 | -24.57±3.37 | -20.23±3.06 | -15.86±3.81 | -21.65±2.49 | -18.86±1.95 | -18.79±7.98 | -11.04±8.81 | -16.19±5.16 | -12.72±5.19 | -11.75±4.46 | -11.83±1.69 | -14.39±7.53 | -17.06±5.27 | -12.46±4.87 |
| **Afterdepolarization (mV)** | 0.00±0.00 | 0.00±0.00 | 0.00±0.00 | 0.00±0.00 | 0.00±0.00 | 0.00±0.00 | 0.00±0.00 | 0.00±0.00 | 0.00±0.00 | 1.04±2.34 | 0.00±0.00 | 0.00±0.00 | 2.54±2.40 | 0.25±0.55 | 0.23±0.62 | 0.00±0.00 |
| **ISI adaptation index** | 1.20±0.17 | 1.16±0.06 | 1.10±0.10 | 1.01±0.38 | 1.24±0.23 | 1.10±0.04 | 1.09±0.07 | 1.14±0.15 | 1.31±0.52 | 1.33±0.15 | 1.17±0.05 | 1.18±0.20 | 1.17±0.12 | 1.21±0.06 | 1.15±0.09 | 1.32±0.07 |
| **Sag ratio** | 1.09±0.08 | 1.08±0.05 | 1.17±0.09 | 1.09±0.06 | 1.10±0.04 | 1.05±0.02 | 1.12±0.11 | 1.17±0.20 | 1.23±0.12 | 1.11±0.08 | 1.11±0.07 | 1.13±0.08 | 1.11±0.02 | 1.05±0.28 | 1.17±0.14 | 1.10±0.10 |
| **Latency (ms)** | 97.82±95.17 | 101.05±125.15 | 63.63±37.09 | 72.06±78.85 | 61.59±33.36 | 31.71±9.31 | 34.03±20.68 | 59.23±57.15 | 106.83±38.82 | 131.92±118.34 | 86.47±10.68 | 137.19±199.28 | 94.17±32.84 | 118.83±136.96 | 122.56±65.06 | 98.07±20.91 |
| **Latency @ +20pA current (ms)** | 33.47±25.49 | 63.94±114.88 | 24.94±5.50 | 47.51±63.46 | 24.07±12.04 | 20.96±4.54 | 18.74±8.02 | 28.52±20.56 | 60.36±18.35 | 46.56±33.92 | 39.92±7.40 | 89.49±106.56 | 54.67±17.58 | 64.43±65.33 | 30.26±12.96 | 58.14±10.57 |
| **Spike frequency adaptation** | 0.86±0.06 | 0.82±0.14 | 0.89±0.09 | 0.87±0.08 | 0.91±0.06 | 0.72±0.09 | 0.78±0.10 | 0.66±0.22 | 0.60±0.18 | 0.47±0.24 | 0.63±0.11 | 0.50±0.29 | 0.61±0.16 | 0.49±0.28 | 0.68±0.26 | 0.65±0.21 |
| **ISI Fano factor** | 0.00±0.00 | 0.00±0.00 | 0.00±0.00 | 0.00±0.00 | 0.00±0.00 | 0.00±0.00 | 0.00±0.00 | 0.01±0.03 | 0.01±0.01 | 0.02±0.03 | 0.00±0.00 | 0.01±0.00 | 0.00±0.00 | 0.01±0.00 | 0.01±0.03 | 0.00±0.00 |
| **ISI coefficient of variation** | 0.14±0.05 | 0.24±0.30 | 0.10±0.06 | 0.11±0.03 | 0.09±0.03 | 0.22±0.08 | 0.19±0.10 | 0.44±0.57 | 0.35±0.14 | 0.39±0.28 | 0.29±0.06 | 0.34±0.16 | 0.35±0.18 | 0.43±0.18 | 0.35±0.44 | 0.34±0.13 |
| **ISI average adaptation index** | 1.12±0.13 | 1.03±0.01 | 1.31±0.66 | 1.04±0.07 | 1.03±0.02 | 1.03±0.01 | 1.07±0.07 | 1.09±0.05 | 1.29±0.33 | 1.28±0.26 | 1.16±0.07 | 1.17±0.14 | 1.21±0.16 | 1.22±0.07 | 1.09±0.13 | 1.25±0.16 |
| **Rebound (mV)** | 1.06±2.15 | 1.30±0.67 | 4.06±1.84 | 3.05±2.30 | 3.82±2.89 | 1.25±0.93 | 0.37±0.52 | 1.06±1.86 | 4.35±1.58 | 3.94±3.51 | 3.65±1.76 | 2.96±1.45 | 3.40±0.89 | 0.87±0.85 | 4.52±4.04 | 2.48±3.34 |
| **Sag time (s)** | 0.19±0.20 | 0.29±0.16 | 0.38±0.06 | 0.38±0.08 | 0.19±0.09 | 0.21±0.10 | 0.07±0.14 | 0.19±0.21 | 0.40±0.07 | 0.41±0.14 | 0.32±0.20 | 0.30±0.14 | 0.44±0.04 | 0.21±0.15 | 0.39±0.15 | 0.33±0.16 |
| **Sag area (mV*s)** | 22.56±24.35 | 31.63±16.41 | 40.43±5.85 | 43.90±13.50 | 22.21±10.22 | 22.08±10.46 | 6.89±12.23 | 19.14±21.11 | 48.92±11.71 | 47.92±17.34 | 38.06±21.45 | 37.42±21.96 | 56.77±7.63 | 20.17±14.47 | 54.46±24.44 | 38.12±19.42 |
| **AP amplitude adaptation index** | 0.94±0.01 | 0.91±0.03 | 0.94±0.03 | 0.93±0.03 | 0.93±0.04 | 0.92±0.01 | 0.94±0.02 | 0.90±0.03 | 0.87±0.11 | 0.85±0.07 | 0.91±0.02 | 0.91±0.02 | 0.90±0.05 | 0.90±0.08 | 0.91±0.03 | 0.90±0.06 |
| **AP amplitude average adaptation index** | 0.99±0.00 | 0.99±0.00 | 0.99±0.01 | 0.99±0.00 | 0.99±0.00 | 0.99±0.00 | 0.99±0.01 | 0.99±0.01 | 0.97±0.03 | 0.96±0.03 | 0.98±0.00 | 0.98±0.01 | 0.97±0.02 | 0.97±0.03 | 0.98±0.01 | 0.97±0.02 |
| **AP Fano factor** | 2.97±1.04 | 1.61±0.60 | 1.55±0.93 | 1.21±1.43 | 1.44±1.33 | 0.73±0.34 | 1.20±0.50 | 0.59±0.32 | 1.49±1.46 | 1.82±0.99 | 2.36±1.17 | 2.69±2.71 | 2.01±0.86 | 1.00±0.80 | 1.50±1.21 | 3.82±1.44 |
| **AP coefficient of variation** | 0.25±0.04 | 0.17±0.04 | 0.17±0.07 | 0.14±0.09 | 0.16±0.10 | 0.11±0.03 | 0.15±0.04 | 0.10±0.03 | 0.14±0.07 | 0.17±0.05 | 0.21±0.06 | 0.20±0.11 | 0.19±0.05 | 0.12±0.06 | 0.16±0.07 | 0.27±0.07 |
| **Burstiness** | 0.00±0.00 | 0.00±0.00 | 0.00±0.00 | 0.00±0.00 | 0.00±0.00 | 0.00±0.00 | 0.00±0.00 | 0.00±0.00 | 0.00±0.00 | 0.07±0.15 | 0.00±0.00 | 0.00±0.00 | 0.00±0.00 | 0.00±0.00 | 0.03±0.07 | 0.14±0.15 |
| **Wildness** | 0.00±0.00 | 0.00±0.00 | 0.00±0.00 | 0.00±0.00 | 0.00±0.00 | 0.00±0.00 | 0.00±0.00 | 0.00±0.00 | 0.00±0.00 | 0.00±0.00 | 0.00±0.00 | 0.00±0.00 | 0.00±0.00 | 0.00±0.00 | 0.00±0.00 | 0.00±0.00 |
| **Rebound number of APs** | 0.00±0.00 | 0.00±0.00 | 0.00±0.00 | 0.00±0.00 | 0.00±0.00 | 0.00±0.00 | 0.00±0.00 | 0.00±0.00 | 0.00±0.00 | 0.00±0.00 | 0.00±0.00 | 0.00±0.00 | 0.00±0.00 | 0.00±0.00 | 0.00±0.00 | 0.00±0.00 |
| n | 7 | 6 | 7 | 7 | 6 | 8 | 8 | 9 | 5 | 6 | 6 | 7 | 6 | 6 | 7 | 6 |
| mice used | 5 | 5 | 6 | 5 | 5 | 6 | 6 | 6 | 5 | 5 | 5 | 6 | 5 | 4 | 6 | 5 |
| **Data are presented as Mean ± SD** | | | | | | | | | | | | | | | | |
