## Supplemental table 2 for "Convergent Inhibitory Cortical Circuit Disruption Drives Genetically Distinct Absence Seizures"

| Genes | Probe |
|---|---|
| *Pvalb* | GCGAATTAACCCTCACTAAAGGGTCTGCTCATCCAAGTTGCAGGATGTCGATGACAGACGTGCTC<br>AGCGCTGAGGACATCAAGAAGGCGATAGGAGCCTTTGCTGCTGCAGACTCCTTCGACCACAAAAA<br>GTTCTTCCAGATGGTGGGCCTGAAGAAAAAGAACCCGGATGAGGTGAAGAAGGTGTTCCATATTC<br>TGGACAAAGACAAAAGTGGCTTCATTGAGGAGGATGAGCTGGGGTCCATTCTGAAGGGCTTCTCC<br>TCAGATGCCAGAGACTTGTCTGCTAAAGAAACAAAGACGCTTCTGGCCGCTGGAGACAAGGATGG<br>GGACGGCAAGATTGGGGTTGAAGAATTCTCCACTCTGGTGGCTGAAAGCTAAGTGGCGCTGACT<br>GCTTGGGTCCCCCACCTCTCCATCCCCAACGCCCCATCTCAGCCCTTCTCGCGGCCCTCCTGAG<br>TTTCTGTTCAGTTTGTTTGTGTTATTTTTTACTCCCCCATCCTCTATGGCCCTCGGATGACGCCATT<br>CTTCTGGAAATGCTGGAGAAACAATAAAGGCTGTACCAATCTGACACCACCTGTAGGGAGGACCC<br>AGGCCTGGCAGGGTGTTGGTTTGGCAAGTTTTTTTTCTTTCTTTTTAGGGCAGTGGGGGTATAGTA<br>GAAAAAGTGAGATAAGTCAAAGGACAACGCCCCGATATCTCCTGCCTGCTTGGTACTGAGTGCTC<br>ATGTGGGTCACCTCGTTCAATCTCTGCACCTTTCCCACAAGGAGATGGGGGTGATGGATCGTCCA<br>TCTTAAAGATACAGAAACTGCCTTTTAAAGAGCAGAAGGGAAGGGAAGGGGTTGAGTCCTTCAGG<br>CCCTATAGTGAGTCGTATTACGC |
| *Sst* | GCGAATTAACCCTCACTAAAGGGTGAAGGAGACGCTACCGAAGCCGTCGCTGCTGCCTGAGGAC<br>CTGCGACTAGACTGACCCACCGCGCTCCAGCTTGGCTGCCTGAGGCAAGGAAGATGCTGTCCTG<br>CCGTCTCCAGTGCGCCCTGGCTGCGCTCTGCATCGTCCTGGCTTTGGGCGGTGTCACCGGCGC<br>GCCCTCGGACCCCAGACTCCGTCAGTTTCTGCAGAAGTCTCTGGCGGCTGCCACCGGGAAACAG<br>GAACTGGCCAAGTACTTCTTGGCAGAGCTGCTGTCCGAGCCCAACCAGACAGAGAATGATGCCC<br>TGGAGCCCGAGGATTTGCCCCAGGCAGCTGAGCAGGACGAGATGAGGCTGGAGCTGCAGAGGT<br>CTGCCAACTCGAACCCAGCAATGGCACCCCGGGAACGCAAAGCTGGCTGCAAGAACTTCTTCTG<br>GAAGACATTCACATCCTGTTAGCTTTAATATTGTTGTCCTAGCCAGACCTCTGATCCCTCTCCCCC<br>AAACCCCATATCTCTTCCTTAACTCCTGGCCCCCGATGCTCAACTTGACCCTGCACCCTATAGTGA<br>GTCGTATTACGC |
| *Vip* | GCGAATTAACCCTCACTAAAGGGCCTTCCCTAGAGCAGAACTTCAGCACCCTAGACAGCTGCCAC<br>GAAGCCGGAAAGGCAGCCCTGCCTGAAGGAAACAGCCAAGGAGGCACCGAGATGGAAGCCAGA<br>AGCAAGCCTCAGTTCCTGGCATTCCTGATACTCTTCAGTGTGCTGTTCTCTCAGTCGCTGGCCTG<br>GCCTCTCTTTGGACCACCTTCTGTAGTGAGTAGGCTGGATGACAGGATGCCGTTTGAAGGAGCAG<br>GTGACCCTGACCAAGTCTCTTTAAAAGCAGACTCTGACATCTTGCAGAATCCCTTAGCAGAAAATG<br>GCACACCCTATTATGATGTGTCAAGAAATGCCAGGCATGCTGATGGAGTTTTCACCAGCGATTACA<br>GCAGACTTCTGGGTCAGATTTCTGCCAAAAAATACCTTGAGTCACTCATTGGCAAACGAATCAGCA<br>GCAGCATCTCGGAAGATCCTGTGCCAATCAAACGACACTCTGATGCCGTCTTCACAGATAACTAC<br>ACCCGCCTCAGAAAGCAAATGGCTGTGAAGAAATACCTGAACTCCATCCTGAATGGAAAGAGGAG<br>CAGTGAGGGAGATTCTGCAGACTTTCTTGAAGAGCTGGAGAAATGATGGGAAGAGGCCTCTGGG<br>CAGAGCTGAAATCAGAGAATTCTCGAAGGAAAACAACCACGTGATTACATTATGAGTTCTACATGT<br>CTAATTCAAGAAAAAAACTTCCATAGCAAAACCAAATAAAATGTGTTGTGAATATTGTGGTTTCCTT<br>TATGTAATAACTGTGATGTTTACATTGTAAATATTATTTGAGCATTCTAACATTCATCTGTAGCTCAT<br>GAAATGCTTATAATTTCATATGCTATATATTCTTTCAAAGAAAAGTATATTTAATGATAGGTAGATAC<br>TAGATTAATTGCAATTATCTGAAGCTTTCTGCAAGGGTAGCAATCGAGGAAAATTGATGTCCCTATA<br>GTGAGTCGTATTACGC |
| *Npy* | GCGAATTAACCCTCACTAAAGGGTCACAGAGGCACCCAGAGCAGAGCACCCGCCGCTCAGCGAC<br>GACTGCCCGCCCGCCACGATGCTAGGTAACAAGCGAATGGGGCTGTGTGGACTGACCCTCGCTC<br>TATCTCTGCTCGTGTGTTTGGGCATTCTGGCTGAGGGGTACCCCTCCAAGCCGGACAATCCGGG<br>CGAGGACGCGCCAGCAGAGGACATGGCCAGATACTACTCCGCTCTGCGACACTACATCAATCTC<br>ATCACCAGACAGAGATATGGCAAGAGATCCAGCCCTGAGACACTGATTTCAGACCTCTTAATGAA<br>GGAAAGCACAGAAAACGCCCCCAGAACAAGGCTTGAAGACCCTTCCATGTGGTGATGGGAAATG<br>AAACTTGTTCTCCCGACTTTTCCAAGTTTCCACCCTCATCTCATCTCATCCCCTGAAACCAGTCTGC<br>CTGTCCCACCAATGCATGCCACCACTAGGCTGGACTCCGCCCCATTCCCTATAGTGAGTCGTATT<br>ACGC |
| *Cacng2* | GCGAATTAACCCTCACTAAAGGGCACATATGGAAACTGGAGACCAGAATTTTAGGAAAAGAGATTA<br>AGGCATCTCACTTGGGGGGGTGGGGGGGTGTTTTTTATTTATTTTCCCTTTTTTTAAAAAAATCGCT<br>GCAACTGGAACAGTTTTTTGATCTCAAAAGGCAAGCCTCTCTTCCCGTGTGATCTTTATAATTTACA<br>CACTTTTCCGTGAGCTTTCTTATCTCCCTTTTTTTTATATCTCTCCATATTCTCTATTCACACATATAT<br>CCATTATATTAGTAGTGGAATTACCAATCGCACCCTCACACACACGCTCCTGAGAACCAGAAGTCG<br>GTTGGGTGTTTACCCTATAGTGAGTCGTATTACGC |
| *Nxph2* | GCGAATTAACCCTCACTAAAGGGCCTTCTATTTCTGGCAGTGGACTGCAATTCCTTGTCCAGATTT<br>CAAATTTTAAAAATATATTCAAATGTGAAGTGCATCTGTTTTGGTGAGAGTAACCTGATCTAAAACG<br>TTAAACTGACTGAAACCAGCATTTTCATGTAAGATGGTAACATTCTGGAAATATTGCTTCTCTGTAA<br>GAATCATGAGCAGTGACCATGTGTAAAGGGAGAAACCAAATTGTGTATGCAATAAGCACTGCTCTT<br>CAAATGAGCCTGCCTTTGAAATGCTTTGTAAGGTTTCCTTAGAAAGAGTAGGTTCAGTAAAGTAAC<br>TGAAGAAATGGTATTTAAGAGATGGGTTGTGTTTTTCTGGTGTTGAAACCTTTTTAAAAATTACTAT<br>CTCTTGTTGCCAATTTTAATAGCTACAGTGTCTCAAGAGAATCAAATACAATACCATCTGAGCAGTC<br>TGAGTAGAAAGTGCTTGAAGCACTTTACTATTATGACTCTCATTTGAGTGGTATTGGAACATACCAT<br>TTAGGTATTAGATATGCAAATGTGGATTTAAGTCAACTCAACAATCACGAAGATGATATTTTTCAAT<br>GTTGTGGGATTATTTTTTAAATTCTTATTCATATGAAAGTTTGGGACTTCATTCCATTATTTAATAGT<br>CCAATTTTATATATACATTTGTGGAACAACTCTGAAATTGAAAGAAAGTATTTCTGGGCAGAAAATT<br>CCACAGTCATGGGTTCATTATATGTAAGGCCTCAGATTAGGTTTTCCTTGTGTTTGAGTAATCACCA<br>GAATGAGACTAAGAATTCTGTGTGGAAATAATGAGTTATATTTCAGTATTTCTAAATATTAAGTGATT<br>ATATTTATAATGTCTGTATTACTGAGCTTTTGAGATTACTTTACTCTCAGTGTACTCATCTTCGAGAA<br>AAGTTGAATGGCCATCTTTATGATGTACCCTGGCACTCCCTATAGTGAGTCGTATTACGC |
| *Ntsr1* | GCGAATTAACCCTCACTAAAGGGTATTCTTGCCCCTTATCCCTCTCCTACCTCCACCTTTAAAAACA<br>GAAAAGAGGCGGTTTCTCTCCTGGCCCTACAAAAGGCCTTTAACAAGAAGAAATTAGCACTCACC<br>AAGGACGGTCTCCTTTGTTCCCAGACTAATGGATGTTTTAGAAGCAAGAAATGAAAGCACCACATT<br>GGGCCTGGACAGATGAGCTGTTGTAACCATAACAGCACCTTGGAACTGCACTGGCAGGGCGAGA<br>CAGTGTCTGATGTTGGACTTGGGTTCAGAGACACAGGCTCCATCTCAGCCCCTTTATGCCTCTAC<br>TCCTGCCCTGGTCCAGCAGATACCATAGCACTCTCTGAGCCTTATGTCCAGACCCTTTCTTGGCA<br>GGCTGGTACACTGCCCTTCCCAGAGTGGTCCAGAGAAGCCCCAAAATTACTGTGTAAGGTCCAG<br>GGCCCACAGCTGGAAGCTGTGGGAATCCATGCCACACCTGGATGGCTATGGCCCACTAAAGAGA<br>CCCCAATTCCCACATGCCCAGGAAGGATAAAAGGGCTGGCCTGGAATCAACACAGGACAAGCTT<br>CAAGTGGCTTTGCACGGGGACCTTCACCTCTTGGATCTGCAGAAGGATGGAAGATAAGTGGGGT<br>CCAGCCTCCCCAGTCCAGGTGGCTTTGCTGGGGACATGCATGGTTCTGCGTCTCATATACAGATG<br>TATAGACTAAGGTCTGACAGCAGCTGGCTTCAGGCTGGGACTTCTGAGAAGGGGCACTGAGCCA<br>GGTCCTCAGCAAGATGCCAGTGTCTGAGACTCCACAGGTAAGGGGAGGCCAAGCACCCCAGCAT<br>CTAGCTGGCCAGCAGCCCTGGACTGAGGCATATGTCAATTTGTTAACCAGCCGCCAAGCAGCCA<br>CCCTGGCCCAGGTTCTAGGCGCCTGGAAAAGCAGGCACCCTTCATCTCTGGGAGCTGTCTCCCT<br>ATAGTGAGTCGTATTACGC |
| *Cacna1a* | GCGAATTAACCCTCACTAAAGGGAATGTCCTCATAGGGTTGCTTGCAAGAGACTCCTGCGGATGG<br>ACCTGCCGGTAGCAGATGACAACACAGTTCACTTCAACTCCACCCTGATGGCTCTGATCCGAACC<br>GCCCTTGATATCAAAATTGCCAAGGGTGGAGCTGACAAGCAGCAGATGGATGCAGAGCTTCGCAA<br>GGAGATGATGGCCATTTGGCCCAACCTGTCTCAGAAGACCCTGGATCTGCTGGTCACACCTCACA<br>AGTCCACGGACCTGACAGTGGGTAAGATCTACGCAGCCATGATGATCATGGAGTATTACCGGCAG<br>AGCAAGGCCAAGAAACTGCAGGCCATGCGAGAGGAGCAGAACCGGACACCACTCATGTTCCAGC<br>GCATGGAGCCTCCATCACCAACACAGGAGGGAGGACCCAGCCAGAACGCCCTTCCCTCCACTCA<br>GCTGGACCCAGGAGGAGGCCTGATGGCTCACGAAGGCGGCATGAAGGAGAGCCCGTCCTGGGT<br>GACCCAGCGGGCCCAAGAGATGTTCCAGAAGACTGGCACCTGGAGCCCAGAGCGAGGGCCACC<br>CATCGACATGCCTAACAGCCAGCCCAACTCCCAGTCTGTGGAGATGCGAGAAATGGGTACTGATG<br>GCTACTCAGACAGCGAACACTACCTCCCCATGGAAGGCCAGACCAGGGCCGCCTCCATGCCCCG<br>CCTCCCAGCAGAGAACCAGAGGAGAAGGGGCCGGCCACGTGGAAATGACCTCAGTACCATCTCT<br>GATACCAGCCCCATGAAGCGCTCAGCCTCTGTGCTGGGACCCAAAGCCCGGCGACTGGATGACT<br>ACTCACCCCTATAGTGAGTCGTATTACGC |
| *tdTomato* | GCGAATTAACCCTCACTAAAGGGATCAAAGAGTTCATGCGCTTCAAGGTGCGCATGGAGGGCTCC<br>ATGAACGGCCACGAGTTCGAGATCGAGGGCGAGGGCGAGGGCCGCCCCTACGAGGGCACCCA<br>GACCGCCAAGCTGAAGGTGACCAAGGGCGGCCCCCTGCCCTTCGCCTGGGACATCCTGTCCCC<br>CCAGTTCATGTACGGCTCCAAGGCGTACGTGAAGCACCCCGCCGACATCCCCGATTACAAGAAG<br>CTGTCCTTCCCCGAGGGCTTCAAGTGGGAGCGCGTGATGAACTTCGAGGACGGCGGTCTGGTGA<br>CCGTGACCCAGGACTCCTCCCTGCAGGACGGCACGCTGATCTACAAGGTGAAGATGCGCGGCAC<br>CAACTTCCCCCCCGACGGCCCCGTAATGCAGAAGAAGACCATGGGCTGGGAGGCCTCCACCGA<br>GCGCCTGTACCCCCGCGACGGCGTGCTGAAGGGCGAGATCCACCAGGCCCTGAAGCTGAAGGA<br>CGGCGGCCACTACCTGGTGGAGTTCAAGACCATCTACATGGCCAAGAAGCCCGTGCAACTGCCC<br>GGCTACTACTACGTGGACACCAAGCTGGACATCACCTCCCACAACGAGGACTACACCATCGTGGA<br>ACCCTATAGTGAGTCGTATTACGC |
| *Tac1* | GCGAATTAACCCTCACTAAAGGGTCGAACATGAAAATCCTCGTGGCCGTGGCGGTCTTTTTTCTC<br>GTTTCCACTCAACTGTTTGCAGAGGAAATCGATGCCAACGATGATCTAAATTATTGGTCCGACTGG<br>TCCGACAGTGACCAGATCAAGGAGGCAATGCCGGAGCCCTTTGAGCATCTTCTGCAGAGAATCG<br>CCCGAAGACCCAAGCCTCAGCAGTTCTTTGGATTAATGGGCAAGCGGGATGCTGATTCCTCAGTT<br>GAAAAACAAGTGGCCCTGTTAAAGGCTCTTTATGGACATGGCCAGATCTCTCACAAAAGGCATAAA<br>ACAGATTCCTTTGTTGGACTAATGGGCAAAAGAGCTTTAAATTCTGTCGCTTATGAAAGAAGCGCA<br>ATGCAGAACTACGAAAGAAGACGTAAATAAACCCCTGAACGCACTATCTATTCATCTTCATCTGTG<br>TCAGTGAGCAGTGAACGGTAAAATAAAAATGTGCGCTATGAGGAATGATTATTTATTTAACACATGT<br>TGTTGTGAGTGAAAAACTCAAAAAAGAAGTGTTTATTTTTTCATATTGTGCCAATAAGCCTTGTAATT<br>CTAATGTGGTGACCTCCCCAAAAGTAGAAATTAGTTGTAACTTCAGCAAAGCACAGTGATGAAGGA<br>GCTGTCCAAGCTTGCCAGTGACGCAGTCTCCAAAGAAAGGGCCCTTCTGTGAGCCAGCGCAGGC<br>AGCTGCTGCTGGAAGCAGAGACTCCTGTGCGTCTTCTCTCGCGCTACCCCTGGTTCTGCTTTCAT<br>GCTGATAATGTACTGAGACTTCGGTATTTGACTCTATTTGTATCCTAGCAGCATGTTTCCTGTTTCG<br>TGACTATATAGAGATGTTTTGAAAAAGTTTCAATGTAATTCTCTGGTCTTCAGTCATTGTATGATGT<br>GCCCTATAGTGAGTCGTATTACGC |
