## supplemental figures for "Convergent Inhibitory Cortical Circuit Disruption Drives Genetically Distinct Absence Seizures"

### Supplemental Information

#### Extended Data Figures

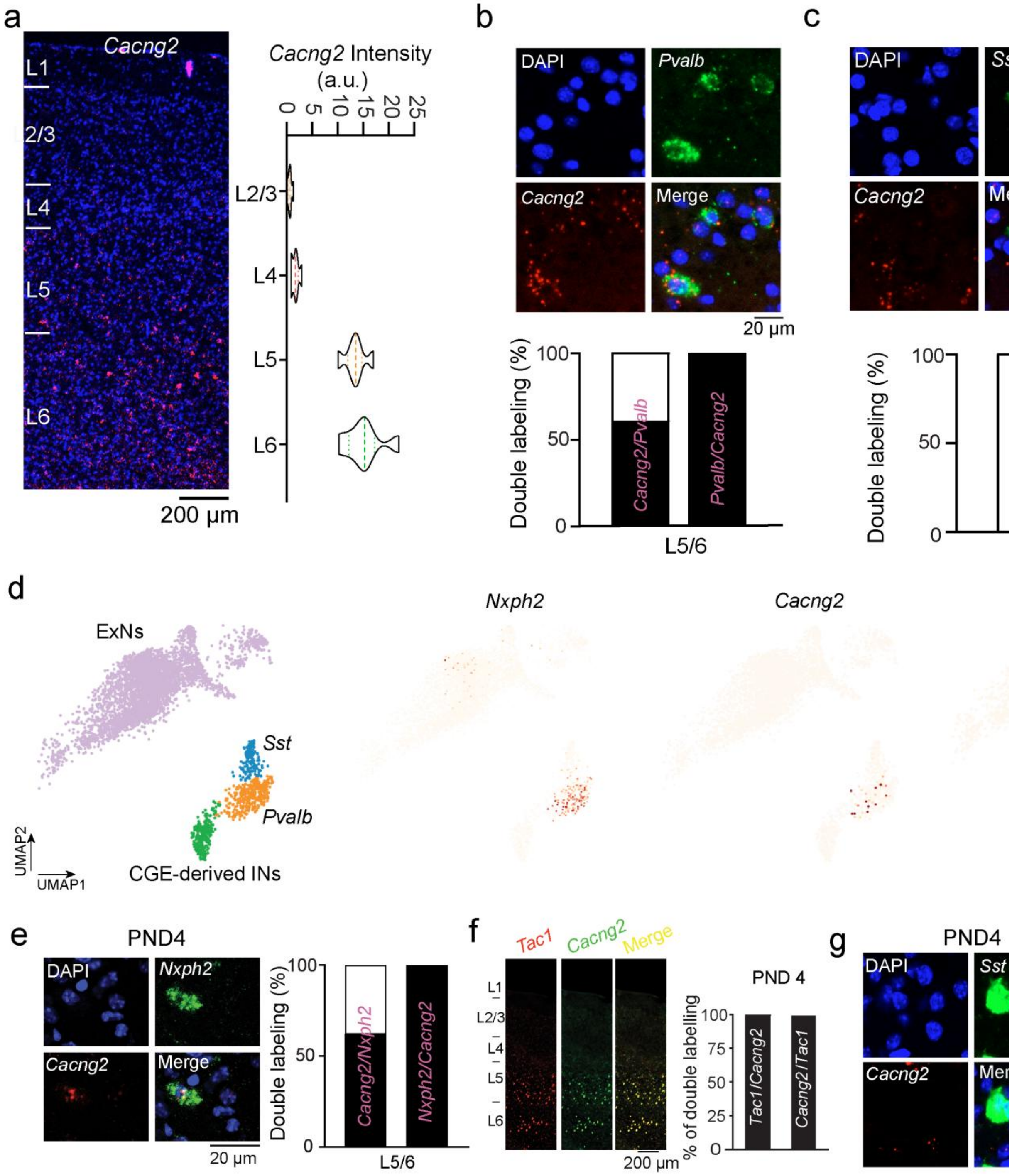


**Extended Data Fig. 1 | *Cacng2* expression in the mouse neocortex. Related to Fig. 1. a,** Left: A representative micrograph of FISH staining for *Cacng2* in a parasagittal section of adult mouse

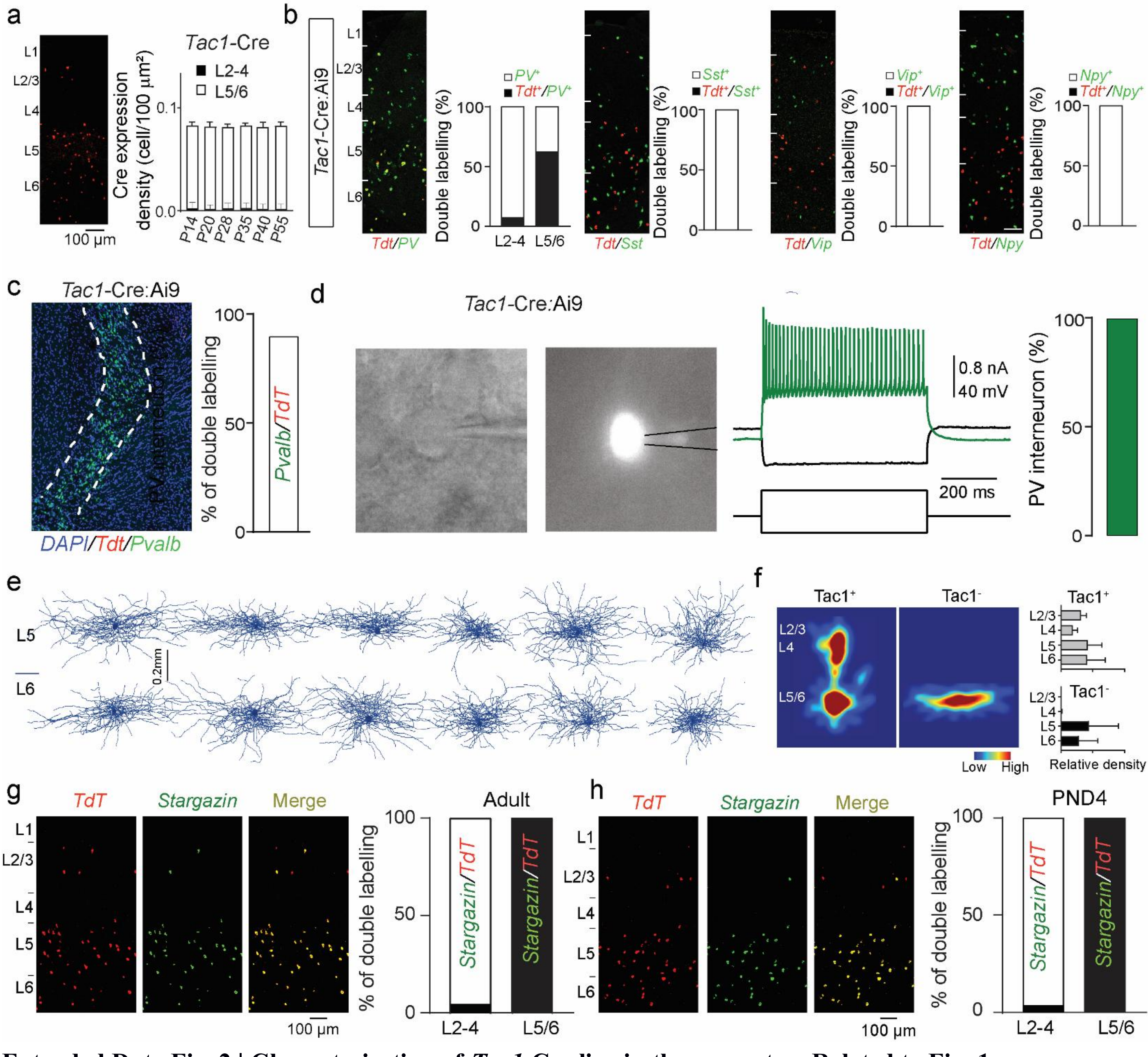


**Extended Data Fig. 2 | Characterization of *Tac1*-Cre line in the neocortex. Related to Fig. 1.** **a,** Left: Micrograph shows RNA FISH detection of Cre mRNA in the *Tac1*-Cre line at P55. The representative image is the parasagittal S1 section from at least two brain-wide experiments. Right: Developmental characterization of Cre expression in the *Tac1*-Cre line. Quantification in layers 2-4 was performed across 6 consecutive sections from 6 mice at each age. Quantification in layers 5/6 was performed across 6 consecutive sections from 6 mice at P14, P20, and P55, across 5 consecutive sections from 6 mice at P28 and P40, and across 4 consecutive sections from 6 mice at P35. **b,** Representative double FISH staining for *Tdt* mRNA (red) together with *Pvalb*, *Sst*, *Vip*, or *Npy* mRNA (green) in parasagittal S1 sections of adult *Tac1*-Cre:Ai9 mice. Bar graphs depict the proportion of $Tdt^{+}/Pvalb^{+}$ cells within total $Pvalb^{+}$ cells in superficial layers (L2-4) and layers 5/6 (L5/6). For *Sst*, *Vip*, and *Npy*, bar graphs depict the proportion of $Tdt^{+}/Sst^{+}$, $Tdt^{+}/Vip^{+}$, and $Tdt^{+}/Npy^{+}$ cells within total $Sst^{+}$, $Vip^{+}$, and $Npy^{+}$ cells, respectively. Quantification for *Pvalb* was performed across 7 consecutive sections from 6 mice; for *Sst*, across 6 consecutive sections from 6 mice; for *Vip*, across 6 consecutive sections from 5 mice; and for *Npy*, across 7 consecutive sections from 6 mice. **c,** Left: Representative micrograph showing double FISH staining for *Tdt* mRNA (red) and *Pvalb* mRNA (green) in the thalamic reticular nucleus (RTN) of adult *Tac1*-Cre:Ai9 mice. Right: Bar graph depicting the proportion of double-labeled cells ($Pvalb^{+}/Tdt^{+}$) within the total single-labeled $Tdt^{+}$ cells in the RTN, counted across 7 consecutive sections from 8 mice. **d,** Representative whole-cell recording and firing pattern of a *Tac1*-Cre-labeled interneuron in deep layers of S1. Left: Brightfield image of the recorded neuron. Middle: Fluorescence image of the labeled cell. Right: Representative firing pattern in response to depolarizing current injection. Bar graph shows the proportion of recorded deep-layer *Tac1*-Cre-labeled neurons displaying fast-spiking properties characteristic of *Pvalb* interneurons (n = 20 cells from 8 mice). **e,** Representative morphologies of $Tac1^{-}$ interneurons in layers 5 and 6 of S1. Morphological reconstructions were obtained from n = 12 cells from 8 mice. **f,** Left: Axon density maps of $Tac1^{+}$ and $Tac1^{-}$ interneurons. Middle: Representative layer-wise axon density distributions. Right: Quantification of relative axon density across cortical layers for $Tac1^{+}$ and $Tac1^{-}$ interneurons. Morphological reconstructions were obtained from n = 12 cells per group from 8 mice. **g,** Left: Representative immunofluorescence micrographs showing TdT (red), stargazin (green), and merged signals in the S1 cortex of adult *Tac1*-Cre:Ai9 mice. Right: Bar graphs depicting the proportion of double-labeled cells ($stargazin^{+}/Tdt^{+}$) within the total single-labeled $Tdt^{+}$ cells in superficial layers (L2–4) and layers 5/6 (L5/6), counted across 8 sections from 8 mice. In L2–4, 2 of 42 $TdT^{+}$ cells were $stargazin^{+}/TdT^{+}$ double-labeled cells (2/42, 4.8%). In L5/6, all $TdT^{+}$ cells were $stargazin^{+}/TdT^{+}$ double-labeled cells (329/329, 100%). **h,** Left: Representative immunofluorescence micrographs showing TdT (red), stargazin (green), and merged signals in the S1 cortex of *Tac1*-Cre:Ai9 mice at postnatal day 4 (PND4). Right: Bar graphs depicting the proportion of double-labeled cells ($stargazin^{+}/Tdt^{+}$) within the total single-labeled $Tdt^{+}$ cells in superficial layers (L2-4) and layers 5/6 (L5/6), counted across 9 sections from 7 mice. In L2–4, 2 of 47 $TdT^{+}$ cells were $stargazin^{+}/TdT^{+}$ double-labeled cells (2/47, 4.3%). In L5/6, all $TdT^{+}$ cells were $stargazin^{+}/TdT^{+}$ double-labeled cells (367/367, 100%).

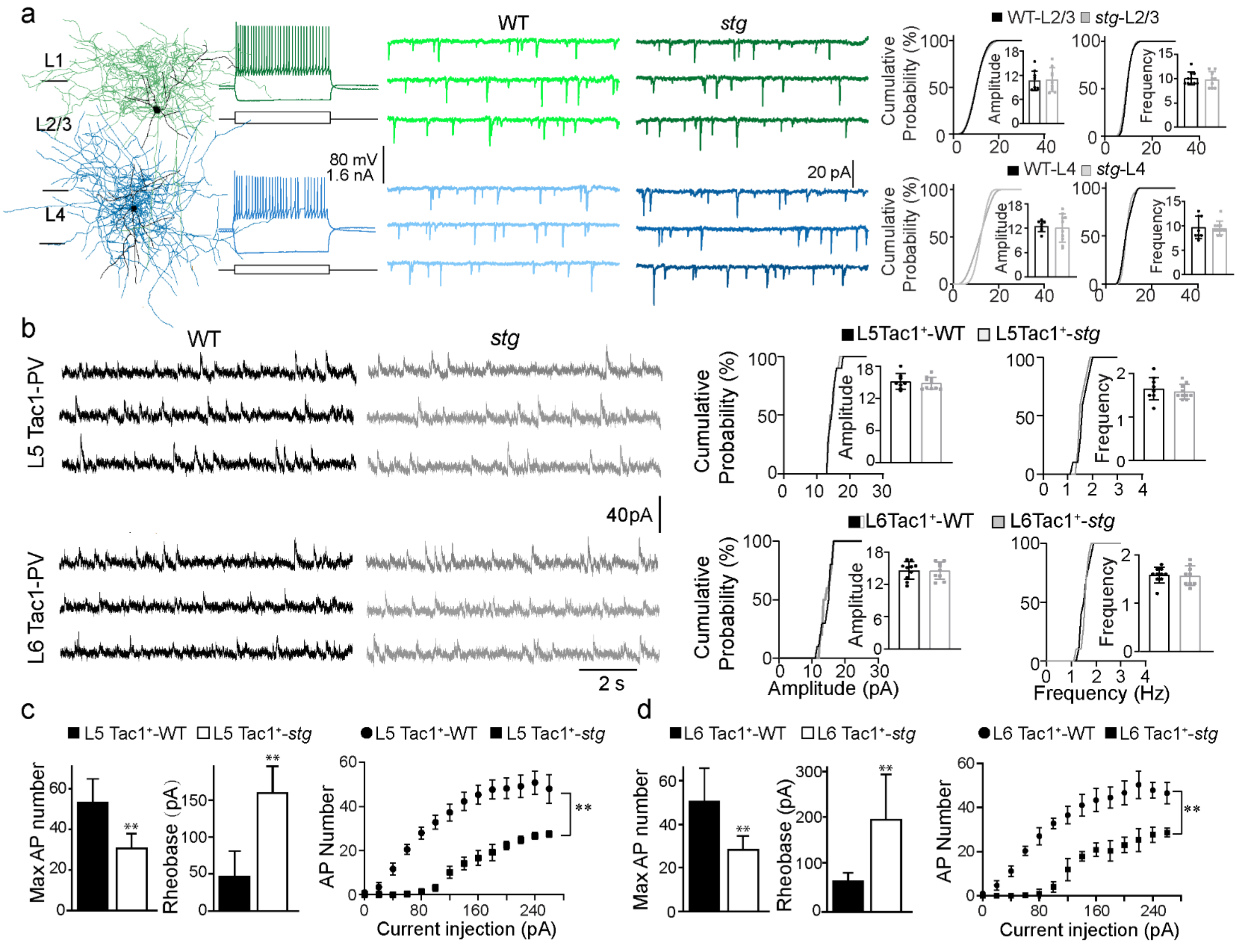

a
L1
L2/3
L4
WT
stg
80 mV
1.6 nA
20 pA
WT-L2/3
stg-L2/3
WT-L4
stg-L4
Cumulative Probability (%)
Amplitude
Frequency
b
L5 Tac1-PV
L6 Tac1-PV
40pA
2 s
L5Tac1+-WT
L5Tac1+-stg
L6Tac1+-WT
L6Tac1+-stg
Amplitude (pA)
Frequency (Hz)
c
L5 Tac1+-WT
L5 Tac1+-stg
Max AP number
Rheobase (pA)
AP Number
Current injection (pA)
d
L6 Tac1+-WT
L6 Tac1+-stg

**Extended Data Fig. 3 | Characterization of Tac1$^+$ *Pvalb* interneurons in *stg*. Related to Fig. 2.** **a,** Spontaneous excitatory inputs onto superficial *Pvalb* interneurons (L2/3 and L4). Left: Identification of *Pvalb* interneurons in L2/3 and L4 by their typical morphology and fast-spiking firing patterns. Middle: Representative traces of spontaneous excitatory postsynaptic currents (sEPSCs). Right: Cumulative probability plots and inset bar graphs summarize sEPSC amplitude and frequency in WT and *stg* mice. Cells for WT and *stg*, respectively: L2/3 (n = 10, 9) and L4 (n = 8, 10). Data are color-coded by layer and were analyzed using Student's t-test. **b,** NMDA receptor-mediated EPSCs in deep-layer (L5/6) Tac1$^+$ *Pvalb* interneurons. Left: Representative NMDA-EPSC traces. Right: Cumulative probability plots and inset bar graphs summarize NMDA-EPSC amplitude and frequency in WT and *stg* mice. Cells for WT and *stg*, respectively: L5 (n = 10, 10) and L6 (n = 10, 10). Statistical comparisons were performed using Student's *t*-test. **c,** Intrinsic excitability of L5 Tac1$^+$ *Pvalb* interneurons. Left: Bar graphs showing the maximum number of action potentials (APs) and rheobase in L5 *Tac1*$^+$ interneurons from WT and *stg* mice. Right: Firing response to increasing current injections (F-I curve). For both panels, n = 6 neurons per genotype. Maximal AP number and rheobase were compared using Student's t-test (maximal AP number: $t(10) = 4.466$, **$p < 0.01$; rheobase: $t(10) = 5.519$, **$p < 0.01$), and the F-I curve was analyzed by two-way ANOVA ($F(1, 10) = 12.10$, **$p < 0.01$). See Extended Data Table 1 for additional electrophysiological data. **d,** Intrinsic excitability of L6 Tac1$^+$ *Pvalb* interneurons. Left: Bar graphs showing the maximum number of APs and rheobase in L6 Tac1$^+$ interneurons from WT and *stg* mice. Right: Firing response to increasing current injections (F-I curve). For both panels, n = 6 neurons per genotype. Maximal AP number and rheobase were compared using Student's t-test (maximal AP number: $t(10) = 3.541$, **$p < 0.01$; rheobase: $t(10) = 3.472$, **$p < 0.01$), and the F-I curve was analyzed by two-way ANOVA ($F(1, 10) = 31.69$, **$p < 0.01$). See Extended Data Table 1 for additional electrophysiological data.

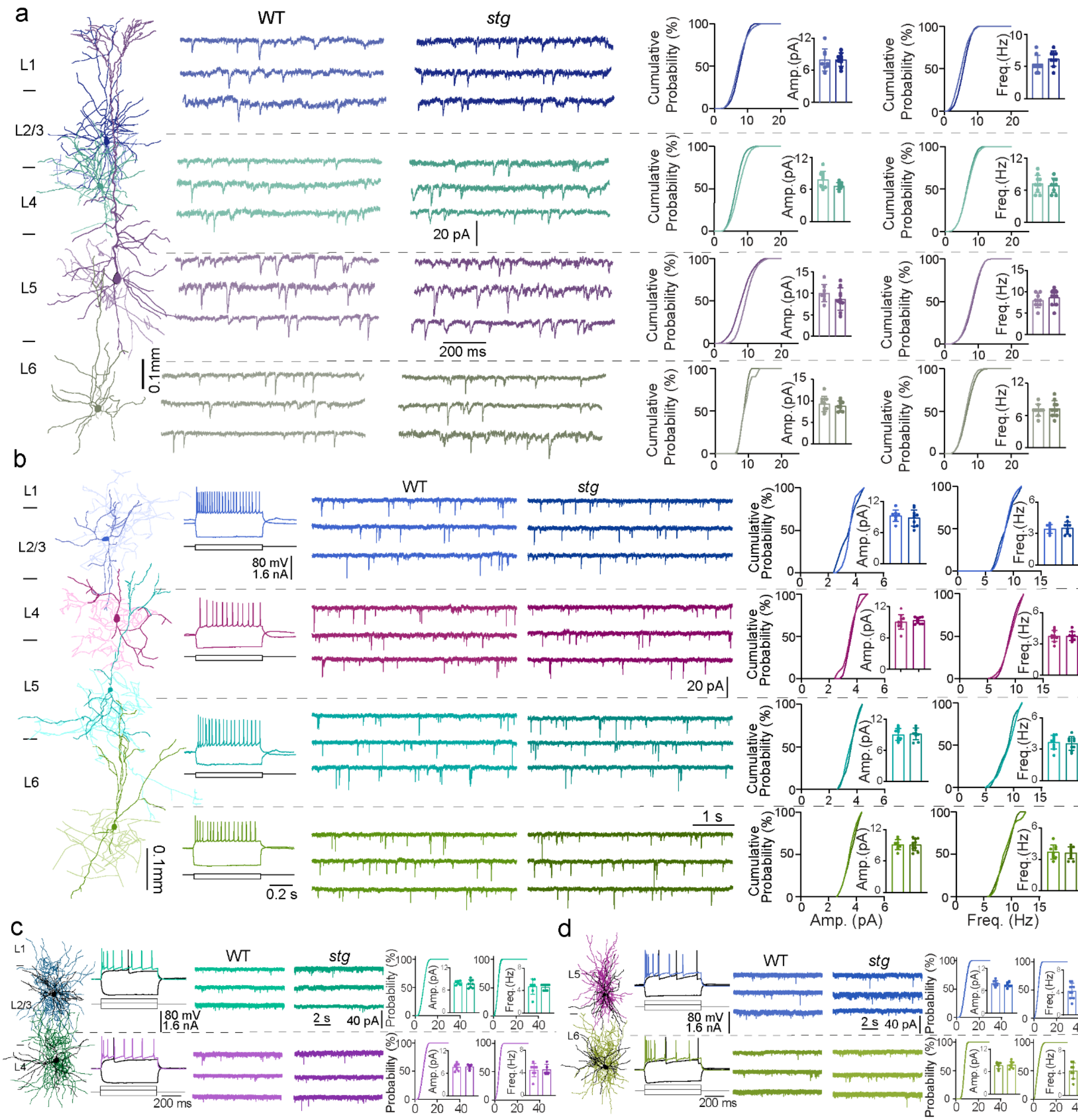


**Extended Data Fig. 4 | Synaptic transmission of principal cells (PCs), *Vip* interneurons, and neurogliaform cells across cortical layers in S1 of WT and *stg* mice. Related to Fig. 3. a,** Spontaneous excitatory postsynaptic currents (sEPSCs) in principal cells (PCs) across layers and genotypes. From left to right: Representative morphology of a recorded PC, sample sEPSC traces from each cortical layer in wild-type (WT) and stargazin mutant (*stg*) mice, and cumulative probability plots with inset bar graphs summarizing sEPSC amplitude and frequency. Cells for WT and *stg*, respectively: L2/3 (n = 8, 9), L4 (n = 10, 9), L5 (n = 8, 10), L6 (n = 9, 10). **b,** sEPSCs in *Vip* interneurons. From left to right: Representative morphology and firing patterns, sample sEPSC traces, and cumulative probability plots with inset bar graphs for sEPSC amplitude and frequency. Cells for WT and *stg*, respectively: L2/3 (n = 8, 10), L4 (n = 10, 9), L5 (n = 10, 9), L6 (n = 9, 10). **c,d,** sEPSCs in neurogliaform cells (NGCs) in superficial (c) and deep (d) layers. In both panels: representative morphology and firing patterns (left), sample sEPSC traces (middle), and summary plots for sEPSC amplitude and frequency (right). Cells for WT and *stg*, respectively: L2/3 (n = 7, 8), L4 (n = 8, 5), L5 (n = 10, 9), L6 (n = 8, 10). Statistical comparisons were performed using Student's *t*-test.

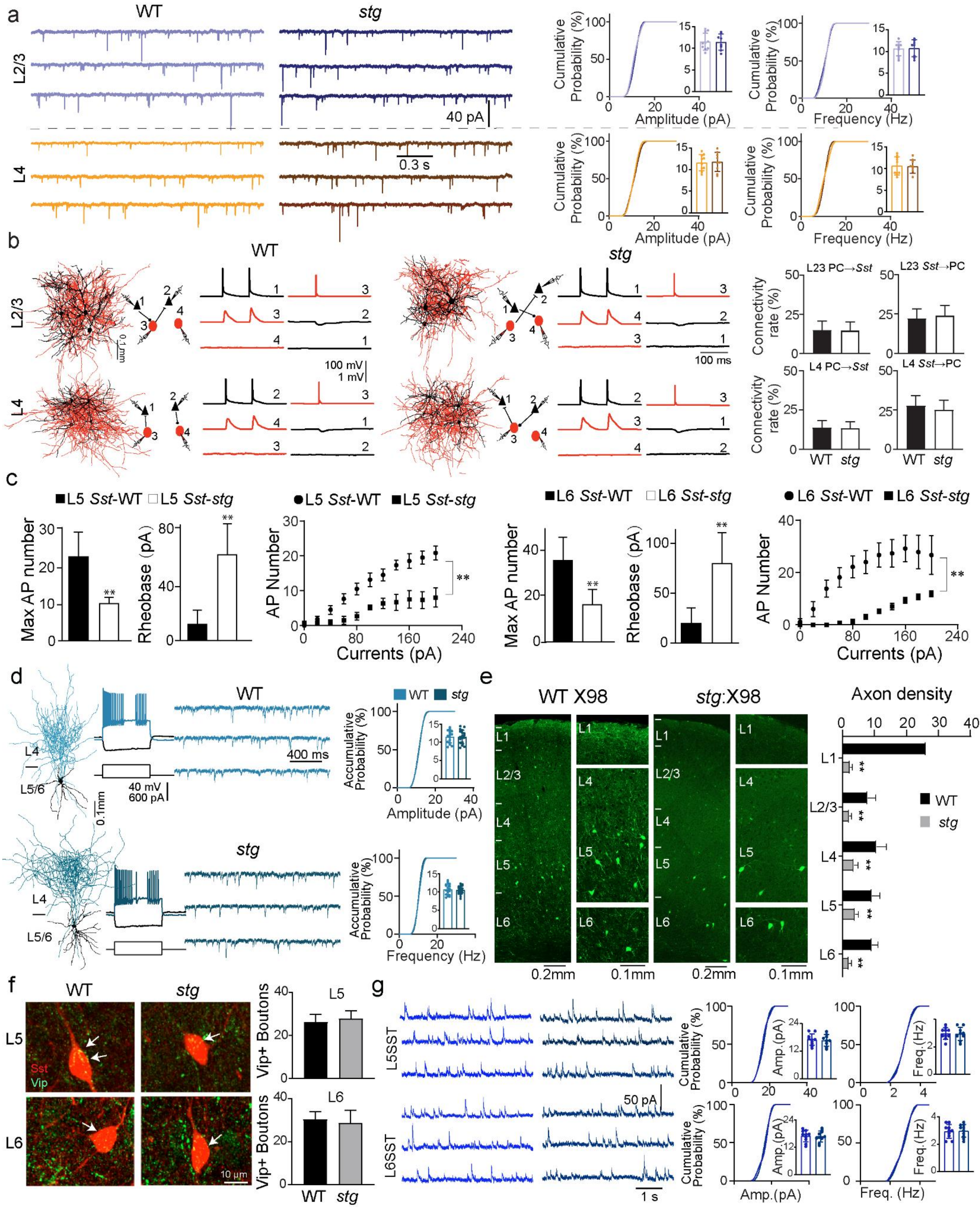
a
WT
stg
L2/3
L4
40 pA
0.3 s
Cumulative Probability (%)
Amplitude (pA)
Frequency (Hz)
b
L2/3 PC→Sst
L23 Sst→PC
L4 PC→Sst
L4 Sst→PC
Connectivity rate (%)
100 mV
1 mV
100 ms
0.1 mm
c
L5 Sst-WT
L5 Sst-stg
L6 Sst-WT
L6 Sst-stg
Max AP number
Rheobase (pA)
AP Number
Currents (pA)
d
L4
L5/6
0.1mm
40 mV
600 pA
400 ms
Accumulative Probability (%)
e
WT X98
stg:X98
Axon density
L1
L2/3
L4
L5
L6
0.2mm
0.1mm
f
Sst
Vip
Vip+ Boutons
10 μm
g
L5SST
L6SST
50 pA
1 s
Amp.(pA)
Freq.(Hz)
Freq. (Hz)

**Extended Data Fig. 5 | Characterization of *Sst* interneurons in WT and *stg* mice. Related to Fig. 3. a,** Sample traces of spontaneous excitatory postsynaptic currents (sEPSCs) recorded in *Sst* interneurons across cortical layers and genotypes (WT vs. *stg*). Data are color-coded by layer. Cumulative probability plots and inset bar graphs summarize sEPSC amplitude and frequency in *Sst* interneurons from each layer. Cells for WT and *stg*, respectively: L2/3 (n = 10, 8) and L4 (n = 10, 9). Statistical comparisons were performed using Student's *t*-test. **b,** Left: Representative examples of simultaneous 4-cell recordings from principal excitatory cells (PCs, black triangles) and Sst interneurons (SSTs, red circles) in WT and *stg* mice. Reconstructed morphologies of recorded neurons are shown together with schematic connectivity diagrams and representative unitary synaptic traces. Recorded neurons were located close to each other (generally less than 150 µm apart). Vertical scale bars indicate the amplitudes of injected currents in nA, action potentials (APs) in mV, and unitary excitatory or inhibitory postsynaptic potentials (uEPSPs or uIPSPs) in mV. Right: Connection probabilities of PC→*Sst* and *Sst*→PC across cortical layers and genotypes (WT vs. *stg*). PC→*Sst* in L2/3: 6/40 in WT and 6/41 in *stg*; *Sst*→PC in L2/3: 10/45 in WT and 10/40 in *stg*. PC→*Sst* in L4: 8/57 in WT and 8/61 in *stg*; *Sst*→PC in L4: 13/47 in WT and 12/48 in *stg*. **c,** Intrinsic excitability of L5 and L6 *Sst* interneurons. Left panels: Bar graphs showing the maximum number of action potentials (APs) and rheobase in L5 or L6 *Sst* neurons from WT and *stg* mice. Right panels: Firing responses to increasing current injections (F-I curves). For all panels, n = 6 neurons per genotype. For L5 *Sst* interneurons, maximal AP number and rheobase were compared using Student's *t*-test (maximal AP number: $t(10) = 4.815$, $^{**}p < 0.01$; rheobase: $t(10) = 5.010$, $^{**}p < 0.01$), and the F-I curve was analyzed by two-way ANOVA ($F(1, 10) = 25.21$, $^{**}p < 0.01$). For L6 *Sst* interneurons, maximal AP number and rheobase were compared using Student's *t*-test (maximal AP number: $t(10) = 4.025$, $^{**}p < 0.01$; rheobase: $t(10) = 4.380$, $^{**}p < 0.01$), and the F-I curve was analyzed by two-way ANOVA ($F(1, 10) = 13.75$, $^{**}p < 0.01$). See Table S1 for additional electrophysiological data. **d,** Quasi-fast-spiking *Sst* interneurons. Left: Representative morphologies. Middle: Representative firing patterns and sample sEPSC traces. Right: Summary plots of sEPSC amplitude and frequency. Cells for WT and *stg*, respectively: n = 16 and n = 19. **e,** Left: Representative micrographs showing axonal distributions in X98 mice. In WT mice, X98-labeled axons span all cortical layers (L1-L6), whereas in *stg*:X98 mice, axonal loss is observed across these layers. Right: Bar graphs quantify axon density across cortical layers (L1, L2/3, L4, L5, and L6) in WT and *stg* mice. For L1: n = 6 sections from 6 WT mice and n = 5 sections from 5 *stg* mice; for L2/3: n = 6 sections from 5 WT mice and n = 6 sections from 5 *stg* mice; for L4: n = 5 sections from 5 WT mice and n = 5 sections from 5 *stg* mice; for L5: n = 5 sections from 4 WT mice and n = 6 sections from 5 *stg* mice; for L6: n = 6 sections from 5 WT mice and n = 6 sections from 6 *stg* mice. Statistical comparisons were performed using Student's *t*-test (L1: $t(9) = 18.76$, $^{**}p < 0.01$; L2/3: $t(10) = 5.614$, $^{**}p < 0.01$; L4: $t(8) = 4.780$, $^{**}p < 0.01$; L5: $t(9) = 4.813$, $^{**}p < 0.01$; L6: $t(10) = 8.521$, $^{**}p < 0.01$. **f,** Left: Representative micrographs showing immunostaining for Vip in *Sst*-Cre:Ai9 (WT) and *stg*:*Sst*-Cre:Ai9 (*stg*) mice. Arrows denote $Vip^{+}$ boutons overlapping with the soma of *Sst* interneurons. Right: Bar graphs showing the average number of $Vip^{+}$ boutons on each *Sst* interneuron in L5 (upper) and L6 (bottom) from WT and *stg* mice (n = 5 cells for L5-WT, n = 6 cells for L5-*stg*, n = 5 cells for L6-WT, and n = 6 cells for L6-*stg*). **g,** Left: Sample traces of spontaneous inhibitory postsynaptic currents (sIPSCs) recorded in L5/6 *Sst* interneurons from WT and *stg* mice. Right: Cumulative probability plots and inset bar graphs summarize sIPSC amplitude and frequency in L5 (upper) and L6 (lower) *Sst* interneurons. Cells for WT and *stg*, respectively: L5 (n = 10, 9) and L6 (n = 10, 10). Statistical comparisons were performed using Student's *t*-test.

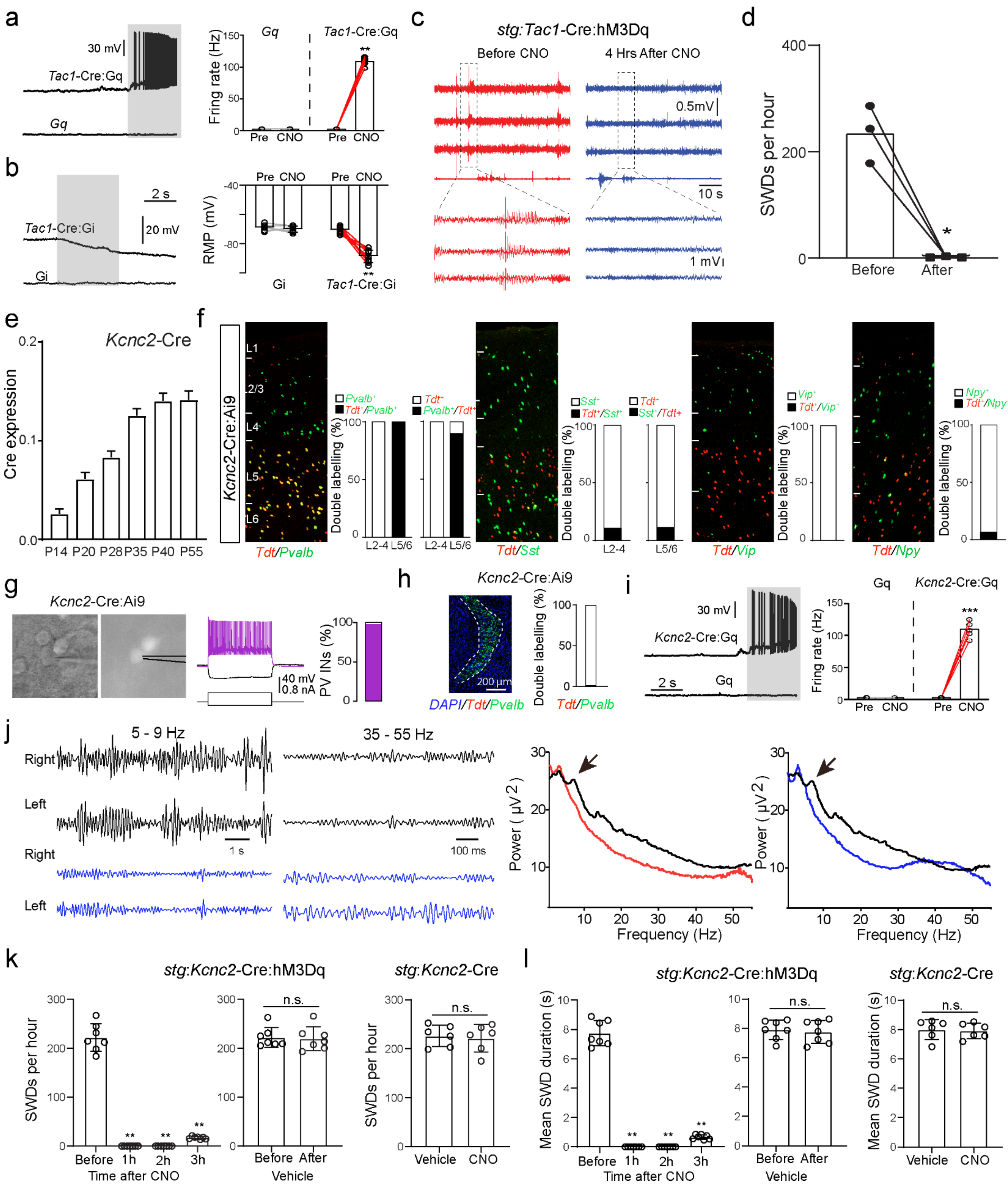
a
30 mV
Tac1-Cre:Gq
Gq
Fring rate (Hz)
Gq
Tac1-Cre:Gq
Pre CNO Pre CNO
b
2 s
20 mV
Tac1-Cre:Gi
Gi
RMP (mV)
Pre CNO Pre CNO
Gi
Tac1-Cre:Gi
c
stg:Tac1-Cre:hM3Dq
Before CNO
4 Hrs After CNO
0.5mV
10 s
1 mV
d
SWDs per hour
Before
After
e
Kcnc2-Cre
Cre expression
P14 P20 P28 P35 P40 P55
f
Kcnc2-Cre:Ai9
L1
L2/3
L4
L5
L6
Tdt/Pvalb
Pvalb+
Tdt+/Pvalb+
Tdt+
Pvalb+/Tdt+
Double labelling (%)
L2-4 L5/6 L2-4 L5/6
Tdt/Sst
Sst+
Tdt+/Sst+
Tdt+
Sst+/Tdt+
L2-4
L5/6
Tdt/Vip
Vip+
Tdt+/Vip+
Tdt/Npy
Npy+
Tdt+/Npy
g
Kcnc2-Cre:Ai9
40 mV
0.8 nA
PV INs (%)
h
Kcnc2-Cre:Ai9
200 μm
DAPI/Tdt/Pvalb
Double labelling (%)
Tdt/Pvalb
i
30 mV
Kcnc2-Cre:Gq
2 s
Gq
Fring rate (Hz)
Gq
Kcnc2-Cre:Gq
Pre CNO Pre CNO
j
5 - 9 Hz
35 - 55 Hz
Right
Left
Right
Left
1 s
100 ms
Power ( μV2 )
Frequency (Hz)
Power ( μV2 )
Frequency (Hz)
k
stg:Kcnc2-Cre:hM3Dq
stg:Kcnc2-Cre
SWDs per hour
n.s.
n.s.
Before 1h 2h 3h
Time after CNO
Before After
Vehicle
SWDs per hour
Vehicle CNO
l
stg:Kcnc2-Cre:hM3Dq
stg:Kcnc2-Cre
Mean SWD duration (s)
n.s.
n.s.
Before 1h 2h 3h
Time after CNO
Before After
Vehicle
Mean SWD duration (s)
Vehicle CNO

**Extended Data Fig. 6 | Validation of chemogenetic manipulations and characterization of the Kcnc2-Cre line. Related to Fig. 4. a,** Representative membrane potential traces from recorded L5/6 neurons in *Tac1*-Cre:Gq mice in response to clozapine-N-oxide (CNO, 10 µM; gray shading). Right, summary of firing rates before and after CNO administration. n = 8 cells from 3 *Tac1*-Cre:Gq mice; paired t-test, t(7) = 4.562, $^{**}p < 0.01$. **b,** Representative membrane potential traces from recorded L5/6 neurons in *Tac1*-Cre:Gi mice in response to clozapine-N-oxide (CNO, 10 µM; gray shading). Right, summary of membrane potential before and after CNO administration. n = 9 cells from 6 *Tac1*-Cre:Gi mice; paired t-test, t(8) = 3.401, $^{**}p < 0.01$. **c,** To test whether orally administered CNO acts rapidly in the brain, CNO was provided to *stg*:Tac1-Cre:hM3Dq mice and seizure activity was monitored. Representative EEG traces recorded before (red) and 4 h after CNO drinking (40 mg/L; blue) show a marked reduction in spike-wave discharges (SWDs). Bottom, expanded traces. **d,** Quantification of SWDs per hour within a 1-hour EEG window (approximately 12:00-13:00) demonstrates a significant reduction in discharge frequency following CNO administration. n = 3 mice; paired t-test, t(2) = 7.311, $^{*}p < 0.05$. **e,** Bar graph showing Cre-dependent tdTomato expression driven by *Kcnc2*-Cre across cortical layers at different developmental stages (P14–P55). *Kcnc2*-Cre: n = 5 slices from 3 mice at P14; n = 4 slices from 4 mice at P20; n = 5 slices from 4 mice at P28; n = 6 slices from 3 mice at P35; n = 5 slices from 5 mice at P40; n = 6 slices from 6 mice at P55. **f,** FISH staining for the cardinal interneuron markers *Pvalb*, *Sst*, *Vip*, and *Npy* in *Kcnc2*-Cre:Ai9 mice at P45 across cortical layers. TdTomato (*Tdt*) signals predominantly overlap with *Pvalb* in L5/6, with minor overlap with *Sst*, *Vip*, and *Npy* in L2/3. n = 5 slices from 5 mice for *Pvalb*; n = 5 slices from 4 mice for *Sst*; n = 3 slices from 4 mice for *Vip*; n = 3 slices from 4 mice for *Npy*. **g,** Electrophysiological characterization of labeled neurons in *Kcnc2*-Cre:Ai9 mice by patch-clamp recording. Left, example patch-clamp image; middle, representative firing pattern; right, quantification. 45 recorded neurons in *Kcnc2*-Cre:Ai9 mice were identified as *Pvalb* interneurons based on their fast-spiking properties. **h,** Representative membrane potential traces from recorded L5/6 neurons in *Kcnc2*-Cre:Gq mice in response to clozapine-N-oxide (CNO, 10 µM; gray shading). Right, summary of firing rates before and after CNO administration. n = 6 cells from 4 *Kcnc2*-Cre:Gq mice; paired t-test, t(5) = 22.85, $^{***}p < 0.001$. **i,** Left: FISH staining for *Pvalb* in the thalamic reticular nucleus (TRN) of *Kcnc2*-Cre:Ai9 mice. Right: Quantification of the percentage of TdTomato-positive cells overlapping with *Pvalb* in the TRN. n = 4 slices from 3 mice. **j,** Left: EEG frequency band traces before (top, black) and after chemogenetic activation (bottom, blue) in *stg*:*Kcnc2*-Cre:hM3Dq mice, showing disappearance of the dominant 5-9 Hz absence seizure activity after CNO treatment. Right: Power spectrum analysis shows a marked reduction in 5-9 Hz power (black, baseline; red, shortly after activation; blue, 2 h after activation), consistent with seizure suppression (arrows), together with an increase in gamma-band power (35-55 Hz). **k,** Quantification of SWDs per hour in *stg*:*Kcnc2*-Cre:hM3Dq mice before and after CNO treatment. Data were analyzed using one-way repeated-measures ANOVA, followed by Dunnett's multiple comparisons test against baseline (Before). A significant effect of time was observed (F(1.015, 6.089) = 403.2, $^{**}p < 0.01$). Dunnett's test showed significant differences between Before and 1 h, 2 h, and 3 h (all $^{**}p < 0.01$). n = 7 mice. Vehicle treatment produced no significant change (n = 7 mice, paired t-test). CNO treatment in control *stg*:*Kcnc2*-Cre mice without hM3Dq expression also produced no significant change (n = 6 mice, Student's t-test). **l,** Quantification of mean SWD duration in *stg*:*Kcnc2*-Cre:hM3Dq mice before and after CNO treatment. Data were analyzed using one-way repeated-measures ANOVA, followed by Dunnett's multiple comparisons test against baseline (Before). A significant effect of time was observed ($F(1.025, 6.150) = 471.7$, $**p < 0.01$). Dunnett's test showed significant differences between Before and 1 h, 2 h, and 3 h (all $**p < 0.01$). n = 7 mice. Vehicle treatment produced no significant change (n = 7 mice, paired t-test). CNO treatment in control *stg*:*Kcnc2*-Cre mice without hM3Dq expression also produced no significant change (n = 6 mice, Student's t-test).

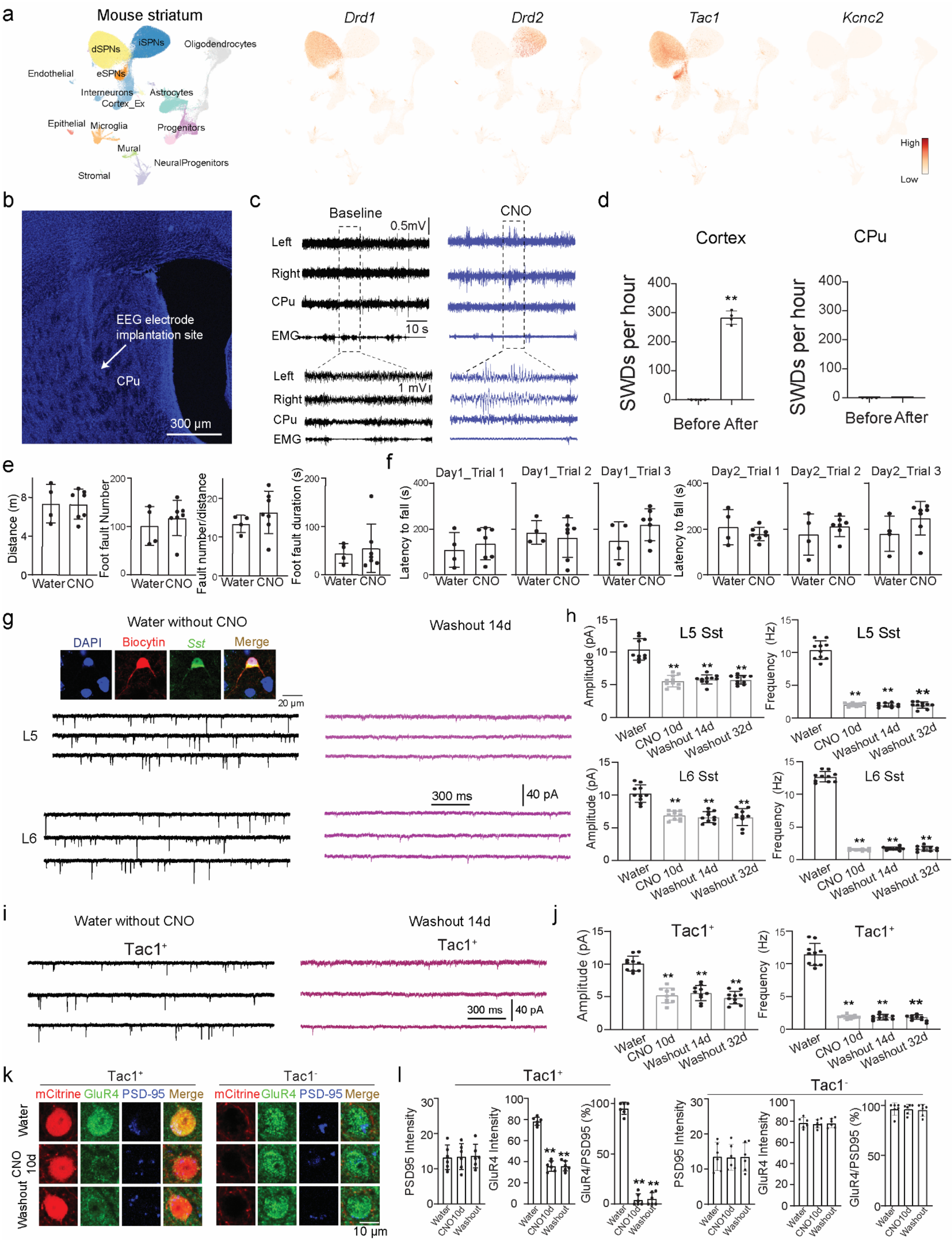

a
Mouse striatum
dSPNs
iSPNs
Oligodendrocytes
Endothelial
eSPNs
Interneurons
Cortex_Ex
Astrocytes
Epithelial
Microglia
Progenitors
Mural
NeuralProgenitors
Stromal
Drd1
Drd2
Tac1
Kcnc2
High
Low
b
EEG electrode implantation site
CPu
300 μm
c
Baseline
CNO
0.5mV
Left
Right
CPu
EMG
10 s
1 mV
d
Cortex
CPu
SWDs per hour
Before After
e
Distance (m)
Foot fault Number
Fault number/distance
Foot fault duration (s)
Water CNO
f
Day1_Trial 1
Day1_Trial 2
Day1_Trial 3
Day2_Trial 1
Day2_Trial 2
Day2_Trial 3
Latency to fall (s)
g
Water without CNO
Washout 14d
DAPI
Biocytin
Sst
Merge
20 μm
L5
L6
300 ms
40 pA
h
L5 Sst
L6 Sst
Amplitude (pA)
Frequency (Hz)
Water
CNO 10d
Washout 14d
Washout 32d
i
Water without CNO
Washout 14d
Tac1⁺
300 ms
40 pA
j
Tac1⁺
Amplitude (pA)
Frequency (Hz)
k
Tac1⁺
Tac1⁻
mCitrine
GluR4
PSD-95
Merge
Water
CNO 10d
Washout
10 μm
l
Tac1⁺
Tac1⁻
PSD95 Intensity
GluR4 Intensity
GluR4/PSD95 (%)
Water
CNO10d
Washout

**Extended Data Fig. 7 | Chronic Tac1$^{+}$ interneuron inhibition induces cortical but not striatal spike-wave activity and persistent circuit remodeling. Related to Fig. 5. a,** UMAP visualization of mouse CPu neurons adapted from Anderson et al. (2023), showing *Tac1* co-expression in the *Drd1* but not *Drd2* cluster. **b,** Representative image of the EEG electrode implantation site in the CPu core. **c,d,** EEG recordings show that 10-14 days of CNO treatment in *Tac1*-Cre:Gi mice induces seizure activity in the cortex but not in the CPu. Quantification in (d), paired t-test, t(3) = 25.18, $^{**}p < 0.01$. **e,f,** Foot-slip and rotarod tests show no significant motor deficits after CNO treatment. Water, n = 4; CNO, n = 7; unpaired Student's t-test. **g,h,** Representative traces and quantification of sEPSCs in L5 and L6 Sst interneurons. CNO treatment significantly reduced sEPSC amplitude and frequency, with persistent effects after washout. n = 10 cells per group. One-way ANOVA with Dunnett's multiple comparisons test versus Water: L5 amplitude, F(3, 36) = 49.49, $^{**}p < 0.01$; L5 frequency, F(3, 36) = 297.9, $^{**}p < 0.01$; L6 amplitude, F(3, 36) = 27.23, $^{**}p < 0.01$; L6 frequency, F(3,36) = 1412, $^{**}p < 0.01$; all Dunnett's comparisons vs Water, $^{**}p < 0.01$. **i,j,** Representative traces and quantification of sEPSCs in *Tac1*$^{+}$ interneurons. CNO treatment significantly reduced sEPSC amplitude and frequency, with persistent effects after washout. n = 10 cells per group. One-way ANOVA with Dunnett's multiple comparisons test versus Water: amplitude, F(3, 36) = 48.35, $^{**}p < 0.01$; frequency, F(3, 36) = 282.7, $^{**}p < 0.01$; all Dunnett's comparisons vs Water, $^{**}p < 0.01$. **k,l,** Representative images and quantification of synaptic GluR4 in *Tac1*-Cre:Gi mice. CNO treatment disrupted GluR4 synaptic trafficking in Tac1$^{+}$ neurons but not *Tac1*$^{-}$ neurons, without changing total PSD95 levels. Red, mCitrine; green, GluR4; blue, PSD95. Washout, 14 days. In Tac1$^{+}$ neurons, GluR4 intensity was reduced (one-way ANOVA, F(2, 15) = 173.6, $^{**}p < 0.01$) and GluR4/PSD95 colocalization was decreased (F(2, 15) = 448.4, $^{**}p < 0.01$), followed by Dunnett's multiple comparisons test versus Water (all $^{**}p < 0.01$). n = 6 cells per group. No significant changes were detected in Tac1$^{-}$ neurons.

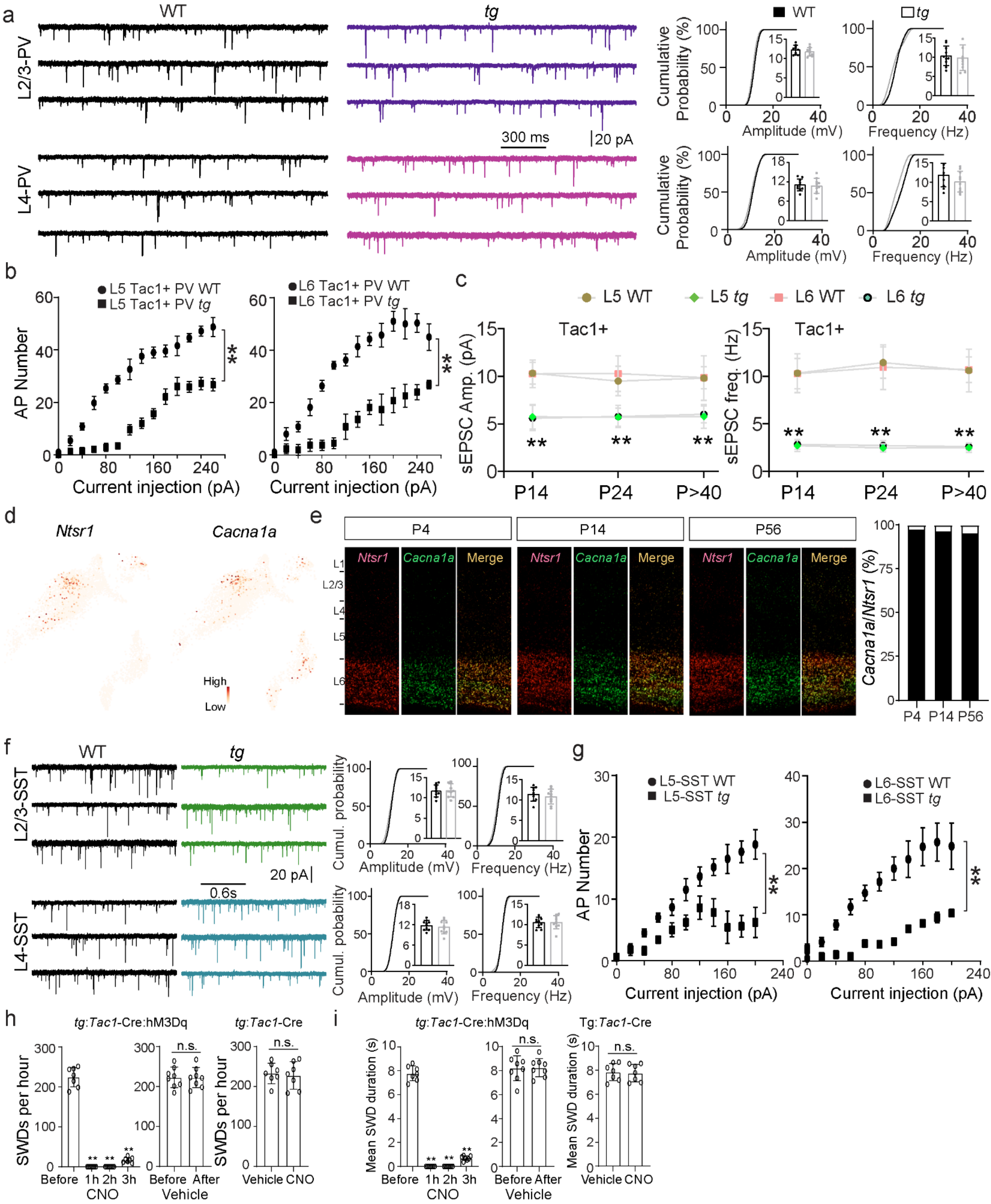
a
WT
tg
L2/3-PV
L4-PV
300 ms
20 pA
WT
tg
Cumulative Probability (%)
Amplitude (mV)
Frequency (Hz)
b
L5 Tac1+ PV WT
L5 Tac1+ PV tg
L6 Tac1+ PV WT
L6 Tac1+ PV tg
AP Number
Current injection (pA)
c
L5 WT
L5 tg
L6 WT
L6 tg
Tac1+
sEPSC Amp. (pA)
sEPSC freq. (Hz)
P14
P24
P>40
d
Ntsr1
Cacna1a
High
Low
e
P4
P14
P56
Ntsr1
Cacna1a
Merge
L1
L2/3
L4
L5
L6
Cacna1a/Ntsr1 (%)
f
WT
tg
L2/3-SST
L4-SST
20 pA
0.6s
Cumul. probability
Amplitude (mV)
Frequency (Hz)
g
L5-SST WT
L5-SST tg
L6-SST WT
L6-SST tg
AP Number
Current injection (pA)
h
tg:Tac1-Cre:hM3Dq
tg:Tac1-Cre
SWDs per hour
Before
1h
2h
3h
CNO
After
Vehicle
n.s.
i
tg:Tac1-Cre:hM3Dq
Tg:Tac1-Cre
Mean SWD duration (s)
Before
1h
2h
3h
CNO
After
Vehicle
n.s.

**Extended Data Fig. 8 | Characterization of cell type-specific synaptic and intrinsic excitability defects, and developmental *Cacna1a* expression in *tg*. Related to Fig. 6. a,** Representative traces of spontaneous excitatory postsynaptic currents (sEPSCs) recorded from L2/3 *Pvalb* (PV) interneurons and L4 PV interneurons in WT and *tg* mice. Cumulative probability plots and inset bar graphs show no significant differences in sEPSC amplitude or frequency between WT and Tg mice in either L2/3 or L4 PV interneurons. n = 10 cells for L2/3 PV WT, n = 10 cells for L2/3 PV *tg*; n = 10 cells for L4 PV WT, n = 10 cells for L4 PV *tg*. **b,** Firing responses, measured as the number of action potentials evoked by increasing current step injections, in L5 $Tac1^+$ interneurons (left) and L6 $Tac1^+$ interneurons (right) from WT and *tg* mice. In L5 $Tac1^+$ interneurons, WT: n = 5 cells, *tg*: n = 7 cells; two-way ANOVA, $F(1, 10) = 215.3$, $^{**}p < 0.01$. In L6 $Tac1^+$ interneurons, WT: n = 5 cells, *tg*: n = 5 cells; two-way ANOVA, $F(1, 8) = 74.62$, $^{**}p < 0.01$. **c,** Quantification of sEPSC amplitude and frequency in L5 and L6 $Tac1^+$ interneurons across developmental stages (P14, P24, and P > 40) in WT and *tg* mice. For L5 *Tac1*$^+$ interneurons: P14, n = 9 WT and n = 9 *tg*; P24, n = 10 WT and n = 8 *tg*; P > 40, n = 8 WT and n = 10 *tg*. For L6 *Tac1*$^+$ interneurons: P14, n = 7 WT and n = 7 *tg*; P24, n = 10 WT and n = 10 *tg*; P > 40, n = 8 WT and n = 9 *tg*. Student's t-test was used for WT versus *tg* comparisons at each developmental stage; significant differences are indicated by $^{**}p < 0.01$. **d,** UMAP visualization of cortical neuronal populations at P4 showing strong co-distribution of *Ntsr1* and *Cacna1a*. **e,** Double staining for *Ntsr1* (L6 corticothalamic marker, red) and *Cacna1a* (green) at P4, P14, and P56. Merged images are shown on the right. The proportion of *Ntsr1* neurons co-expressing *Cacna1a* was 97.8% at P4 (n = 6 mice), 96.7% at P14 (n = 7 mice), and 95.6% at P56 (n = 10 mice). **f,** Representative traces of sEPSCs recorded from L2/3 *Sst* interneurons and L4 *Sst* interneurons in WT and *tg* mice. Cumulative probability plots and inset bar graphs show no significant differences in sEPSC amplitude or frequency between WT and Tg mice in either L2/3 or L4 *Sst* interneurons. n = 10 cells for L2/3 *Sst* WT, n = 10 cells for L2/3 *Sst tg*; n = 10 cells for L4 *Sst* WT, n = 10 cells for L4 *Sst tg*. **g,** Firing responses, measured as the number of action potentials evoked by increasing current step injections, in L5 *Sst* interneurons (left) and L6 *Sst* interneurons (right) from WT and Tg mice. In L5 *Sst* interneurons, WT: n = 6 cells, *tg*: n = 5 cells; two-way ANOVA, $F(1,9) = 18.91$, $^{**}p < 0.01$. In L6 *Sst* interneurons, WT: n = 7 cells, *tg*: n = 5 cells; two-way ANOVA, $F(1,10) = 15.64$, $^{**}p < 0.01$. **h,** Quantification of spike-wave discharges (SWDs) per hour in *tg*:*Tac1*-Cre:hM3Dq mice before and after CNO treatment. Data were analyzed using one-way repeated-measures ANOVA followed by Dunnett's multiple comparisons test against baseline (Before). CNO treatment produced a significant time-dependent effect on SWD frequency ($F(1.123, 6.736) = 537.4$, $p < 0.01$). Post hoc Dunnett's comparisons revealed significant reductions at 1 h, 2 h, and 3 h compared with Before (all $^{**}p < 0.01$). n = 7 mice. Vehicle-treated mice showed no significant change in SWD frequency (n = 8 mice, paired t-test). In control *tg*:*Tac1*-Cre mice lacking hM3Dq expression, CNO treatment also did not significantly alter SWD frequency (n = 7 mice, Student's t-test). **i,** Quantification of mean SWD duration in *tg*:*Tac1*-Cre:hM3Dq mice before and after CNO treatment. Data were analyzed using one-way repeated-measures ANOVA followed by Dunnett's multiple comparisons test against baseline (Before). A significant effect of time was detected after CNO treatment ($F(1.086, 6.517) = 854.5$, $p < 0.01$). Dunnett's post hoc analysis showed that mean SWD duration was significantly different at 1 h, 2 h, and 3 h compared with Before (all $^{**}p < 0.01$). n = 7 mice. No significant change was observed after vehicle treatment (n = 8 mice, paired t-test). Similarly, CNO treatment did not significantly affect mean SWD duration in control *tg*:Tac1-Cre mice without hM3Dq expression (n = 7 mice, Student's t-test).
